# Fluorescent protein-based ticker tapes for multiplexed recordings of transcriptional histories in single cells in culture and in vivo

**DOI:** 10.1101/2025.09.08.675004

**Authors:** Ruizhao Wang, Jian Jiang, Zhuoyuan Li, Tianning Liu, Yangdong Wang, Mingqi Xie, Kiryl D. Piatkevich

**Affiliations:** College of Life Sciences, Zhejiang University, Hangzhou, Zhejiang, China; School of Life Sciences, Westlake University, Hangzhou, China; Westlake Laboratory of Life Sciences and Biomedicine, Hangzhou, China; Institute of Basic Medical Sciences, Westlake Institute for Advanced Study, Hangzhou, China; Key Laboratory of Growth Regulation and Translational Research of Zhejiang Province, School of Medicine and School of Life Sciences, Westlake University, Hangzhou, Zhejiang 310024, China; School of Life Sciences, Fudan University, Shanghai, China; School of Mathematical Sciences, Peking University, China; School of Engineering, Westlake University, Hangzhou, Zhejiang 310030, China

## Abstract

Recording and imaging of promoter activities in real time is critical for deciphering cellular states and dynamic signaling crosstalk, but technologies capable of simultaneously capturing multiple transient events in living cells are lacking. Here, we custom-design fluorescent protein-based ticker tapes (FPTT) for multiplexed and longitudinal recording of physiological activities in single cells. FPTT integrates multi-spectral monomeric fluorescent proteins with self-assembling protein fibers, enabling massively parallel analysis of signaling dynamics under varying cellular conditions. Using FPTT, we were able to log dose-dependent and reversible transcriptional histories of endogenous cFos signaling in primary hippocampus neurons at a 3-hour temporal resolution via biological timestamps and an extended recording time of over 8 days. Furthermore, we expanded the imaging toolset by engineering genetically encoded FPTT variants for human NFκB, JAK/STAT3, mTOR, NFAT and cAMP signaling, allowing for the quantification of potential crosstalk between cFos and NFκB pathways in neurons, multiplexed recording of STAT3– and cAMP-specific promoters during drug-induced liver injury in mice, as well as simultaneous analysis of up to four major signaling pathways involved in experimental T-cell activation. This platform advances single-cell analysis by providing a versatile tool to investigate transcriptional histories and signaling interplay across diverse biological contexts, with broad applications in developmental biology and disease modeling.

## Introduction

Studying cellular dynamics requires molecular tools capable of recording multiple physiological events simultaneously within individual cells. Such technologies are critical for elucidating the interplay between signaling pathways^1^, intracellular interactions^2^, developmental processes^3^, and disease mechanisms^4^. Advances in synthetic biology have produced DNA– and RNA-based molecular recorders for tracing gene expression histories of target promoters over extended periods of time^5–11^. However, these methods are often disruptive and require cell homogenization followed by sequencing readout, which could come at the expense of missing key spatiotemporal information and therefore prohibit real-time monitoring. On the other hand, genetically encoded fluorescence biosensors allow for high-resolution spatiotemporal reporting of physiological dynamics in real time^12–16^, but their utility is constrained by the limited spectral diversity of available biosensors, restricting possible combinations for simultaneous observations of cellular regions and events^17–19^. Furthermore, real-time imaging of the existing biosensors is also complicated by photobleaching and usually does not exceed an hour-long time scale^15,20,21^.

The recently introduced protein-based ticker tape systems showed remarkable potential for capturing transient^22^ or continuous physiological activities at a single-cell resolution^23,24^, providing a fundamentally new alternative to the existing nucleic acid-based recorders and fluorescent biosensors. For instance, the expression recording island (XRI)^23^, composed of a filament-forming *Escherichia coli* isoaspartyl dipeptidase 1POK(E239Y)^25^ fused to an epitope tag, enabled cFos activity recording in cultured mouse primary neurons and in vivo brain tissue. An alternative system utilized a fusion protein combining the catalytic domain of Pak4 kinase and the iBox domain of its inhibitor Inka1^26^ (referred to as iPAK4), coupled with a fluorescent probe, to report cFos-driven activity in cultured mammalian neurons^24^. These tools encode transcriptional activation events into elongating protein chains, potentially enabling multiplexed recordings within single cells. Bidirectional filament growth permits the insertion of specified protein tags at both ends upon activity-specific triggers, with temporal event sequences deciphered post hoc via immunostaining or fluorescent dyes along the formed protein filaments. However, XRI exhibits limited cell type compatibility, suboptimal temporal resolution, and reliance on post-fixation antibody-based readouts. Although the iPAK4 approach offered improved temporal resolution and broader cell-type compatibility compared to XRI, it was reliant on administration of exogenous dyes and therefore also complicated *in vivo* applications. Furthermore, while both systems have primarily focused on recording single cFos activity, their capacity for the simultaneous recording of multiple signaling pathways remains largely unexplored.

Here, we report a fluorescent protein-based ticker tape (FPTT) that overcomes these limitations by allowing visualization and quantification of multiple promoter activities at a single-cell resolution and using simple endpoint analysis under a conventional fluorescence microscope. Mechanistically, FPTT integrates spectrally distinct monomeric fluorescent proteins fused with iPAK4 and captures single activation events of small drug inducible and synthetic physiologically relevant promoters into a large-scale multiplexed “recording transcript” under fluctuating cellular conditions in single mammalian cells **(Fig. 1A)**. FPTT not only allows dose-dependent and reversible recording of cFos-specific promoters in primary hippocampus neurons, but also achieves a balance between temporal resolution and duration of recordings by enabling the incorporation of cumate-inducible biological timestamps at 3-hour intervals over up to 8 consecutive days. Furthermore, we show that FPTT was also compatible with other signaling-specific promoters engineered to contain response elements for human NFκB, STAT3, mTOR, NFAT and cAMP signaling, respectively. Using such an expanded FPTT toolkit, we were able to quantify potential delayed response between cFos and NFκB crosstalk in neurons, a putative orthogonality between STAT3 and cAMP signaling in human cell lines, a possible crosstalk between STAT3 and NFAT during KCl treatment, as well as oscillating dynamics of mTOR signaling in cell culture. Capitalizing on various multi-channel recording strategies optimized for expression in Sleeping Beauty-based transposons, we were able to use FPTT to study multiple promoter activities during experimental T-cell activation *in vitro* as well as drug-induced liver injury in mice in terms of sequential order and potential interdependence of activation. Taken together, FPTT broadens the utility of protein-based recording and has the potential to inaugurate a transformative platform for dissecting complex biological processes for mechanistic studies.

**Fig. 1.**
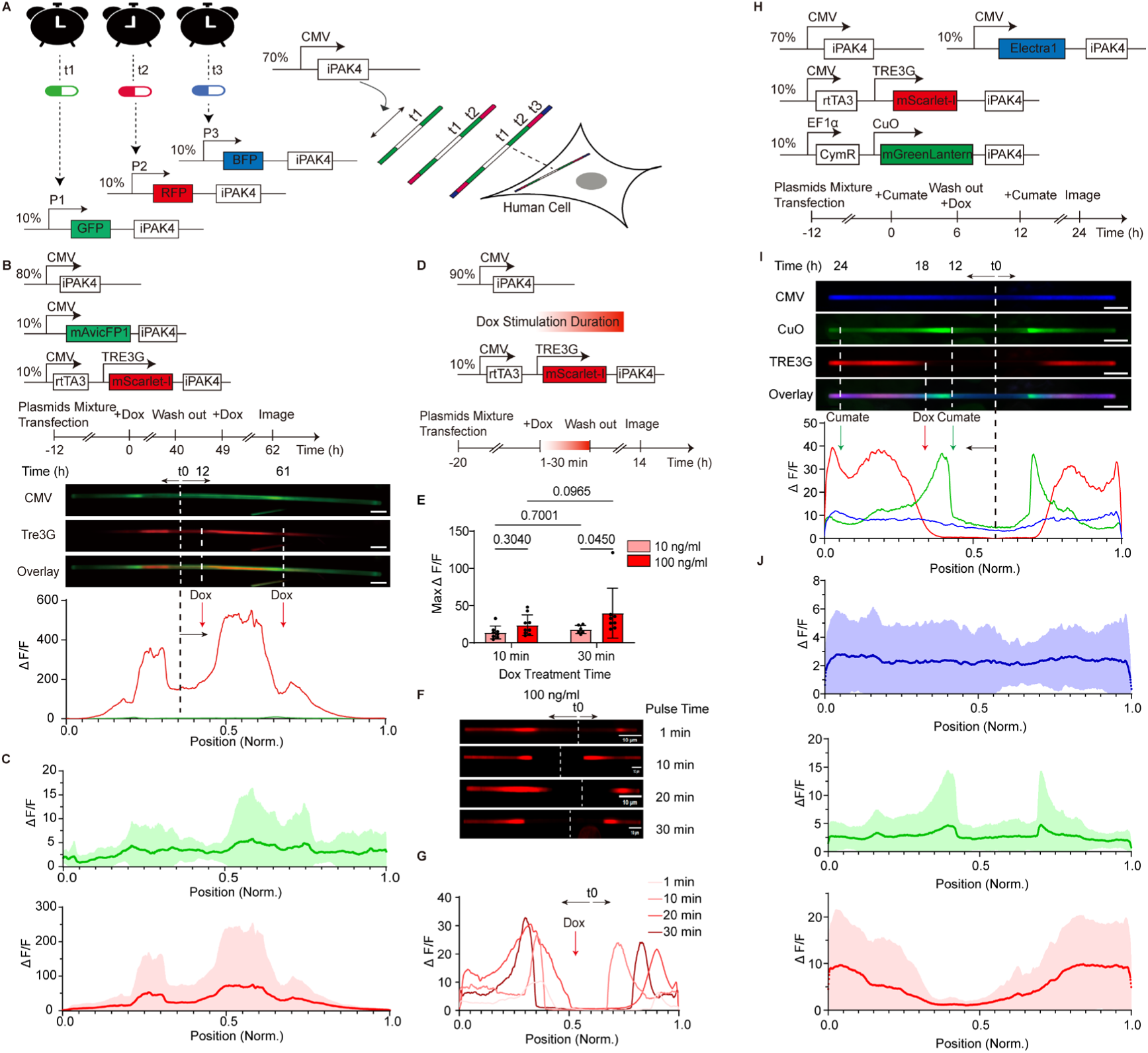
Time-dependent, reversible, and multichannel fluorescent recording in cultured mammalian cells. **A**, Scheme for intracellular recording of multiple physiological activities with fluorescent protein-tagged ticker tape. iPAK4, expressed under control of constitutive promoter, forms the continuously growing fiber scaffold. FPs-iPAK4 expression driven by diverse kinds of physiologically relevant promoters (P1, P2, P3, etc.), introducing color bands into the fiber at separate times points (t1, t2, t3, etc.) corresponding to promoter activation chronology. **B**, Schematic of the expression cassettes and experimental protocol used for Dox-induced transcriptional activity recordings in HEK293T cells. Transcription was activated via addition of Dox at t = 0 h and 49h respectively, and images were captured at t = 62h. Representative images and corresponding fluorescence intensity changes (ΔF/F) profile of the fiber was shown. The red line indicates Dox-inducible TRE3G expression, and the green line indicates constitutive CMV activity. t0 and black dash line, the start timepoints of fiber growth; black arrow, the direction of fiber growth; red arrow, the application of Dox. Scale bars, 10 μm. **C**, Averaged ΔF/F profiles of two times induction (n = 9 fibers from three independent cultures). The red line indicates Dox-inducible TRE3G expression, and the green line indicates constitutive CMV activity. Solid line, mean; shaded area, SD. **D**, Schematic of the expression cassettes used for Dox-induced pulse transcriptional activity recordings in HEK293T cells and experimental protocol for recording a short pulse of Dox-induced transcription activities. Transcription was activated via addition of two concentrations of Dox (10 ng/ml or 100 ng/ml) for 1, 10, 20, and 30 minutes, and images were captured at 14 hours post treatment. **E**, The maximum fold changes (ΔF/F) of each fiber as treated in conditions illustrated in D. Each black dot indicated one maximum ΔF/F of one fiber. Data are shown as the mean±SD of n=3 independent experiments, *p* values are calculated using two-way ANOVA. **F**, Representative images of fibers as treated with 100 ng/ml Dox with 1, 10, 20, and 30 minutes. t0 and white lines, the start timepoint of each fiber growth; black arrow, the direction of fiber growth. Scale bars, 10 μm. **G**, Corresponding ΔF/F profile of the fiber shown in F. t0, the start timepoint of each fiber growth; black arrow, the direction of fiber growth; red arrow, the application of Dox. **H**, Schematic of the expression cassettes and experimental protocol used for multiplexed transcriptional activity recordings in HEK293T cells. Cumate controllable activity was activated via addition of 1× Cumate water solution at t = 0, 12 h, respectively. Then the Tet-On system was activated via addition of Dox at t = 6 hours. **I**, Representative images and corresponding ΔF/F profile of a fiber formed under conditions illustrated in H. t0 and black dash line, the start timepoints of fiber growth; black arrow, the direction of fiber growth; red arrow, the application of Dox; green arrow, the application of Cumate. Scale bars, 10 μm. **J**, Averaged ΔF/F profiles (n = 13 fibers from three independent cultures). The blue line indicates constitutive CMV activity, the red line indicates Dox-inducible TRE3G expression, and the green line indicates Cumate induced CuO activity. Solid line, mean; shaded area, SD.

## Results

### Selection and optimization of protein filaments to form single fluorescent fibers per cell

To validate the feasibility of generating spectrally multiplexed ticker tapes that form single fiber per cell, we first assessed the subcellular localization of 1POK(E239Y) and iPAK4 fusions with spectrally diverse fluorescence proteins (FPs) by transient expression in HEK293T and HeLa cells. Although 1POK(E239Y) was shown to form single fibers when expressed with small epitope tags in cultured neurons in previous studies^23^, it formed irregularly shaped fibers or small aggregates with multiple short fibers in HEK293T and HeLa cells, depending on the fused FP (**Fig. S1**). Similarly, the expression of monomeric mEGFP-tagged iPAK4 only produced relatively short and truncated aggregates instead of single fibers (**Fig. S2**). In contrast, iPAK4 consistently formed well-defined rod-shaped structures with various monomeric FPs when an untagged iPAK4 fraction was expressed in abundance (**Fig. S3**). Importantly, most cells formed a single fiber per cell, though some exhibited more than two fibers or aggregates, likely due to uneven expression levels in the co-transfection system—an observation consistent with previous findings using HaloTag-iPAK4^24^. Thus, the monomeric nature of the selected FP appeared to be critical for fiber formation, since fusion of the dimeric StayGold^27^ with iPAK4 under same experimental setups again failed to form rod-shaped structures in HEK293T cells (**Fig. S4**). Adjusting the expression ratio of 1POK(E239Y) with FP-1POK to 9:1, by analogy with the iPAK4 system, also failed to yield single filaments (**Fig. S5**).

We observed that the formed iPAK4 fibers were significantly longer than the cell dimensions. To confirm their intracellular localization, we labeled plasma membrane by co-expression of mCherry-CAAX and nucleus by staining with Hoechst 33342. Our findings indicated that even fibers approximately 100 μm in length remained intracellularly incorporated, although a slight membrane deformation was observed, consistent with previous studies^24^ (**Fig. S6**). To assess the physiological impact of this deformation, we conducted independent experiments assessing cell viability and proliferation using several approaches.

We confirmed the formation of multiple fibers both before and after trypsin digestion; however, iPAK4 expression did not statistically impact cell survival as determined using Trypan Blue staining (**Fig. S7**). Furthermore, a CCK-8 assay, used to assess both cell viability and proliferation, also revealed no significant negative effect on cell proliferation, although a minor effect was detected 72 hours post-plating (**Fig. S8**). Quantification of HEK293T cells with or without transfected iPAK4 fibers using a live/dead assay via flow cytometry, expressing iPAK4 showed no statistically significant effect on early apoptosis (Annexin V^+^ PI^-^), late apoptosis and necrosis (Annexin V^+^ PI^+^) and overall cell death (Annexin V^-^ PI^-^) (**Fig. S9**). We also observed FPTT-expressing cells containing two nuclei (**Fig. S10**), suggesting that the presence of iPAK4 fibers might not obstruct cell division further confirming previous study reporting that iPAK4 expression did not impede mitosis^24^.

To further characterize the growth kinetics, we quantified the growth of multiple fibers using live imaging in HEK293T cells, starting 6 hours post-transfection and continuing for 48 hours (**Supplementary movie 1**). We observed that fiber formation initiates centrally following nucleation, with assembly proceeding from the middle towards the ends. While prior research described a nearly linear growth phase post-nucleation lasting approximately 20 hours, characterized by asymmetrical growth with distinct fast and slow-growing ends^24^, our study also identified an initial rapid growth phase within the first hour post-nucleation, succeeded by a linear growth phase. However, this linear phase exhibited substantial cell-to-cell variability in growth rates, leading to diverse durations of linear growth, sometimes uneven growth, and differences in final fiber lengths (**Fig. S11; Supplementary Movie 1**). Following the linear growth phase, a steady growth phase was observed. Remarkably, in some exceptional instances, fibers exceeded 150 μm in length, with growth continuing for another 20 hours starting from 20 hours post-transfection, thereby demonstrating the potential for prolonged real-time recordings (**Fig. S11**).

To further optimize the FPTT system, we explored different expression strategies including varying ratios of iPAK4 and FP-iPAK4 fusion by co-transfecting two separate plasmids as well as testing a bicistronic plasmid containing an IRES element (**Fig. S12, S13**). While the range of co-transfection ratios from 9:1 to 5:1 showed consistent fiber formation, the single bicistronic plasmid with IRES failed to produce fibers during short-term expression periods over 15 hours, potentially due to insufficient iPAK4 monomer concentration available for nucleation (**Fig. S12**). Although a 5:1 (w/w) ratio was adequate for intracellular fiber formation, it occasionally led to cellular aggregates (**Fig. S13**), whereas the stricter 9:1 (w/w) ratio provided more consistent results at short-term and long-term expression time points. As a result, in subsequent experimental setups, we opted to use the 9:1(w/w) co-transfection ratio of iPAK4: mFP-iPAK4 plasmids to demonstrate the system’s applicability, which we termed Fluorescent Protein-based Ticker Tape (FPTT).

Next, to verify possibility for multiplexed imaging, we co-transfected the spectrally diverse mXFP-tagged iPAK4 variants together with iPAK4 in 1:9 ratio under the control of a constitutive promoter. All these FPs colocalized on a single fiber, indicating their compatibility for simultaneous fusion within the rods (**Fig. S14**). Taking together, our results show that iPAK4 forms linear fibers with monomeric FPs but not with dimeric FPs when co-transfected in the ratio of iPAK4: mFP-iPAK4 9:1 (w/w) in HEK293T cells, whereas 1POK fails with any combination with FPs. Thus, we focused on the FPTT as the engineering basis when moving towards the goal of real-time monitoring and endpoint readout of multiple transcriptional activities in single cells.

### Time-dependent, reversible, and multichannel fluorescent recording in cultured mammalian cells

To study the temporal dynamics of recording instant promoter activities using FP-based ticker tapes, we used the well-established Tet-ON system^28^ to allow time-controlled activation by a small molecule trigger doxycycline (Dox, 10 ng/ml) at specific time points in HEK293T and HeLa cells. In both cell types, we co-expressed a green tape driven by a constitutive CMV promoter with a TRE3G-controlled red tape or vice versa. End-point imaging was conducted following either a single (**Fig. S15**) or two consecutive pulse inductions (**Fig. 1B, Fig. S15**) or without Dox treatment **(Fig. S16**). To quantify promoter activation, green and red fluorescence intensity fold change profiles (ΔF/F) along each fiber were calculated and plotted against fiber length. The obtained profiles indeed exhibited two asymmetrical peaks that corresponded to time-controlled promoter activation on a fast-growing end and a slower-growing end (**Fig. 1B**). Furthermore, ΔF/F profiles were also calculated to allow quantitative comparison of relative activation strengths^12,29,30^. Although the fiber growth and corresponding intensity profiles exhibited asymmetry with distinct fast– and slow-growing ends, the recorded transcriptional events demonstrated a symmetrical distribution from the central nucleation point outwards. This inherent symmetry enabled us to determine the temporal order of transcriptional events by analyzing growth solely in one direction, from the fiber’s midpoint (t0) to its distal end. Consequently, single and averaged ΔF/F profiles were analyzed unidirectionally from the middle to the end, comparing them to a control where the Tre3G promoter showed minimal or no activated pattern (**Fig. 1B-C, Fig. S16**). We confirmed that timepoint-controlled transcriptional activity was indeed marked with a distinct pattern behind the constitutive expression fluorescence signals after specific time intervals (**Fig. 1B-C**, **Fig. S15**). These results indicate that the FPTT system could record temporal information over multiple days while allowing relative quantitative comparison of inducible promoter activation strengths.

Next, we used different transient pulses (1 to 30 minutes) of Dox stimulation (10 ng/ml or 100 ng/ml) to study the sensitivity of FPTT recording (**Fig. 1D**). Quantification of peak ΔF/F revealed a correlation of fluorescence amplitude only slightly with a duration of Dox pulse but highly related to dose concentration (**Fig. 1E**). Imaging the cells at 34 h post-transfection revealed fibers with distinct fluorescence patterns against fiber length (**Fig. 1F**). We observed well-defined peaks even during minutes-scale stimulation, corresponding to reversible promoter activation (**Fig. 1G**). We also observed a statistically significant effect, indicating that the Tre3G promoter achieved full activation with 100 ng/ml of the inducer Dox within 30 minutes (**Fig. S17**). These results suggest that the ticker tape system is sensitive enough for recording even brief and reversible promoter’s activation on a daily time scale, thus allowing relative quantification of expression levels.

To determine if the time order of activation could also be accurately recorded for multiplexed transcriptional activities, we used two different small molecule-inducible synthetic promoters to co-express two spectrally distinct iPAK4-fused mFPs in parallel. A cumate-inducible^31^ green ticker tape was activated at a 12-hour timescale to create a timestamp for Tet-On activation. Dox was added to the medium between two cumate timestamps, while a blue fluorescent ticker tape was constitutively expressed under the CMV promoter (**Fig. 1H, I**). By measuring the fluorescence profile, we observed that the Dox-activated red fluorescent fiber was clearly marked between two green fluorescence peaks induced by cumate (**Fig. 1I, J**). This confirms that the FPTT technology is amenable to the recording of at least three transcriptional activities in a time-ordered manner.

To showcase a physiological scenario where such a tool for tracking cellular activities could be essential for continuous recordings of promoter activities, we evaluated our FPTT system in cultured primary neurons. First, we placed a green mFP-tagged iPAK4 under the control of the cFos promoter^32^ for dose-dependent recording of externally supplied Phorbol 12-myristate 13-acetate (PMA), a known activator of protein kinase C (PKC) and cFos signaling^24,33^. Primary mouse hippocampus neurons were co-transfected with a P_hSyn_-driven iPAK4 expression vector (90%, w/w) and a P_cFOS_-driven mAvicFP1-iPAK4 expression vector (10%, w/w), followed by PMA administration after 24 h. Six hours after PMA induction, the neurons were imaged, and intensity profiles of the formed fibers were analyzed (**Fig. S18 A**). Although cFos promoter exhibited a basal level of expression without PMA^34,35^, addition of PMA induced a clear green fluorescent band at the end of the fibers (**Fig. S18 B**). By analyzing the averaged ΔF/F profiles of each fiber in each concentration, we found that the fluorescent signal from the ticker tape exhibited a dose-dependent increase in response to PMA concentrations. The dynamic range of ΔF/F spanned from a 2-fold to an 8-fold increase in response to the PMA concentrations ranging from 10 ng/ml to 500 ng/ml (**Fig. S18 B-S8D**). Although neurons formed significantly longer fibers (in some cases, exceeding 200 μm) compared to HEK cells, a previous study reported no adverse effects on cell viability or physiological parameters (*e.g*., action potentials, membrane potential, input resistance, and membrane capacitance).

To further test the capability for long-term recordings, transfected cells were treated with 50 ng/ml PMA at 48 hours post-transfection, with an interval of 12 hours on the first two days, 20 hours, and 30 hours for another two cycles, respectively (**Fig. 2A, B**). Continuous monitoring of a single neuronal fiber revealed at least four distinct cFos activation events within the above-mentioned time span of up to four days *in vitro*, although an uneven growth was also detected during this timescale, similar to that seen in HEK293T cells and previous findings (**Fig. 2B; Fig. S19**). We also induced transcriptional activity by applying cumate for 18 hours every 24 hours, repeating this process multiple times until reaching day 8 post-transfection (**Fig. 2C, D**). We successfully recorded at least eight activations in eight days from the initial treatment (**Fig. 2D**), demonstrating the potential of our fluorescent ticker tape for long-term recordings of cellular events. To our knowledge, this extended duration, with a one-day resolution, surpasses the capabilities of existing protein recording tools ever reported^23,24^, which has been a long-sought goal of neuronal imaging.

**Fig. 2.**
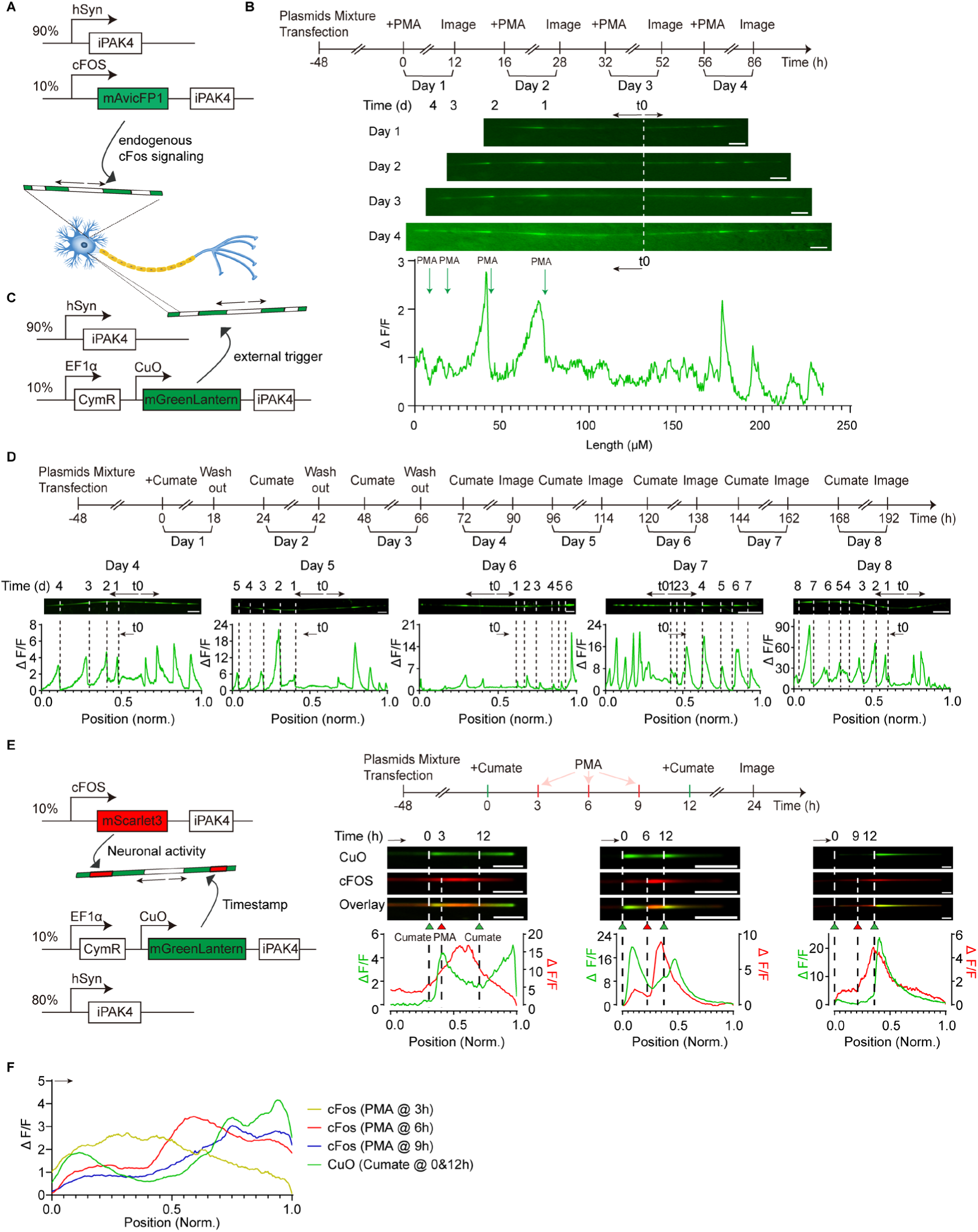
Reversible, long-term recording of neuronal activity over multiple days and its time resolutions with biological timestamp. **A**, The expression cassette used for PMA-induced cFOS transcriptional activity recordings in mouse primary hippocampus neurons over days. **B**, Experimental protocol for recording PMA-induced cFOS transcription activities over days. Cultured neurons with transfection described in A were stimulated with 50 ng/ml PMA every 16 hours for four consecutive days while daily microscopic analysis of green fluorescence signals indicative for cFOS activation was performed. Representative images and corresponding ΔF/F profiles of same fiber via daily activation by PMA as indicated in B. t0 and white dash line, the start timepoints of fiber growth; black arrow, the direction of fiber growth; green arrow, the application of PMA. Scale bars, 10 μm. **C**, The expression cassette used for Cumate-induced transcriptional activity recordings in mouse primary hippocampus neurons over days. **D**, Transfected cells were stimulated with 1X aqueous Cumate solution over 18 hours each day. Microscopic analysis with green fluorescence signals (ΔF/F) indicative for Cumate stimulation was performed on days 4-8. Representative images and ΔF/F profiles of the fiber from day 4 to day8. Each black dash lines on fibers indicate one induction of CuO activity. t0, the start timepoints of fiber growth; black arrow, the direction of fiber growth. Scale bars, 10 μm. Day 4, N = 22 fibers; Day 5, N = 20 fibers; Day 6, N = 30 fibers; Day 7, N = 23 fibers; Day 8, N = 38 fibers from three biological replicates. **E**, The expression cassette and experimental protocol used for multiplexed transcriptional activity recordings in mouse primary hippocampus neurons. Transfected neurons were exposed to successive pulses of Cumate at 0h and 12h, 50ng/ml PMA was added to the medium at 3h, 6h, or 9h from the first-time addition of 1X Cumate water solution. Images were acquired at 12h after the last stimulation event. Representative images and corresponding ΔF/F profiles of the fiber treated with PMA at 3h, 6h, or 9h behind the first-time stimulation of CuO activity. Dash lines with green triangle represent one time induction of CuO activity, and red tangle indicate the activation of cFOS activity. Black arrow, the direction of fiber growth. Scale bars, 10 μm. **F**, The averaged ΔF/F profiles of multiplexed activities. Dash lines with green triangle represent two-time induction of CuO activity at 0 and 12h, and red tangle indicate the PMA activated cFOS at 3h, 6h, and 9h behind the first activation of CuO activity. Black arrow, the direction of fiber growth. PMA addition at t = 3 h, N = 5 fibers; t = 6 h, N = 12 fibers; t = 9 h, N = 9 fibers from two biological replicates.

The ability to record physiological activities in neurons over an extended period of up to eight days (**Fig. 2D**) creates a need for timestamps as reference points for aligning the recorded activity with user-defined time points. To address this, we used the cumate system to create time stamps on the fiber by adding cumate to the cells at defined time points (**Fig. 2E**). Correspondingly, we co-transfected neurons with the cFos/cumate FPTT system and activated the cumate promoter at 12-hour intervals. Within each 12-hour interval between cumate activations, we activated the cFos promoter by adding PMA at three different time points: 3, 6, and 9 hours after the first cumate induction (**Fig. 2E**). Endpoint analysis of fibers was conducted 24 hours after the initial cumate induction. The intensity profiles of the formed fibers exhibited two distinct peaks of cumate activation and one peak of cFos activation in response to PMA (**Fig. 2E**). Importantly, cFos FPTT peak was positioned between cumate peaks with gradual shift from earlier to later cumate peak for 3 tested time points (**Fig. 2F**). This experiment demonstrated the ability to timestamp PMA-induced cFos activity with a resolution of at least three hours.

### Systematic analysis of potential crosstalk and interdependence between different pairs of signaling pathways

Signaling pathways play a pivotal role in the regulation of various biological processes and conditions, including development, inflammation, and central metabolism. For example, IL-6-induced STAT3 activity is upregulated in the inflammation process^36^. NFAT proteins are regulated by Ca^2+^ and the Ca^2+^/calmodulin-dependent serine phosphatase calcineurin and activated as a result of calcium flux to regulate downstream related gene expressions^37–39^. In addition, the transcription factor NF-κB regulates multiple aspects of innate and adaptive immune functions and serves as a pivotal mediator of inflammatory responses^40,41^. cAMP, as the first identified second messenger, converts and amplifies extracellular signals by activating protein kinase A, which subsequently phosphorylates the cAMP response-element binding-protein (CREB) to activate and regulate gene transcription^42,43^. The intricate interplay of these signaling pathways orchestrates complex cellular behaviors, allowing cells to respond dynamically to internal and external stimuli. However, the temporal correlations of these pathways’ activation in response to a biologically meaningful stimulus are poorly studied due to the lack of the appropriate technologies.

Having demonstrated the ability of FPTT for long-term, multiplexed recordings of physiological and pharmacological activities in various mammalian cell types, we next aimed for multiple endogenous signaling readouts by implementing physiologically relevant promoters. To investigate the capability of recording different signaling pathways in biologically relevant cellular activities, we selected a set of validated synthetic promoters engineered to contain response elements for endogenous STAT3, NFAT, NFκB and cAMP-signaling^44–47^, respectively, and cloned various FP-iPAK4 constructs under each promoter for validation in HEK293T cells (**Fig. S20 A-D**). In the case of the PKA/cAMP-dependent system, a vanillic acid-responsive GPCR coupling to the Gαs-PKA axis (MOR9-1) was co-transfected or stably integrated into HEK293T cells^47^ (**Fig. S20 D**). Transfected cells were stimulated by adding IL-6 (500 ng/ml), KCl (45 mM), hTNF-α (0.5 ng/ml), and vanillic acid (VA, 500 μM) to induce STAT3, NFAT, NFκB, and PKA/cAMP signaling pathways, correspondingly. As a control of the basal activity, we used transfected cells in the same experimental setting without any exogenous stimulation. Results show that a clear fluorescent band was observed at the end of all fibers, indicating that the specific stimulations indeed activated the corresponding signaling pathways. Averaged ΔF/F profiles under each condition revealed variability in the basal expression levels of certain signaling pathways and distinct fold changes upon induction. For example, STAT3 and NFκB exhibited higher basal expression levels compared to other pathways, while cAMP demonstrated more pronounced activation maxima (**Fig. S20 A, C, D**). Notably, under control conditions, *i.e*., no stimulation groups, the NFAT-reported ticker tape showed an exceptionally low basal expression level, which limited the number of fibers obtained in this as well as in subsequent experiments (**Fig. S20 B**).

#### Visualization of potential crosstalk between cFos and NFkB signaling in neurons

The coordinated activities between NFκB and cFos are crucial in neuronal signaling, particularly in response to various stimuli^48–50^. To investigate the temporal relationship between these two transcription factors, we first tested the responsiveness of NFκB and cFos ticker tapes to each other’s known agonists, such as PMA and TNF-α^51^. 48h after co-transfection of primary murine hippocampus neurons with a P_hSyn_-driven iPAK4 expression vector (90%, w/w) and a P_NFκB_-driven mChilada-iPAK4 expression vector (10%, w/w), cells were stimulated with 200 ng/ml mTNF-α before microscopic images were acquired. A noticeable increase in signal intensity was observed as well as a clear band emerged on the end of the fiber compared with the control without any stimulations (**Fig. S21**), although NFκB showed substantial basal expression as indicated in the middle of the fiber before induction happens, possibly due to its crucial role in neuron development^52^. We also tested the effect of both PMA and TNF-α on cFos and NFκB independently. Primary mouse hippocampus neurons were co-transfected with a P_hSyn_-driven iPAK4 expression vector (90%, w/w) and a P_NFkB_-driven mScarlet3-iPAK4 expression vector or a P_cFOS_-driven mGreenLantern-iPAK4 expression vector (10%, w/w) before cells were stimulated with 50 ng/ml PMA or 200 ng/ml mTNF-α on two consecutive days (**Fig. S22**). Our findings demonstrated that TNF-α and PMA activated both cFos and NFκB transcription (**Fig. S18**, **Fig. S21**, **Fig. S22**).

The same results were also obtained when neurons were co-transfected with cFos-driven mGreenLantern-expressing and NFκB-driven mScarlet3-expressing ticker tapes, followed by sequential 24 h-treatments with TNF-α (200 ng/ml) and PMA (500 ng/ml), respectively (**Fig. 3A, B**). In this time frame, without TNF-α and PMA treatment, no obvious activated patterns were observed on the fibers and profiles, although a higher background of NFκB was found around the start timepoints, similar as independent experiment in NFκB treatment without PMA (**Fig. S23**). However, in a sequential treatment with TNF-α and PMA, although TNF-α and PMA activated both cFos and NFκB transcription, the resulting fiber intervals suggest that cFos activity could precede NFκB, as evidenced by its closer proximity to the fiber’s center (**Fig. 3C; Fig. S24**). To rule out potential biases due to selected fluorescent protein, we swapped the promoter controlling the FP-iPAK4 cassette, confirming that cFos consistently responded to stimuli prior to NFκB – no matter whether green or red monomeric FP variants were used (**Fig. 3D, E; Fig. S24,)**. Statistically significant activation of cFos and NFκB were also observed compared to the control (**Fig. S23 D, H**). These results confirm the capability of the FPTT technology to accurately report the relative timing of activation events for multiple cellular events in parallel in response to specific stimuli.

**Fig. 3:**
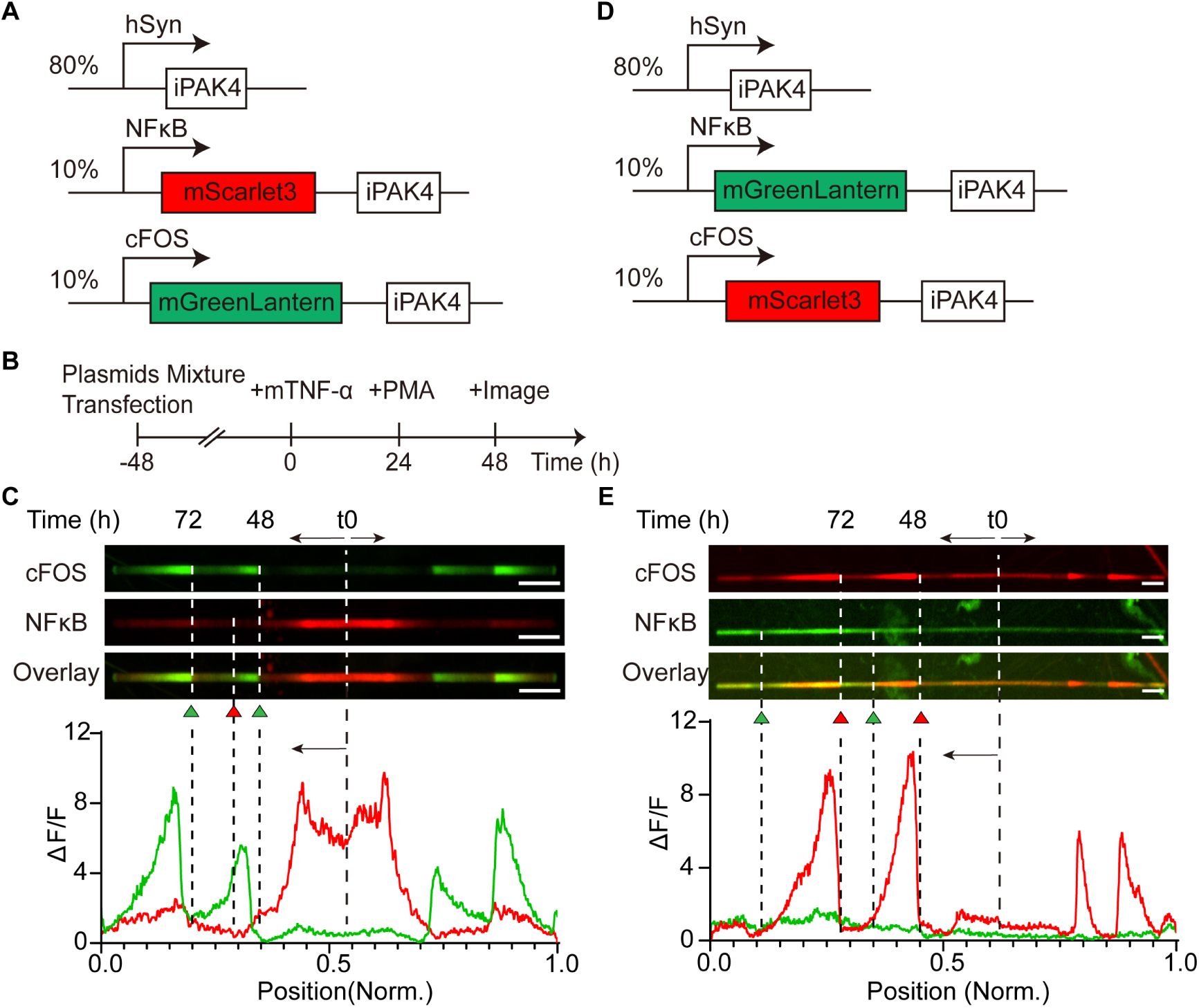
Visualization of potential crosstalk between cFos and NFkB signaling in neurons. **A**, The expression cassette used for recording the crosstalk between cFOS and NFκB in mouse primary hippocampus neurons. **B**, Experimental protocol for recording the crosstalk between cFOS and NFκB signaling. Transfected neurons were exposed to stimulation with 200 ng/ml mTNF-α for one day followed by another day of cultivation in cell culture medium containing 50 ng/mL PMA before images were acquired after another 24h. **C**, Representative image and corresponding ΔF/F profiles of the fiber treated in B under A constructs. Dash lines with green triangle represent cFOS activation, and red tangle indicate NFκB induction. Red solid line, NFκB activity; green solid line, cFOS activity. t0, the start timepoint of each fiber growth; black arrow, the direction of fiber growth. Scale bars, 10 μm. **D**, The expression cassette used for recording the crosstalk between cFOS and NFκB in mouse primary hippocampus neurons, with reversible fluorescent proteins as used in A. **E**, Representative image and corresponding ΔF/F profiles of the fiber treated in B under D constructs. Dash lines with red triangle represent cFOS activation, and green tangle indicate NFκB induction. Red solid line, cFOS activity; green solid line, NFκB activity. t0, the start timepoint of each fiber growth; black arrow, the direction of fiber growth. Scale bars, 10 μm.

#### Quantification of orthogonal cAMP and STAT3 signaling in mammalian cells and in a mouse model of liver injury

While the crosstalk between NFκB and cFos pathways could have been anticipated for neurons^53^, information on potential interdependence between cAMP and STAT3 signaling is less convergent. While some reports reveal a correlation of these pathways during inflammation and liver diseases^54^, recent research findings speculate that the signaling cascades for each other’s receptor-to-transcription coupling may be mutually orthogonal from a qualitative point of view^47^. To address this question, we expressed a P_CREB_-driven mGreenlantern-based FPTT system into native or MOR9-1-transgenic HEK-293T cells, followed by 9 hours of treatment with IL-6 (a STAT3 agonist) or vanillic acid (a MOR9-1/PKA agonist). Results show that IL-6 had no significant effects on cAMP expression, whereas vanillic acid effectively activated cAMP signaling either through transiently or stably expressed MOR9-1 (**Fig. S25**). Similarly, a P_STAT3_-driven mScarlet3-based FPTT was expressed in these cell types, confirming that only IL-6, but not vanillic acid, could activate STAT3-specific promoters (**Fig. S26**). The same results were also obtained when both a P_CREB_-driven mGreenlantern-based FPTT and a P_STAT3_-driven mScarlet3-based construct were co-expressed into same MOR9-1-expressing cell configurations (**Fig. S27**).

Thus, capitalizing on this putative orthogonality between STAT3 and cAMP signaling, in which VA exclusively activated cAMP and IL-6 exclusively activated STAT3-specific promoters, we used the FPTT technology to record the activation dynamics of these two pathways in mice **(Fig. 4)**. In fact, STAT3 activation is known to facilitate liver regeneration and offers protection against oxidative damage during acute liver injury, whereas immunosuppressive cAMP signaling modulates metabolic flux to satisfy the energy demands of regeneration^55–57^. However, severe liver damage or liver diseases, such as fibrosis and non-alcoholic fatty liver disease, can lead to aberrant STAT3 activity and impaired cAMP signaling^58–60^.

**Fig. 4.**
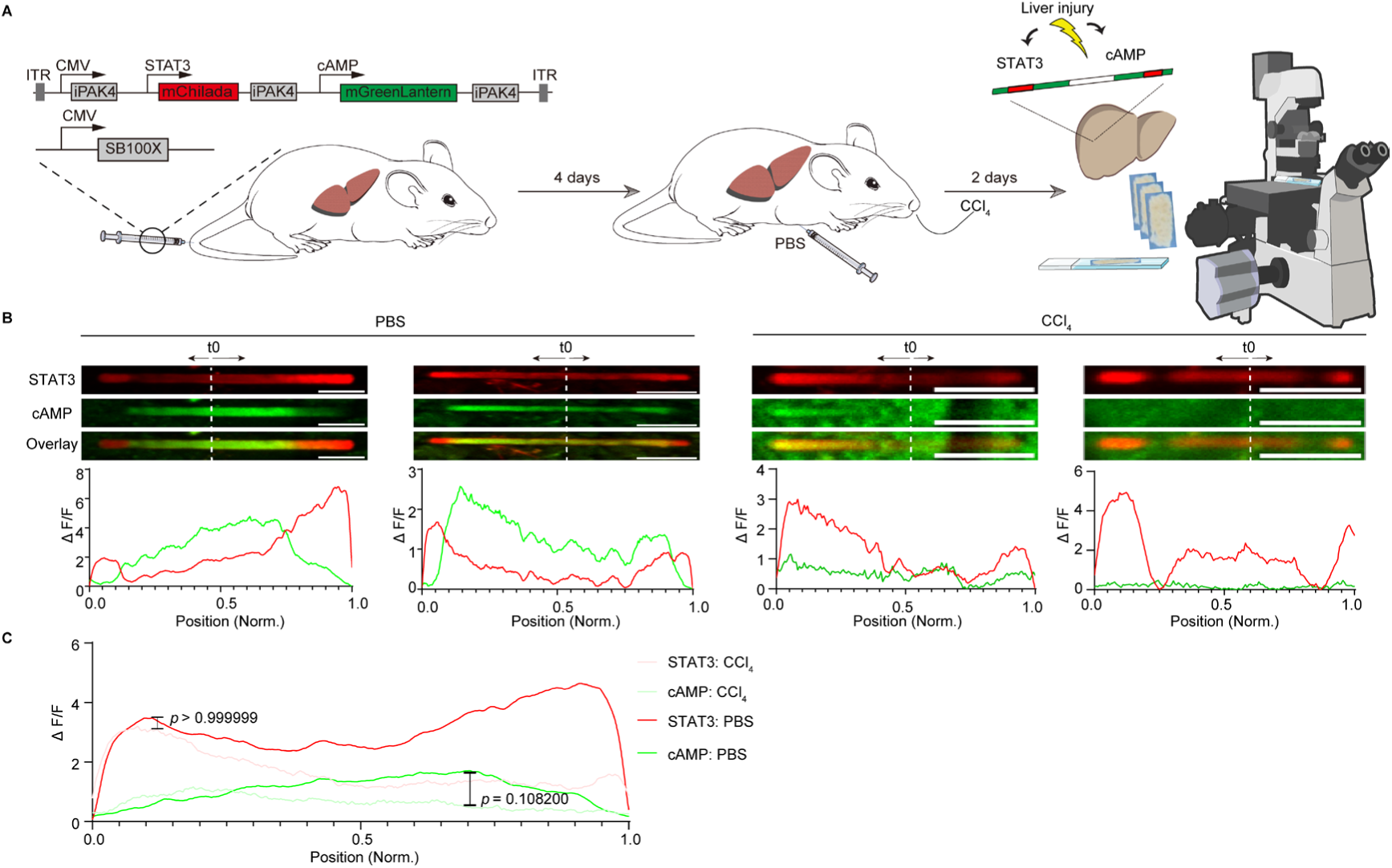
Multiplexed recording of STAT3 and cAMP signaling in a mouse model of liver injury. **A**, The “all in one” plasmid for reporting STAT3 and cAMP activity in ticker tape was delivered with SB100X into male C57BL/6 mice through hydrodynamic tail vein injection. After 4 days, mice were i.p. injected with PBS (vehicle control) or administrated orally with 1mg/kg CCl_4_. iPAK4 fibers were imaged from dissected livers after 2 days. **B**, Representative images and corresponding ΔF/F profiles of the fiber dissected from the mouse liver in PBS group (n = 12 fibers from 4 mice) and CCl_4_ treated group (n = 7 fibers from 3 mice). Red solid lines indicate STAT3 activity, and green lines indicate cAMP activity. t0 and white dash line, the start timepoint of each fiber growth; black arrow, the direction of fiber growth. Scale bars, 10 μm. **C**, Averaged ΔF/F profiles of STAT3 and cAMP activity in control (PBS) group and CCl_4_ treated group. Statistical significance (*p*-values) was determined via a multiple unpaired *t*-test. Red solid line, STAT3 activity; green solid line, cAMP activity. *P* > 0.9999 of STAT3 fold changes between control group and CCl_4_ treated group; *p* = 0.1082 of cAMP fold changes between control group and CCl_4_ treated group.

We initially explored the use of AAVs for liver delivery. While tail-vein injection of AAV-DJ-CAG-iPAK4 and AAV-DJ-STAT3-mEGFP-iPAK4 in 9:1 ratio resulted in detectible mEGFP-iPAK4 expression in both PBS– and LPS-treated groups, we observed no fiber formation in the mouse liver (**Fig. S28**). We hypothesized that the long-distance delivery via AAVs might hinder the maintenance of the precise ratio required for iPAK4 nucleation and fiber formation within targeted organs. Indeed, upon stereotaxic injection of a specific AAV combination (hSyn-iPAK4: cFos-mEGFP-iPAK4: NFκB-mChilada-iPAK4 in 8:1:1 ratio) into the mouse visual cortex, abundant two-color fiber formation was observed (**Fig. S29**). Although, wide variability of intensity profiles for individual fibers hindered population analysis of intensity changes on large population level, these results demonstrated ability of FPTT fiber formation in neurons in vivo.

To advance towards *in vivo* applications while minimizing the number of required vectors, we engineered a single expression unit for parallel P_STAT3_– and P_CREB_-specific FPTT recording. This unit was subsequently cloned into a Sleeping Beauty (SB)-based transposon system (**Fig. S30A**). We hypothesized that the expression levels driven by the CMV promoter within this single unit would be sufficient for the nucleation compared to both STAT3– and cAMP-responsive FPTT components, thereby obviating the need for multi-plasmid co-transfection at an 8:1:1 ratio for *in vivo* studies. We first expressed and validated this single unit in MOR9-1-transgenic stable cell lines treated with VA or IL-6, as previously described (**Fig. S30 B**). Quantification of intensity profiles and maximum fold changes for each fiber confirmed that VA and IL-6 independently activate cAMP and STAT3 signaling pathways, mirroring the results obtained from co-transfection with multiple plasmids at an 8:1:1 ratio (**Fig. S30C-E**).

To enhance expression efficiency in the mouse liver, this single FPTT transposon plasmid was co-delivered with the hyperactive SB100X transposase^61^ into male C57BL/6 mice via hydrodynamic tail vein injection. After 4 days, mice received 1 mg/kg CCl_4_ or PBS (vehicle control) for an additional two days. Subsequently, iPAK4 fibers were imaged and analyzed from dissected liver slices (**Fig. 4A**). While both cAMP and STAT3 signals were robust in the absence of drug-induced liver injury (PBS treatment), cAMP activity was notably suppressed during acute liver damage induced by CCl_4_ treatment (**Fig. 4B**). Altogether, these results demonstrated capability of multi-color FPTT fiber formation in vivo in liver and in neurons in the brain with potential to read out physiological promoter activation.

#### FPTT reveals an unexpected crosstalk between STAT3 and NFAT pathways

To test whether FPTT can be used for systematic analysis of temporal dynamics and interdependence between any pair of signaling pathways, we applied an analogous cell-based experimental scheme on STAT3– and NFAT-specific promoters. Specifically, we first co-transfected HEK-293T cells with a constitutive iPAK4 expression vector (90%, w/w) and a P_STAT3_-driven mScarlet3-iPAK4 expression vector (10%, w/w) or a P_NFAT_-driven mGreenLantern-iPAK4 expression vector (10%, w/w) independently. Averaged fluorescence intensity changes (ΔF/F) were monitored over 10 h of stimulation with IL-6 (500 ng/ml) or KCl (45 mM). While our results recapitulated the well-known finding that IL-6 was specific for STAT3-signaling^62,63^, KCl surprisingly activated both STAT3– and NFAT-driven promoters (**Fig. S31A-E**). This finding was also in line with the results obtained from an analogous cell-based reporter assay, in which a P_STAT3_-driven or P_NFAT_-driven NanoLuc expression vector was used as a quantitative readout (**Fig. S31F, G**).

Evidently, the correlation between STAT3 and NFAT signaling during KCl-dependent Ca^2+^ stimulation was also observed in multiplexed FPTT experiments when both promoters each driving a different mFP-iPAK4 fusion were expressed in the same cell. When HEK-293T cells were co-transfected with a constitutive iPAK4 expression vector (80%, w/w), a P_STAT3_-driven mScarlet3-iPAK4 expression vector (10%, w/w) and a P_NFAT_-driven mGreenLantern-iPAK4 expression vector (10%, w/w) prior to treatment with either IL-6 (500 ng/ml) or KCl (45mM) (**Fig. S32A,B**), analysis of fluorescent signals recorded onto the FPTT fibers revealed that IL-6 specifically activated STAT3 promoter activity with around two folds (represented by red signals). This specificity was observed despite the higher basal expression levels of STAT3, consistent with previous findings (**Fig. S32 C**). Conversely, KCl stimulation was capable of inducing activation of both STAT3– and NFAT-reporter systems (indicated by green and red signals) **(Fig. S32C, Figure S31 D, E)**.

Furthermore, when the cells were first stimulated with IL-6 (500 ng/ml) for 8 h and then switched to stimulation with KCl (45mM) for another 8 h (**Fig. S32 D**), a clear asynchronous response could be observed between IL-6-dependent red patterns (STAT3 activity) and KCl-dependent green patterns (NFAT activity) (**Fig. S32 E**). When we switched the treatment order to KCl first and then IL-6, a different color pattern was recorded as KCl activated both STAT3 and NFAT, while IL-6 stimulation only boosted STAT3 activity (**Fig. S32 F, G**). Such temporal resolution are unique advantages of the FPTT technology over conventional molecular recording technologies and/or cell-based reporter assays **(Fig. S31F,G)**, as FPTT are not only able to cellular response within a fixed time interval but also capture multiple transient events at the same time.

### Design and validation of a FPTT-compatible promoter responsive to mTOR signaling

In contrast to NFκB and other signaling pathways that exhibit a seemingly all-or-nothing or “digitalized” activation dynamics^64,65^, the eukaryotic mechanistic target of rapamycin (mTOR) pathway is constantly switching between growth and starvation states^66^. Recently, it was further shown that mTOR activity oscillates throughout the cell cycle, promoting progression through interphase and entry into mitosis^67^. To measure such dynamics, conventional technologies ranging from Western Blot to immunofluorescence may only capture distinct cell states and prohibit real-time imaging of transient activities^67^. To investigate whether the FPTT technology is also amenable to the recording of such complex pathway behaviors, we first created a synthetic promoter containing response elements for sterol regulatory-element binding protein 1 (SREBP1; **Fig. 5A**). Because the mTOR pathway is a major upstream effector of nuclear localization and activation of SREBP1-dependent transcription^68^, we reasoned that promoters specific for the N-terminal domain of SREBP1 (nSREBP1) could be used to drive mFP-iPAK4 expression and generate mTOR-specific FPTT readouts (**Fig. 5A**). Specifically, the candidate nSREBP1-specific promoters were engineered to harbor different tandem motifs of sterol regulator element-1^69,70^ (SRE-1, 5’-ATCACCCAC-3’) and/or E-box motifs^71^ (5’-ATCACGTGA-3’) placed upstream of a minimal human cytomegalovirus promoter (P_hCMVmin_). When validated using a cell-based assay involving human placental secreted alkaline phosphatase (SEAP) as a reporter gene, the promoter variant P_mTOR3_ containing three copies of SRE-1 and two copies of E-box exhibited the highest dynamic range, achieving up to 8.9-fold of SEAP expression when comparing the experimental states of mTOR inhibition (stimulation with 500nM Torin-1)^72^ and mTOR activation (stimulation with 5µM MHY1485^73^) **(Fig. 5B)**.

**Fig. 5.**
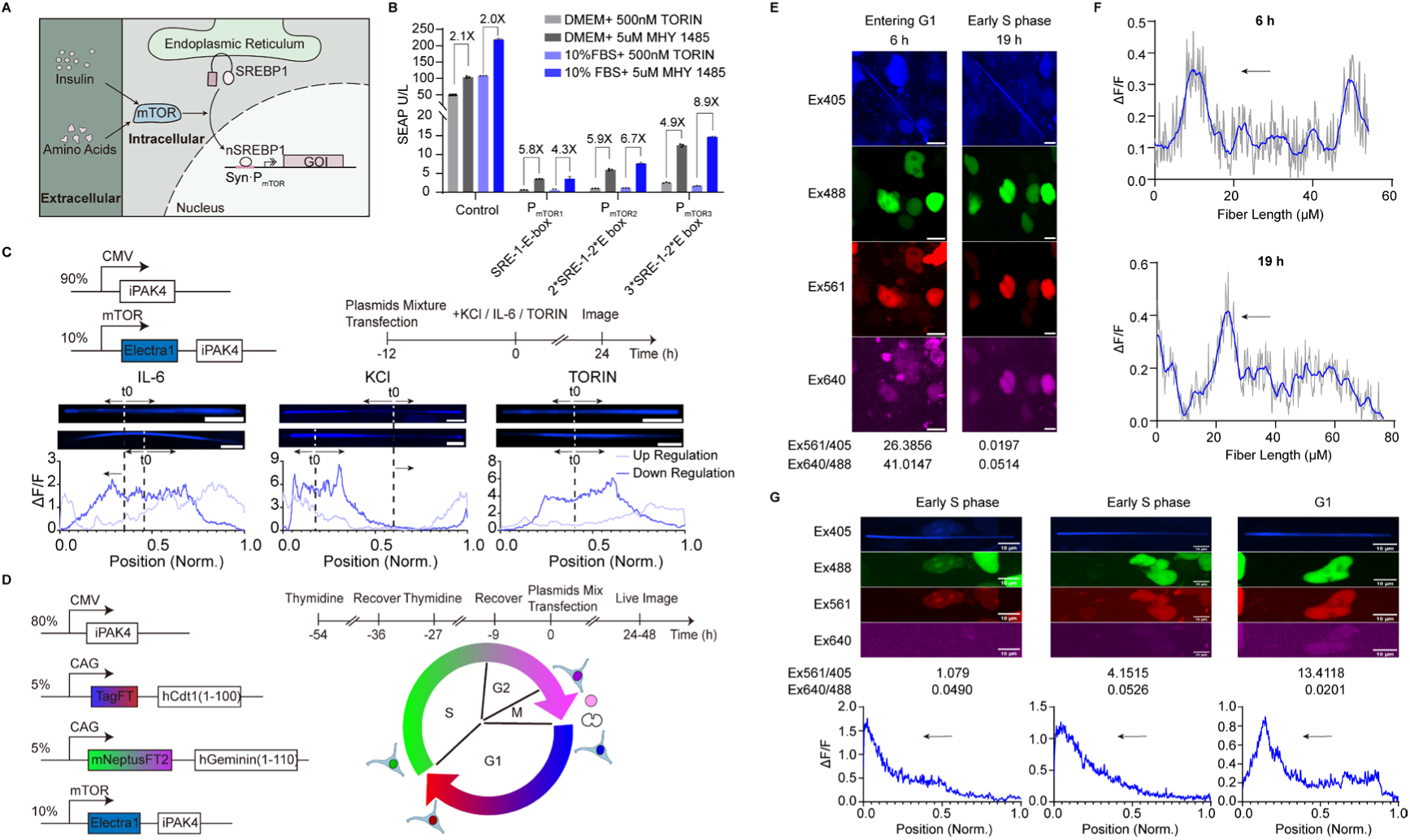
Design and validation of a FPTT-compatible promoter responsive to mTOR signaling. **A**, Scheme of mTOR signaling based SREBP1 protein cleavage. The mTOR kinase located upstream of SREBP1 is activated by insulin, amino acids, etc., and promotes SREBP1 protein cleavage. The active form of SREBP1, nSREBP1, is translocated to the nucleus and binds to nSREBP1-specific promoters and drive gene of interest expression. **B**, nSREBP1-specific promoters response to mTOR inhibitor (Torin1) and mTOR agonist (MHY1485) in DMEM medium supplemented with or without 10% (v/v) FBS. HEK-293 cells were transfected with 200 ng plasmids containing different promoter variants and cultivated in cell culture medium containing mTOR inhibitors (500nM Torin-1) or mTOR activators (5µM MHY1485) either in the presence or absence of 10% FBS. SEAP levels in culture supernatants were measured at 36h post transfection. (pDJ71-pmTOR_SRE-1-E-box_-SEAP-pA, pDJ72-pmTOR_2*SRE-1-2*E-box_-SEAP-pA, pDJ73-pmTOR_3*SRE-1-2*E-box_-SEAP-pA and pMX316, control). Data are shown as the mean±SD of n=3 independent experiments. **C**, The expression cassette and experimental protocol used for recording mTOR activity in IL-6, KCl, and Torin-2 treated HEK293T cells. 12 hours post-transfection, cells are treated with 500 ng/ml IL-6, or 45 mM KCl, or 250 nM Torin-2 for 24 hours before images were acquired. Representative image and corresponding ΔF/F profiles of the fiber were shown. t0 and dash line, the start timepoint of each fiber growth; black arrow, the direction of fiber growth. “upregulation” refers to a high intensity (activated pattern) observed at the final time points imaged, while “downregulation” indicates a lower intensity (inhibited pattern) at the final time points imaged. For up-regulation traces, N = 7 fibers in IL-6 treated, and n =21 fibers in KCl treated, and n = 5 fibers in Torin-2 treated; for low-regulation traces, n = 14, 23, 17 fibers, respectively, from 3 biological replicates. Scale bars, 10 μm. **D**, The expression cassette and experimental protocol used for recording mTOR activity in double thymidine block synchronized HEK293T cells. FucciFT plasmids, with tagFT to report G1 phase under excitation of 405 and 561 nm, and mNeptusFT2 to report S to G2 phases under excitation of 488 and 640 nm were co-transfected with mTOR ticker tape plasmids into HEK293T cells 9 hours after double thymidine block. Live images were acquired 24 hours post-transfection and lasted for 24 hours. **E**, Representative images at G1(6 h post-start of imaging) and S phase (19 h post-start of imaging) with mTOR ticker tape. The ratio of Ex561/405 represents tagFT transitions in G1 phase, and Ex640/480 represents mNeptusFT2 transitions from S to G2 phase. Scale bars, 10 μm. **F**, Corresponding ΔF/F profiles in images E at G1(6 h post-start of imaging) and S phase (19 h post-start of imaging) of mTOR ticker tape. Grey lines, raw files; Blue lines, smooth profiles by averaging 15 values on each side using a second order smoothing polynomial. Black arrow, the direction of fiber growth. **G**, Representative images and corresponding ΔF/F profiles at 24 hours post the start of imaging. Black arrow, the direction of fiber growth. Scale bars, 10 μm.

Thus, we used FPTT to examine how mTOR signaling may respond to various environmental stimuli. Hence, HEK-293T cells were transfected with a P_mTOR3_-driven mElectra1-based FPTT system and exposed to various external stimuli in culture, such as IL-6, KCl, or Torin **(Fig. 5C)**. A distinct bimodal pattern of intensity profiles was observed in KCl-treated cells. Approximately 50% of the recorded fibers exhibited high fluorescence intensity in the middle and low intensity at the tips, indicative of higher mTOR activity at the onset of recording and lower activity towards the end (termed as ‘up-regulation’). Conversely, the remaining fibers displayed the opposite pattern, with low intensity at the beginning and high intensity at the end (termed as ‘down-regulation’; **Fig. 5C**). Similar intensity profile patterns were observed in IL-6 treated cells, however, with a 1:2 ratio of up to down-regulation of mTOR activity (**Fig. 5C**). In Torin-treated cells, we observed only a minor group of fibers (∼20%) showing elevated mTOR signals at the tip of fibers (**Fig. 5C**). Conversely, in untreated cells, we observed a greater heterogeneity in intensity profiles. Many of these profiles exhibited multiple distinct peaks, indicative of both up– and down-regulation, and lacked any predominant or consistent pattern (**Fig. S33**). To facilitate quantitative comparison and ratio calculation with other treated groups, these diverse profiles were nonetheless categorized into two predefined formations (**Fig. S33**).

To explore whether such bimodal activity profiles of mTOR signaling was related to oscillating dynamics throughout the cell cycle^67^, we co-expressed the P_mTOR3_-driven mElectra1-based FPTT system with a previously developed FucciFT system capable of reporting distinct cell cycle states based on ratiometric imaging in several standard channels^74^. Specifically, HEK293T cells synchronized on a double thymidine block were co-transfected with a constitutive iPAK4 expression vector (70%, w/w), a P_mTOR3_-driven Electra1-iPAK4 expression vector (10%, w/w) and 10% w/w of FucciFT plasmids, before single-cell live imaging was performed for 24 h **(Fig. 5D)**. After careful analysis, we found that some cells had low mTOR activity when entering the G1 phase at 6 hours from the start of imaging, but markedly boosted mTOR activity when entering the S phase (correlation between blue fluorescence signals on the fiber and green fluorescence produced by FucciFT; **Fig. 5E, F)**. We also characterized several fibers at the end of 24 h in live imaging, while in the S phase with a close number of fluorescence dividing with Ex 640/488, the mTOR activity was higher, and in the G1 phase, the mTOR activity went down **(Fig. 5G)**. In summary, this demonstrates that the P_mTOR3_-driven FPTT system can effectively capture the intricate cell cycle dynamics of mTOR signaling, offering a resolution typically unavailable from analyses of bulk cell populations using time series blotting.

### Imaging and interpretation of highly multiplexed recording

Finally, we explored how the FPTT technology can be used to monitor multiple signaling pathways in parallel. For example, it is known that JAK/STAT3, NFAT, mTOR, and NFκB are four major pathways involved during T-cell activation^75^, but a functional or temporal dependence between these pathways remains unexplored. To investigate this question using the FPTT, we first examined whether the expression of iPAK4 fibers could elicit an immune response on targeted signaling pathways. Utilizing a NanoLuc assay, both with and without iPAK4 expression, we observed that iPAK4 fiber formation had no discernible effect on STAT3, NFAT, or NFκB activation (**Fig. S34**). However, we noted an approximate two-fold increase in mTOR activation (**Fig. S34**). We further validated that all four signaling pathways remained intact and capable of producing fluorescent fibers in Jurkat T cells following PMA and ionomycin treatment, whereas no activation or fiber formation occurred in the absence of treatment (**Fig. S35**). Building on this, we demonstrated that a single iPAK4-based fiber in HEK293T cells can indeed accommodate at least three distinct fluorescent readouts, enabling multiplexed recording based on specific treatment conditions (**Fig. S36; Fig. S37**), and that calculation of Pearson’s correlation coefficients between fluorescence intensity profiles in the specific channels is an effective way to interpret the functional dependence of two signaling pathways **(Fig. S38)**. Specifically, our data reveal that NFAT and mTOR had no correlations in any treatment, while STAT3 had strong correlations with mTOR in the Torin-treated group **(Fig. S38)**.

Next, we stably integrated expression cassettes for P_STAT3_-driven mScarlet3-iPAK4, P_NFAT_-driven mGreenLantern-iPAK4, P_NFkB_-driven miRFP670-iPAK4 and P_mTOR3_-driven Electra1-iPAK4 into Jurkat T cells genome using the Sleeping Beauty transposase technology, thus establishing a polyclonal cell line (Jurkat-4S) custom designed for multiplexed recording of four promoters in parallel **(Fig. 6A)**. The multiplexed recording was initiated by transient transfection of CMV-iPAK4 followed by treatment with PMA and ionomycin 24 h later to activate T cells^76^**(Fig. 6A,B)**. FPTT fibers were analyzed by fluorescence microscopy and intensity profiles for each channel were measured **(Fig. 6C)**. While in control group without PMA and ionomycin treatment, no fiber formation was observed under a broad-field of view (**Fig. S39**). In treated groups, results show that all four signaling pathways were activated with variable orders in a cell-by-cell case, potentially due to the heterogenous expression levels in the cell population **(Fig. 6C)**. To analyze the correlation in temporal promoter activities by means of dimensionality reduction, we calculated the Pearson correlation coefficients for each pair of intensity profiles, which were presented in 4 by 4 matrices containing six unique pairwise combinations for four promoters **(Fig. 6D)**. We foundSTAT3, NFAT, and mTOR had a consistently high positive correlation with Pearson’s correlations above 0.5 in many fibers (**Fig. 6D**). Particularly high correlation (Pearson’s coefficient above 0.7) was observed for STAT3-mTOR pair in all analyzed fibers (**Fig. 6D**). Population data for 8 analyzed fibers revealed high positive correlations for all pairwise combinations of STAT3, NFAT, and mTOR promoters **(Fig. 6E)**. Thus, in the present case, our data suggest that STAT3, NFAT, and mTOR may be coupled during experimental T-cell activation, indicating the involvement in co-occurring events that mediate synergistic regulation and/or functional convergence^77^. Evidently, though we showed that FPTT has the potential for “high-content” recording of complex cellular physiologies by generating multiple quantitative fluorescent readout signals by a single experiment, further characterization work is required to corroborate such technology and reveal more accurate information on the “true” signaling dynamics in other cell biology contexts.

**Fig. 6.**
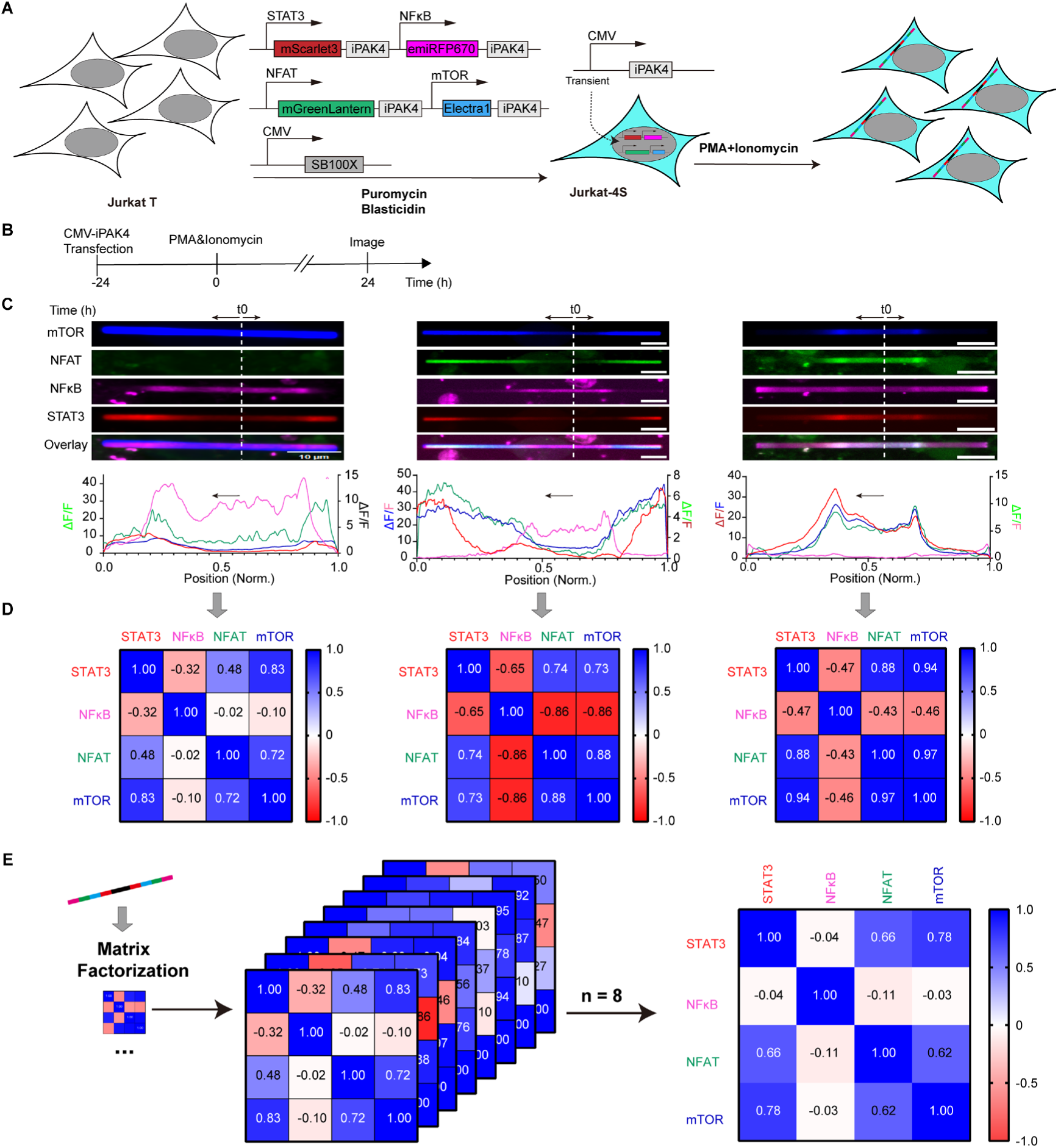
Multiplexed recording of STAT3, NFAT, NFkB, and mTOR activities in Jurkat cells. **A**, Workflow for creating the polyclonal Jurkat_4S_ cell line. Native Jurkat cell were co-transfected with 230 ng each of pDJ655-ITR-pNFAT-mGreenLantern-iPAK4-pA-pmTOR_3*SRE-1-2*E-box_-iPAK4-pA-ITR and pDJ656-pSTAT3-mScarlet3-iPAK4-pA-pNFκB-emiRFP670-iPAK4-pA-ITR and 40 ng of pCMV-T7-SB100 and selected with puromycin and blasticidin. Jurkat_4S_ cell line co-transfected with CMV-iPAK4 and treated with PMA and Ionomycin for multiplexed signaling recording. **B**, Experimental protocol for recording multiplexed STAT3, NFAT, mTOR and NFκB signaling. Transfected Jurkat_4S_ were exposed to stimulation with PMA (50 ng/mL) and Ionomycin (1 µg/mL) for 24h before images were acquired. **C**, Representative images, corresponding ΔF/F profiles of the fibers treated with PMA and Ionomycin. Blue line, mTOR; Green line, NFAT; Magenta line, NFκB; Red line, STAT3. t0 and white dash line, the start timepoint of each fiber growth; black arrow, the direction of fiber growth. Scale bars, 10 μm. **D**, Person correlation coefficient corresponded to representative images in C. **E**, Averaged person correlation coefficient (n = 8 fibers from 3 biological replicates).

## Discussion

Protein-based ticker tapes provide an alternative approach for recording, storing, and interpreting cellular histories of a specific target promoter. In this report, we demonstrated the applicability of protein-ticker tape technologies to enable simultaneous recordings of multiple signaling pathway dynamics within single cells. The core innovation of FPTT lies in its integration of physiologically relevant synthetic promoters with self-assembling iPAK4 protein fused with spectrally distinct monomeric FPs. Reminiscent of a “black box recorder device”, this design enables the formation of multicolor fluorescent fibers within cells, facilitating the visualization, quantification, and activity correlations of up to four signaling pathways through simple decoding using conventional fluorescence microscopy.

While our approach shares a conceptual similarity with previous iPAK4 studies, the original HaloTag-iPAK4 system relies on the delivery of exogenous dyes (e.g., JF dyes). This dependency introduces limitations such as restricted bioavailability and compromised temporal precision, particularly in *in vivo* applications. Critically, the original iPAK4 system validated its capabilities using externally applied perturbations (e.g., drug-induced gene expression), which inherently lead to synchronized cellular responses across a population. In contrast, our system employs monomeric FPs, facilitating less invasive *in vivo* experimentation and uniquely enabling the recording of spontaneous or asynchronous activation of reporter genes. As demonstrated by our studies on cFos and NFκB, cAMP and STAT3, and STAT3 and NFAT, FPTT can capture signaling dynamics under both spontaneous (e.g., control condition without any treatment) and asynchronously treated conditions. Furthermore, the integration of fluorescent proteins in FPTT allows for multiplexed readouts of transcriptional histories, capturing signaling dynamics not only at the endpoint but also in real-time. In contrast, the original iPAK4 system necessitates the sequential application of various dyes at different time points, requiring endpoint data acquisition after cell fixation to prevent data misinterpretation. The iterative process of washing and administering different dyes significantly complicates the monitoring of real-time events and continuous observation of native environments. Moreover, the unpredictable nature of biological events makes it nearly impossible to determine optimal dye-switching timings, and the reliance on dye addition raises concerns about potential dye interference. Our system, conversely, visualizes naturally occurring events as intrinsic fluorescent protein expressions on a “ticker tape” independent of the orthogonality of biological events.

The dye-switching mechanism within the iPAK4 framework primarily serves to generate timestamps for endpoint readouts, such as the absolute timing of PMA-induced cFos driven mEGFP transcription and translation. Our FPTT system, however, supports the observation of multiple biological events in chronological order, providing insights into their relative timing. Our findings robustly demonstrate FPTT’s capacity to detect responses to transient treatments lasting minutes, as well as prolonged exposures extending over hours or even days, with recording times exceeding 8 days (**Supplementary Table 1**). This remarkable sensitivity and extended recording capability provide unprecedented insights into the relative timing between cFos and NFκB responses to stimuli, and the crosstalk between STAT3 and NFAT during KCl treatments—insights not previously achievable or reported.

Signaling pathways are essential for cells to respond and adapt to dynamic environmental changes, particularly in the context of disease^77–80^. However, currently available tools for studying instant activation levels and/or interdependencies between different pathways at the same time remain limited. To this end, the FPTT technology was developed to accurately report the relative timing of activation events for multiple cellular events in parallel in response to specific stimuli, providing a valuable tool for dissecting complex signaling pathways and understanding the intricate dynamics of cellular responses. In fact, we were able to provide the technological basis to reliably obtain fluorescent ticker tapes as a type of “raw data” for each recording event, but future work is evidently required to translate such raw data into biologically valid information. Here, we propose the use of linear correlation between the temporal dynamics of two promoters within a selected time window to quantify potential cross-talk and orthogonality between pathways. Though we showed that this method could be flexibly expanded to generate correlation matrices for up to 4 target promoters in parallel, further validation of the technology is necessary for gaining new biological insight into cellular physiology. Specifically, the data obtained in this study may fall in line with some known findings reported in the fields, such as revealing potential crosstalk between NFκB and cFos pathways in neurons^53^ or functional orthogonality between the STAT3– and cAMP-responsive promoters used to generate corresponding FPTTs^47^. However, the experiments described in this present work were still insufficient to “clarify” specific insights about specific signaling pathways, such as the particular roles of NFκB and NFAT signaling during T-cell activation^81^**(Fig. 6)**, or the biological relevant of cAMP and STAT3 signaling during normal liver physiology and disease^58,60^**(Fig. 4)**. Nevertheless, the FPTT recording system presents improvements in extending available promoters and multiplexing capabilities, with challenges such as long-term stability and minimal photodamage also requiring further addressing. In consideration of these developments, our findings contribute to the field by fostering a deeper understanding of cellular processes and offering a versatile approach for researchers to expand their investigative horizons. The simplicity, modularity and scalability of this approach make it a valuable tool for a wide array of applications in both basic and applied life sciences.

FPTT was developed to fulfill a long-standing objective in biological research, enabling the tracking of cellular histories across various physiological and pathological contexts with high accuracy and temporal resolution. From a practical perspective, FPTT is accessible, easy to scale up, and straightforward to read out and analyze the data. Taken together, this approach significantly lowers the barrier for independent users to adopt this technology. However, future work will be necessary to mitigate the observed cell-to-cell variability in expression levels within co-transfection systems. The application of synthetic biology tools, such as engineered synthetic circuits, offers a promising solution to buffer gene dosage variation among individual mammalian cells. In this aspect, FPTT shares the similar advantage of genetically encoded integrators for neuronal activity and neurotransmission^82,83^, which became ubiquitous tools in neuroscience research. However, unlike integrators that only label cells activated during user-defined time windows typically ranging from hours to 2-3 days^82^, FPTT enables continuous recording of neuronal activity for up to 8 days with a temporal resolution of about 3 hours (**Fig. 2**). At the same FPTT are more universal compared to existing integrators and can be readily extended to any cell types and range of activities for recording. We believe that these unique advantages will make the FPTT tool popular for various applications in cell and synthetic biology.

## Methods

### Molecular cloning and plasmid construction

The cFos-eGFP-iPAK4 and CMV-iPAK4 plasmids were acquired from Addgene (plasmid #177883 and #177880, respectively). Additional plasmids for reporting activity, including pTS566 for STAT3, pMX57 for NFAT, and pKR32 for NF-κB activities, were generously provided by Dr. Leo Scheller. The pDJ73 plasmid, which reports mTOR activity, was synthesized by Tsingke Company in China. We replaced the original SEAP genes under specific promoters with FP-iPAK4 using restriction enzymes from New England BioLabs (NEB, USA) and a One Step PCR Cloning Kit from Novoprotein (China).

The 1POK gene was *de novo* synthesized by Tsingke Company (China), according to the sequences reported in the original paper^23^. The genes of mEGFP (pCytERM-mEGFP-N1, #0000281), mNeonGreen (pAAV-CAG-mNeonGreen-P2A-FusionRed, #0000264), mGreenLantern (pAAV-CAG-mGreenLantern-P2A-FusionRed, #0000278)), mCherry (AttB-mCherry, #0000253), Electra1 (pAAV-CAG-Electra1-P2A-GFP, #0000197) were cloned from the barcoded plasmids provided by WeKwikGene (WKG, China) into their respective ticker tape vectors using custom designed primers following standard molecular cloning protocols. The mVenus, mAvicFP1, mScarlet-I, mScarlet3, mChilada gene were also synthesized by Tsingke Company following fpbase.org sequences.

The cumate inducible system was purchased from the Systems Biosciences company (SBI, USA) and cloned into an AAV vector (pAAV-CAG-mNeonGreen-P2A-FusionRed, #0000264) from WKG. Primers are synthesized by YouKang Biotechnology (Hangzhou, China).

Plasmids were constructed using NovoRec^®^ plus One Step PCR Cloning Kit (Novoprotein, China) with DNA amplified by PrimeSTAR^®^ Max DNA Polymerase from Takara (Japan). Plasmid backbones were digested using NEB restriction enzymes, and constructs were transformed into TOP10 competent cells (BioMed, China) for amplification. Sequencing was conducted by YouKang Biotechnology.

The mammalian plasmids generated in this study are available from the WeKwikGene plasmid repository at Westlake Laboratory (https://wekwikgene.wllsb.edu.cn/) with following barcodes #470-511.

### HEK293T and HeLa Cell Culture

HEK293T (ATCC^®^ ACS-4500™)) and HeLa cells (ATCC^®^CCL-2^TM^) were maintained using standard cell culture protocols. HEK cells (passage number <10) and HeLa cells were plated at 50% confluency onto 10-cm culture-treated dishes (Fisher Scientific, USA). The culture medium consisted of Dulbecco’s modified Eagle medium (DMEM, Wuhan Servicebio Technology, China) supplemented with 10% (v/v) fetal bovine serum (FBS, ExCell Bio, China) and 1% (v/v) penicillin-streptomycin (Yeasen Biotechnology, China). Cells were incubated at 37°C with 5% CO2.

When cells reached 70%-90% confluence, cells were seeded into the 24-glass bottom well plate (Cellvis, USA) pretreated with Matrigel (BD Biosciences, USA) for 30 min at 37°C. 1 μg plasmids containing CMV-iPAK4 (90%, w/w) and FP tagged iPAK4 with specific promoters (10%, w/w) were mixed with 2 μl Hieff Trans Liposomal Transfection Reagent (Yeasen Biotechnology, China) in opti-MEM (Gibco, USA), incubated for 15 minutes, and dropwise added into HEK293T cells. 4 hours after incubation, the medium was replaced with fresh DMEM containing 10% FBS.

To activate STAT3 in HEK293T cells, interleukin 6 (IL-6) (GenScript, Z03134) was used. A stock solution of 50 µg/mL in PBS was diluted to 500 ng/mL in DMEM prior to application. For mTOR inhibition, Torin-1 (MCE, HY-13003) and Torin-2 (MCE, HY-13002) were utilized. A 10 mM stock solution was diluted to 250 nM in DMEM before being added to the cells. To stimulate NFAT activity, potassium chloride (KCl) (ST345, Beyotime) was employed. A 3 M stock solution was diluted to 45 mM in DMEM before use. For activating cAMP pathways, vanillic acid (VA, Rhawn, R017640, China) was used. A 1 M stock solution was diluted to 500 μM in DMEM prior to adding to the culture. All stock solutions were prepared and stored following the manufacturer’s guidelines.

### Jurkat Cell Culture

The Jurkat 6-1 cell line (ATCC^®^TIB-152™) was grown in RPMI 1640 medium (Sartorius AG, Gottingen, Germany) supplemented with 10% (v/v) FBS and 1% (v/v) penicillin/streptomycin. The cells were maintained at 37°C in a humidified atmosphere containing 5% CO_2_. To create the polyclonal Jurkat_4S_ cell line, native Jurkat cells (1.0 ×10^5) were co-transfected with 230 ng each of pDJ655 and pDJ656, and 40 ng pCMV-T7-SB100 using Lipofectamine LTX (ThermoFisher Scientific, USA). Selection was carried out with 1 µg/mL puromycin (ThermoFisher Scientific, USA) and 5 µg/mL blasticidin (Beyotime, China).

Once the cells achieved 70-90% confluence, either native Jurkat cells or Jurkat_4S_ cells were seeded in 24-well plates (Cellvis, USA). The plates were pretreated for 30 minutes at 37°C with Poly-D-lysine (Beyotime, China). Plasmid transfection was conducted using Lipofectamine LTX. After 24 hours, the medium was replaced, and PMA (50 ng/mL) along with Ionomycin (1 µg/mL) (Dakewei, China) was added to the cells to activate Jurkat cells.

### Neuron Culture

Primary mouse hippocampal neurons were isolated from newborn (P0) C57BL/6 mice without regard to sex (according to protocol no. 19-044-KP-2 of Westlake University’s Institutional Animal Care and Use Committee). These neurons were then cultured in a 24-well glass-bottom plate (Cellvis, USA), pretreated with Matrigel (BD Biosciences, USA), following previously established methods^84^. The culture medium consisted of Neuronal Basal-A medium (Gibco, USA) supplemented with B27 (Gibco) and 1X GlutaMax (Gibco). The medium was refreshed every 2-3 days by replacing half the volume. Neurons were transfected at 5 days in vitro (DIV5) using the Calcium-Phosphate method^85^.

To activate cFos, we utilized phorbol 12-myristate 13-acetate (PMA, InvivoGen) prepared as a 5 mg/ml stock solution in dimethyl sulfoxide (DMSO, Sigma) and diluted to the desired concentration in the culture medium, as specified in the figure legends. For cumate-controllable system activation, cumate water solution (SBI, USA) was diluted to the desired concentration in the culture medium, as outlined in the figure legends. To activate NF-κB in neurons, we used mouse-derived tumor necrosis factor alpha (TNF-α, GenScript), prepared as a 20 µg/ml stock solution in phosphate-buffered saline (PBS, Sevicebio), and diluted to the specified concentrations in the figure legends. All chemicals were stored according to the manufacturer’s recommendations.

### SEAP and NanoLuc Assay

Expression levels of human placental secreted alkaline phosphatase (SEAP) in culture supernatants were quantified by measuring the colorimetric absorbance time course of the SEAP-mediated p-nitrophenyl phosphate (pNPP) (Aladdin, Shanghai, China, P109039) to p-nitrophenolate conversion. In brief, 80 µL of heat-inactivated supernatants (30 min at 65℃) mix to 100 µL of 2x SEAP assay buffer and 20 mM pNPP per well in a 96-well plate. The light absorbance time course was measured at 405 nm (37℃) using a Thermo Scientific^TM^ Multiskan^TM^ Sky microplate reader.

Expression levels of NanoLuc were profiled using the Nano-Glo^®^ Luciferase Assay system (Promega, Madison, WI, USA; Cat# N1120) by adding 7.5 µL of sample mix to 7.5 µL 50:1 (v/v) buffer-substrate per well in a black 384 well plate, followed by luminescence recording on a Thermo Scientific^TM^ Fluoroskan^TM^ FL microplate reader.

### Mouse liver

All animal procedures adhered to Westlake University’s guidelines and were approved by their Institutional Animal Care and Use Committee (protocol no. AP#25-048-XMQ). 8-week-old male C57BL/6 mice weighing 25-26 g were selected for hydrodynamic tail vein injection. 100 µg pDJ750 and 5 µg SB100X were diluted in 2.4 mL (8%-10% body) of Ringer solution and injected into mice between 5 and 8 seconds. 4 days after hydrodynamic tail vein injection, the mice were either injected PBS intraperitoneally as control group or administrated orally with 1mg/kg CCl_4_ as treated group. 2 days after PBS injection, mice were sacrificed. Perfused with 4% PFA, liver was isolated and further fixed in 4% PFA for 2 hours at 4℃, cut using a Leica VT1200S vibrating blade microtome at a speed 1.2 mm/s, amplitude 1mm, and thickness of 50 µm.

### Mouse brain

All animal procedures adhered to Westlake University’s guidelines and were approved by their Institutional Animal Care and Use Committee (protocol no. AP#19-044-KP-2). All experimental procedures were conducted in strict accordance with the guidelines set forth by the Westlake University Institutional Animal Care and Use Committee (IACUC) and adhered to national regulations governing animal research. C57BL/6J mice, aged 8-12 weeks, regardless of gender, at the time of surgery, were used for all experiments.

Recombinant adeno-associated viral (rAAV) vectors, DJ serotype, expressing iPAK4 under the control of the hSyn promoter, mEGFP-iPAK4 under the control of STAT3 promoter and mChilada-iPAK4 under the control of NFκB were utilized. All the AAVs were purchased from the BrainVTA (Wuhan, China). Viral titers, determined by quantitative PCR, ranged from 1.0 × 10^13 to 3 × 10^13 genome copies (GC) per mL. Viruses were aliquoted and stored at –80°C until immediately prior to injection.

Mice were deeply anesthetized with isoflurane (4% for induction, 1-2% for maintenance) delivered in oxygen via a nose cone. Anesthetic depth was continuously monitored by assessing respiratory rate and hind limb withdrawal reflex. Body temperature was maintained at 37°C using a feedback-controlled heating pad. Ophthalmic ointment was applied to prevent corneal dehydration.

Upon induction of anesthesia, mice were secured in a stereotaxic frame (RWD, China) using non-traumatic ear bars and a bite bar. The scalp was disinfected with alternating scrubs of 70% ethanol and Povidone-iodine solution. A midline incision was made to expose the skull. The skull surface was then gently cleaned with sterile saline to visualize bregma and lambda, which were subsequently leveled in the anterior-posterior plane.

Small burr holes (approximately 0.5-1.0 mm in diameter) were drilled through the skull using a high-speed dental drill (RWD, China) over the target cortical regions. Stereotaxic coordinates, measured relative to bregma, were as follows: Visual Cortex: Anterior-Posterior (AP): –2.9 mm; Medial-Lateral (ML): ±2.5 mm; Dorsal-Ventral (DV): –0.4 mm from the pial surface.

AAVs were delivered using a glass micropipette (tip diameter 15-25 µm) attached to a 10 µL Hamilton syringe, controlled by an automated injection pump (RWD, China). For each injection site, a total volume of 250 nL of viral solution was infused at a slow rate of 50 nL/min. Following the completion of each infusion, the injection pipette/needle was held in place for an additional 10 minutes to facilitate viral diffusion and minimize backflow before being slowly retracted from the brain.

After all injections were completed, the burr holes were sealed with bone wax or dental cement, and the scalp incision was closed using tissue adhesive. Mice were placed in a recovery cage on a heating pad until fully awake and ambulatory. Animals were monitored daily for signs of pain or distress for at least 3 days post-surgery. Two weeks later, perfused with 4% PFA, brain was isolated and further fixed in 4% PFA overnight at 4℃, cut using a Leica VT1200S vibrating blade microtome at a speed 1.2 mm/s, amplitude 1 mm, and thickness of 80 µm.

### Multi-spectral imaging

Neuron cultures expressing the ticker tape fluorescent protein (Fig. 3G) were imaged using a Nikon Ti2-E widefield fluorescence microscope (Nikon, Japan) equipped with Spectra III Light Engine (LumenCore), 470/28 BP excitation filter (Semrock), an ORCA-Flash 4.0 V3 sCMOS camera (Hamamatsu), ×10 NA 0.45 and ×20 NA 0.75 objective lenses (Nikon), and controlled by NIS Elements AR 5.21.00 software (Nikon).

For multi-spectral imaging, a Nikon Spinning-Disk Field Scanning Confocal System with CSU-W1 SoRa imaging setup (Nikon) was used. This system employed 488 nm excitation for GFP, 561 nm excitation for RFP, and 405 nm excitation for BFP, with either a 40X water objective lens or a 60X oil objective lens (Nikon) for Z-stack acquisition.

### Image processing, data analysis and statistics

Image analysis was performed using ImageJ v.1.54f. Following established methods^17^. Individual fibers in the maximum intensity projections of image stacks were rotated to align their long axis with the x-axis, and fluorescence intensity profiles were plotted along the length of the fibers and exported to Excel (Microsoft Office 365, USA). Fibers that crossed near a fluorescence transition were excluded from analysis. Specifically, F_n_ represents the intensity at each interpolated point along the fibers, while F_min_ denotes the lowest intensity value among all interpolated points along the fibers. To standardize the data, the fibers are rotated to align with the X-axis, and the intensity values at each point along fibers of varying lengths are divided by their respective lengths to normalize them to a total length of 1. Each data point was recalculated as ΔF/F = (F_n_-F_min_)/F_min_, interpolated to a consistent number of X values using a VBA macro (https://doi.org/10.5281/zenodo.11514304), and averaged. Background fluorescence was subtracted from the Tet-On and Cumate-inducible ticker tapes (Fig. 1, Fig. S7), but not from the physiological ticker tape (excluding Fig.1 and Fig. S7, see source data). Final intensity profiles were generated using GraphPad Prism 10 (GraphPad Company, USA). All statistical tests and sample sizes are detailed in the figure legends. The fluorescence images presented in the main text and supplementary files represent at least two biological replicates.

## Data availability

The raw images with metadata supporting the results in all figures will be publicly available on FigShare once accepted. All plasmids utilized in this study can be obtained from WeKwikGene (https://wekwikgene.wllsb. edu.cn/). Source data are provided with this paper. The VBA macro is available from Zenodo (https://doi.org/10.5281/zenodo.11514304).

## Acknowledgements

We thank Lina Yang and Hanbin Zhang from Piatkevich lab for help with neuronal culture preparation. We thank the Laboratory Animal Resources Center and the Microscopy Core Facility at Westlake University for their support in training and data acquisition. We thank G. G. Lambert and N. Shaner for providing the mChilada gene. We are grateful to Dr. Leo Scheller for supplying the plasmids used to report several physiological activities as detailed in the Methods section. This research was supported by start-up funding from the Foundation of Westlake University, the Westlake Laboratory of Life Sciences and Biomedicine, the National Natural Science Foundation of China (grant 32050410298 and 32171093), 2020 BBRF Young Investigator Grant (28961), and the ‘Pioneer’ and ‘Leading Goose’ R&D Program of Zhejiang (2024SSYS0031) to K.D.P.

## Author contributions

K.D.P. conceived the project and together with R.W., M.X., and J.J. made high-level designs and plans, and interpreted the data. R.W. cloned the plasmids and conducted ticker-tape characterization experiments in cultured HEK293T, HeLa, and primary mouse neurons. J.J. designed the mTOR promoter, performed ticker-tape characterization in Jurkat cells and mouse liver, and verified the signaling response by NanoLuc assay. Z.L. performed 3D analysis in multiplexed signaling pathways in Fig. S24. Y. W. performed stereotaxic injections of rAAVs into mouse brain. T. L. prepared liver slices. R.W. acquired all the images and analyzed data. R.W. and K.D.P. wrote the manuscript with help from M.X. and J.J.

## References

1. Purvis, J.E., and Lahav, G. (2013). Encoding and decoding cellular information through signaling dynamics. Cell 152, 945–956. 10.1016/j.cell.2013.02.005.

2. Tang, R., Murray, C.W., Linde, I.L., Kramer, N.J., Lyu, Z., Tsai, M.K., Chen, L.C., Cai, H., Gitler, A.D., Engleman, E., et al. (2020). A versatile system to record cell-cell interactions. eLife 9, e61080. 10.7554/eLife.61080.

3. McKenna, A., and Gagnon, J.A. (2019). Recording development with single cell dynamic lineage tracing. Development 146, dev169730. 10.1242/dev.169730.

4. Sankaran, V.G., Weissman, J.S., and Zon, L.I. (2022). Cellular barcoding to decipher clonal dynamics in disease. Science 378, eabm5874. 10.1126/science.abm5874.

5. David, G., O’Keefe, R.T., and Cai, Y. (2023). Cellular Surveillance: DNA-Based Recording to Monitor and Memorize Biological Events. GEN Biotechnol. 2, 197–210. 10.1089/genbio.2023.0018.

6. Bhattarai-Kline, S., Lear, S.K., Fishman, C.B., Lopez, S.C., Lockshin, E.R., Schubert, M.G., Nivala, J., Church, G.M., and Shipman, S.L. (2022). Recording gene expression order in DNA by CRISPR addition of retron barcodes. Nature 608, 217–225. 10.1038/s41586-022-04994-6.

7. Chen, W., Choi, J., Li, X., Nathans, J.F., Martin, B., Yang, W., Hamazaki, N., Qiu, C., Lalanne, J.-B., Regalado, S., et al. (2024). Symbolic recording of signalling and cis-regulatory element activity to DNA. Nature 632, 1073–1081. 10.1038/s41586-024-07706-4.

8. Choi, J., Chen, W., Minkina, A., Chardon, F.M., Suiter, C.C., Regalado, S.G., Domcke, S., Hamazaki, N., Lee, C., Martin, B., et al. (2022). A time-resolved, multi-symbol molecular recorder via sequential genome editing. Nature 608, 98–107. 10.1038/s41586-022-04922-8.

9. Rodriques, S.G., Chen, L.M., Liu, S., Zhong, E.D., Scherrer, J.R., Boyden, E.S., and Chen, F. (2021). RNA timestamps identify the age of single molecules in RNA sequencing. Nat. Biotechnol. 39, 320–325. 10.1038/s41587-020-0704-z.

10. Liu, Y., Huang, K., and Chen, W. (2024). Resolving cellular dynamics using single-cell temporal transcriptomics. Curr. Opin. Biotechnol. 85, 103060. 10.1016/j.copbio.2023.103060.

11. Askary, A., Chen, W., Choi, J., Du, L.Y., Elowitz, M.B., Gagnon, J.A., Schier, A.F., Seidel, S., Shendure, J., Stadler, T., et al. (2025). The lives of cells, recorded. Nat. Rev. Genet. 26, 203–222. 10.1038/s41576-024-00788-w.

12. Zhang, Y., Rózsa, M., Liang, Y., Bushey, D., Wei, Z., Zheng, J., Reep, D., Broussard, G.J., Tsang, A., Tsegaye, G., et al. (2023). Fast and sensitive GCaMP calcium indicators for imaging neural populations. Nature 615, 884–891. 10.1038/s41586-023-05828-9.

13. Piatkevich, K.D., Bensussen, S., Tseng, H., Shroff, S.N., Lopez-Huerta, V.G., Park, D., Jung, E.E., Shemesh, O.A., Straub, C., Gritton, H.J., et al. (2019). Population imaging of neural activity in awake behaving mice. Nature 574, 413–417. 10.1038/s41586-019-1641-1.

14. Su, Q., Zhang, J., Lin, W., Zhang, J.-F., Newton, A.C., Mehta, S., Yang, J., and Zhang, J. (2024). Sensitive fluorescent biosensor reveals differential subcellular regulation of PKC. Nat. Chem. Biol., 1–11. 10.1038/s41589-024-01758-3.

15. Frei, M.S., Mehta, S., and Zhang, J. (2024). Next-generation genetically encoded fluorescent biosensors illuminate cell signaling and metabolism. Annu. Rev. Biophys. 53, 275–297. 10.1146/annurev-biophys-030722-021359.

16. Yang, J.-M., Chi, W.-Y., Liang, J., Takayanagi, S., Iglesias, P.A., and Huang, C.-H. (2021). Deciphering cell signaling networks with massively multiplexed biosensor barcoding. Cell 184, 6193–6206.e14. 10.1016/j.cell.2021.11.005.

17. Terai, T., and Campbell, R.E. (2022). Barcodes, co-cultures, and deep learning take genetically encoded biosensor multiplexing to the nth degree. Mol. Cell 82, 239–240. 10.1016/j.molcel.2021.12.017.

18. Sadoine, M., Ishikawa, Y., Kleist, T.J., Wudick, M.M., Nakamura, M., Grossmann, G., Frommer, W.B., and Ho, C.-H. (2021). Designs, applications, and limitations of genetically encoded fluorescent sensors to explore plant biology. Plant Physiol. 187, 485–503. 10.1093/plphys/kiab353.

19. Keyes, J., Mehta, S., and Zhang, J. (2021). Strategies for Multiplexed Biosensor Imaging to Study Intracellular Signaling Networks. In Multiplexed Imaging: Methods and Protocols, E. Zamir, ed. (Springer US), pp. 1–20. 10.1007/978-1-0716-1593-5_1.

20. Ni, Q., Mehta, S., and Zhang, J. (2018). Live-cell imaging of cell signaling using genetically encoded fluorescent reporters. FEBS J. 285, 203–219. 10.1111/febs.14134.

21. Zaver, S.A., Johnson, C.J., Berndt, A., and Simpson, C.L. (2023). Live Imaging with Genetically Encoded Physiologic Sensors and Optogenetic Tools. J. Invest. Dermatol. 143, 353–361.e4. 10.1016/j.jid.2022.12.002.

22. Huppertz, M.-C., Wilhelm, J., Grenier, V., Schneider, M.W., Falt, T., Porzberg, N., Hausmann, D., Hoffmann, D.C., Hai, L., Tarnawski, M., et al. (2024). Recording physiological history of cells with chemical labeling. Science 383, 890–897. 10.1126/science.adg0812.

23. Linghu, C., An, B., Shpokayte, M., Celiker, O.T., Shmoel, N., Zhang, C., Park, W.M., Ramirez, S., and Boyden, E.S. (2021). Recording of cellular physiological histories along optically readable self-assembling protein chains (Synthetic Biology) 10.1101/2021.10.13.464006.

24. Lin, D., Li, X., Moult, E., Park, P., Tang, B., Shen, H., Grimm, J.B., Falco, N., Jia, B.Z., Baker, D., et al. (2023). Time-tagged ticker tapes for intracellular recordings. Nat. Biotechnol. 41, 631– 639. 10.1038/s41587-022-01524-7.

25. Garcia-Seisdedos, H., Empereur-Mot, C., Elad, N., and Levy, E.D. (2017). Proteins evolve on the edge of supramolecular self-assembly. Nature 548, 244–247. 10.1038/nature23320.

26. Baskaran, Y., Ang, K.C., Anekal, P.V., Chan, W.L., Grimes, J.M., Manser, E., and Robinson, R.C. (2015). An in cellulo-derived structure of PAK4 in complex with its inhibitor Inka1. Nat. Commun. 6, 8681. 10.1038/ncomms9681.

27. Hirano, M., Ando, R., Shimozono, S., Sugiyama, M., Takeda, N., Kurokawa, H., Deguchi, R., Endo, K., Haga, K., Takai-Todaka, R., et al. (2022). A highly photostable and bright green fluorescent protein. Nat. Biotechnol. 40, 1132–1142. 10.1038/s41587-022-01278-2.

28. Loew, R., Heinz, N., Hampf, M., Bujard, H., and Gossen, M. (2010). Improved Tet-responsive promoters with minimized background expression. BMC Biotechnol. 10, 81. 10.1186/1472-6750-10-81.

29. Koberstein, J.N., Stewart, M.L., Smith, C.B., Tarasov, A.I., Ashcroft, F.M., Stork, P.J.S., and Goodman, R.H. (2022). Monitoring glycolytic dynamics in single cells using a fluorescent biosensor for fructose 1,6-bisphosphate. Proc. Natl. Acad. Sci. 119, e2204407119. 10.1073/pnas.2204407119.

30. Dong, C., Zheng, Y., Long-Iyer, K., Wright, E.C., Li, Y., and Tian, L. (2022). Fluorescence Imaging of Neural Activity, Neurochemical Dynamics, and Drug-Specific Receptor Conformation with Genetically Encoded Sensors. Annu. Rev. Neurosci. 45, 273–294. 10.1146/annurev-neuro-110520-031137.

31. Mullick, A., Xu, Y., Warren, R., Koutroumanis, M., Guilbault, C., Broussau, S., Malenfant, F., Bourget, L., Lamoureux, L., Lo, R., et al. (2006). The cumate gene-switch: a system for regulated expression in mammalian cells. BMC Biotechnol. 6, 43. 10.1186/1472-6750-6-43.

32. Gammie, S.C., and Nelson, R.J. (2001). cFOS and pCREB activation and maternal aggression in mice. Brain Res. 898, 232–241. 10.1016/S0006-8993(01)02189-8.

33. Oh, J., Rhee, H.J., Kim, S.-W., Kim, S.B., You, H.-J., Kim, J.H., and Na, D.S. (2000). Annexin-I inhibits PMA-induced c-*fos* SRE activation by suppressing cytosolic phospholipase A2 signal. FEBS Lett. 477, 244–248. 10.1016/S0014-5793(00)01812-3.

34. Morgan, J.I., Cohen, D.R., Hempstead, J.L., and Curran, T. (1987). Mapping Patterns of c-fos Expression in the Central Nervous System After Seizure. Science 237, 192–197. 10.1126/science.3037702.

35. Cai, G., Lu, Y., Chen, J., Yang, D., Yan, R., Ren, M., He, S., Wu, S., and Zhao, Y. (2022). Brain-wide mapping of c-Fos expression with fluorescence micro-optical sectioning tomography in a chronic sleep deprivation mouse model. Neurobiol. Stress 20, 100478. 10.1016/j.ynstr.2022.100478.

36. Hodge, D.R., Hurt, E.M., and Farrar, W.L. (2005). The role of IL-6 and STAT3 in inflammation and cancer. Eur. J. Cancer 41, 2502–2512. 10.1016/j.ejca.2005.08.016.

37. Hogan, P.G., Chen, L., Nardone, J., and Rao, A. (2003). Transcriptional regulation by calcium, calcineurin, and NFAT. Genes Dev. 17, 2205–2232. 10.1101/gad.1102703.

38. Mancini, M., and Toker, A. (2009). NFAT proteins: emerging roles in cancer progression. Nat. Rev. Cancer 9, 810–820. 10.1038/nrc2735.

39. Macian, F. (2005). NFAT proteins: key regulators of T-cell development and function. Nat. Rev. Immunol. 5, 472–484. 10.1038/nri1632.

40. Li, Q., and Verma, I.M. (2002). NF-κB regulation in the immune system. Nat. Rev. Immunol. 2, 725–734. 10.1038/nri910.

41. Liu, T., Zhang, L., Joo, D., and Sun, S.-C. (2017). NF-κB signaling in inflammation. Signal Transduct. Target. Ther. 2, 1–9. 10.1038/sigtrans.2017.23.

42. Mayr, B., and Montminy, M. (2001). Transcriptional regulation by the phosphorylation-dependent factor CREB. Nat. Rev. Mol. Cell Biol. 2, 599–609. 10.1038/35085068.

43. Sassone-Corsi, P. (2012). The Cyclic AMP Pathway. Cold Spring Harb. Perspect. Biol. 4, a011148. 10.1101/cshperspect.a011148.

44. Scheller, L., Strittmatter, T., Fuchs, D., Bojar, D., and Fussenegger, M. (2018). Generalized extracellular molecule sensor platform for programming cellular behavior. Nat. Chem. Biol. 14, 723–729. 10.1038/s41589-018-0046-z.

45. Ye, H., Baba, M.D.-E., Peng, R.-W., and Fussenegger, M. (2011). A Synthetic Optogenetic Transcription Device Enhances Blood-Glucose Homeostasis in Mice. Science 332, 1565–1568. 10.1126/science.1203535.

46. Saxena, P., Heng, B.C., Bai, P., Folcher, M., Zulewski, H., and Fussenegger, M. (2016). A programmable synthetic lineage-control network that differentiates human IPSCs into glucose-sensitive insulin-secreting beta-like cells. Nat. Commun. 7, 11247. 10.1038/ncomms11247.

47. Shao, J., Qiu, X., Zhang, L., Li, S., Xue, S., Si, Y., Li, Y., Jiang, J., Wu, Y., Xiong, Q., et al. (2024). Multi-layered computational gene networks by engineered tristate logics. Cell 187, 5064–5080.e14. 10.1016/j.cell.2024.07.001.

48. Tu, Y.-C., Huang, D.-Y., Shiah, S.-G., Wang, J.-S., and Lin, W.-W. (2013). Regulation of c-Fos Gene Expression by NF-κB: A p65 Homodimer Binding Site in Mouse Embryonic Fibroblasts but Not Human HEK293 Cells. PLOS ONE 8, e84062. 10.1371/journal.pone.0084062.

49. Maggirwar, S.B., Sarmiere, P.D., Dewhurst, S., and Freeman, R.S. (1998). Nerve Growth Factor-Dependent Activation of NF-κB Contributes to Survival of Sympathetic Neurons. J. Neurosci. 18, 10356–10365. 10.1523/JNEUROSCI.18-24-10356.1998.

50. Angel, P., and Karin, M. (1991). The role of jun, fos and the AP-1 complex in cell-proliferation and transformation. Biochim. Biophys. Acta 1072, 129–157. 10.1016/0304-419x(91)90011-9.

51. Sakurai, H., Suzuki, S., Kawasaki, N., Nakano, H., Okazaki, T., Chino, A., Doi, T., and Saiki, I. (2003). Tumor Necrosis Factor-α-induced IKK Phosphorylation of NF-κB p65 on Serine 536 Is Mediated through the TRAF2, TRAF5, and TAK1 Signaling Pathway*. J. Biol. Chem. 278, 36916–36923. 10.1074/jbc.M301598200.

52. Zhang, Y., and Hu, W. (2012). NFκB signaling regulates embryonic and adult neurogenesis. Front. Biol. 7, 277–291. 10.1007/s11515-012-1233-z.

53. Charital, Y.M., van Haasteren, G., Massiha, A., Schlegel, W., and Fujita, T. (2009). A functional NF-kappaB enhancer element in the first intron contributes to the control of c-fos transcription. Gene 430, 116–122. 10.1016/j.gene.2008.10.014.

54. Tavares, L.P., Negreiros-Lima, G.L., Lima, K.M., E Silva, P.M.R., Pinho, V., Teixeira, M.M., and Sousa, L.P. (2020). Blame the signaling: Role of cAMP for the resolution of inflammation. Pharmacol. Res. 159, 105030. 10.1016/j.phrs.2020.105030.

55. Diehl, A.M., Yang, S.Q., Wolfgang, D., and Wand, G. (1992). Differential expression of guanine nucleotide-binding proteins enhances cAMP synthesis in regenerating rat liver. J. Clin. Invest. 89, 1706–1712. 10.1172/JCI115771.

56. Bai, P., Ye, H., Xie, M., Saxena, P., Zulewski, H., Hamri, G.C.-E., Djonov, V., and Fussenegger, M. (2016). A synthetic biology-based device prevents liver injury in mice. J. Hepatol. 65, 84–94. 10.1016/j.jhep.2016.03.020.

57. Li, L., Cui, L., Lin, P., Liu, Z., Bao, S., Ma, X., Nan, H., Zhu, W., Cen, J., Mao, Y., et al. (2023). Kupffer-cell-derived IL-6 is repurposed for hepatocyte dedifferentiation via activating progenitor genes from injury-specific enhancers. Cell Stem Cell 30, 283–299.e9. 10.1016/j.stem.2023.01.009.

58. Park, J., Zhao, Y., Zhang, F., Zhang, S., Kwong, A.C., Zhang, Y., Hoffmann, H.-H., Bushweller, L., Wu, X., Ashbrook, A.W., et al. (2023). IL-6/STAT3 axis dictates the PNPLA3-mediated susceptibility to non-alcoholic fatty liver disease. J. Hepatol. 78, 45–56. 10.1016/j.jhep.2022.08.022.

59. Zhou, M., Mok, M.T.S., Sun, H., Chan, A.W., Huang, Y., Cheng, A.S.L., and Xu, G. (2017). The anti-diabetic drug exenatide, a glucagon-like peptide-1 receptor agonist, counteracts hepatocarcinogenesis through cAMP–PKA–EGFR–STAT3 axis. Oncogene 36, 4135–4149. 10.1038/onc.2017.38.

60. Wahlang, B., McClain, C., Barve, S., and Gobejishvili, L. (2018). Role of cAMP and phosphodiesterase signaling in liver health and disease. Cell. Signal. 49, 105–115. 10.1016/j.cellsig.2018.06.005.

61. Mátés, L., Chuah, M.K.L., Belay, E., Jerchow, B., Manoj, N., Acosta-Sanchez, A., Grzela, D.P., Schmitt, A., Becker, K., Matrai, J., et al. (2009). Molecular evolution of a novel hyperactive Sleeping Beauty transposase enables robust stable gene transfer in vertebrates. Nat. Genet. 41, 753–761. 10.1038/ng.343.

62. Johnson, D.E., O’Keefe, R.A., and Grandis, J.R. (2018). Targeting the IL-6/JAK/STAT3 signalling axis in cancer. Nat. Rev. Clin. Oncol. 15, 234–248. 10.1038/nrclinonc.2018.8.

63. Billing, U., Jetka, T., Nortmann, L., Wundrack, N., Komorowski, M., Waldherr, S., Schaper, F., and Dittrich, A. (2019). Robustness and Information Transfer within IL-6-induced JAK/STAT Signalling. Commun. Biol. 2, 1–14. 10.1038/s42003-018-0259-4.

64. Kellogg, R.A., Tian, C., Lipniacki, T., Quake, S.R., and Tay, S. (2015). Digital signaling decouples activation probability and population heterogeneity. eLife 4, e08931. 10.7554/eLife.08931.

65. Kingeter, L.M., Paul, S., Maynard, S.K., Cartwright, N.G., and Schaefer, B.C. (2010). Cutting Edge: TCR Ligation Triggers Digital Activation of NF-κB. J. Immunol. 185, 4520–4524. 10.4049/jimmunol.1001051.

66. Goul, C., Peruzzo, R., and Zoncu, R. (2023). The molecular basis of nutrient sensing and signalling by mTORC1 in metabolism regulation and disease. Nat. Rev. Mol. Cell Biol. 24, 857– 875. 10.1038/s41580-023-00641-8.

67. Joshi, J.N., Lerner, A.D., Scallo, F., Grumet, A.N., Matteson, P., Millonig, J.H., and Valvezan, A.J. (2024). mTORC1 activity oscillates throughout the cell cycle, promoting mitotic entry and differentially influencing autophagy induction. Cell Rep. 43, 114543. 10.1016/j.celrep.2024.114543.

68. Shimano, H., and Sato, R. (2017). SREBP-regulated lipid metabolism: convergent physiology — divergent pathophysiology. Nat. Rev. Endocrinol. 13, 710–730. 10.1038/nrendo.2017.91.

69. Amemiya-Kudo, M., Shimano, H., Hasty, A.H., Yahagi, N., Yoshikawa, T., Matsuzaka, T., Okazaki, H., Tamura, Y., Iizuka, Y., Ohashi, K., et al. (2002). Transcriptional activities of nuclear SREBP-1a, –1c, and –2 to different target promoters of lipogenic and cholesterogenic genes. J. Lipid Res. 43, 1220–1235.

70. Yokoyama, C., Wang, X., Briggs, M.R., Admon, A., Wu, J., Hua, X., Goldstein, J.L., and Brown, M.S. (1993). SREBP-1, a basic-helix-loop-helix-leucine zipper protein that controls transcription of the low density lipoprotein receptor gene. Cell 75, 187–197. 10.1016/S0092-8674(05)80095-9.

71. Latasa, M.-J., Griffin, M.J., Moon, Y.S., Kang, C., and Sul, H.S. (2003). Occupancy and function of the –150 sterol regulatory element and –65 E-box in nutritional regulation of the fatty acid synthase gene in living animals. Mol. Cell. Biol. 23, 5896–5907. 10.1128/MCB.23.16.5896-5907.2003.

72. Liu, Q., Chang, J.W., Wang, J., Kang, S.A., Thoreen, C.C., Markhard, A., Hur, W., Zhang, J., Sim, T., Sabatini, D.M., et al. (2010). Discovery of 1-(4-(4-Propionylpiperazin-1-yl)-3-(trifluoromethyl)phenyl)-9-(quinolin-3-yl)benzo[h][1,6]naphthyridin-2(1H)-one as a Highly Potent, Selective Mammalian Target of Rapamycin (mTOR) Inhibitor for the Treatment of Cancer. J. Med. Chem. 53, 7146–7155. 10.1021/jm101144f.

73. S, Z., C, C., S, W., F, J., and Y, X. (2016). MHY1485 activates mTOR and protects osteoblasts from dexamethasone. Biochem. Biophys. Res. Commun. 481. 10.1016/j.bbrc.2016.10.104.

74. Subach, O.M., Vlaskina, A.V., Agapova, Y.K., Nikolaeva, A.Y., Anokhin, K.V., Piatkevich, K.D., Patrushev, M.V., Boyko, K.M., and Subach, F.V. (2023). Blue-to-Red TagFT, mTagFT, mTsFT, and Green-to-FarRed mNeptusFT2 Proteins, Genetically Encoded True and Tandem Fluorescent Timers. Int. J. Mol. Sci. 24, 3279. 10.3390/ijms24043279.

75. Hwang, J.-R., Byeon, Y., Kim, D., and Park, S.-G. (2020). Recent insights of T cell receptor-mediated signaling pathways for T cell activation and development. Exp. Mol. Med. 52, 750– 761. 10.1038/s12276-020-0435-8.

76. Manger, B., Hardy, K.J., Weiss, A., and Stobo, J.D. (1986). Differential effect of cyclosporin A on activation signaling in human T cell lines. J. Clin. Invest. 77, 1501–1506. 10.1172/JCI112464.

77. Sanchez-Vega, F., Mina, M., Armenia, J., Chatila, W.K., Luna, A., La, K.C., Dimitriadoy, S., Liu, D.L., Kantheti, H.S., Saghafinia, S., et al. (2018). Oncogenic Signaling Pathways in The Cancer Genome Atlas. Cell 173, 321–337.e10. 10.1016/j.cell.2018.03.035.

78. Kochen, M.A., Andrews, S.S., Wiley, H.S., Feng, S., and Sauro, H.M. (2022). Dynamics and Sensitivity of Signaling Pathways. Curr. Pathobiol. Rep. 10, 11–22. 10.1007/s40139-022-00230-y.

79. Valls, P.O., and Esposito, A. (2022). Signalling dynamics, cell decisions, and homeostatic control in health and disease. Curr. Opin. Cell Biol. 75, 102066. 10.1016/j.ceb.2022.01.011.

80. Shah, K., Al-Haidari, A., Sun, J., and Kazi, J.U. (2021). T cell receptor (TCR) signaling in health and disease. Signal Transduct. Target. Ther. 6, 1–26. 10.1038/s41392-021-00823-w.

81. Huang, W., Lin, W., Chen, B., Zhang, J., Gao, P., Fan, Y., Lin, Y., and Wei, P. (2023). NFAT and NF-κB dynamically co-regulate TCR and CAR signaling responses in human T cells. Cell Rep. 42, 112663. 10.1016/j.celrep.2023.112663.

82. Barykina, N.V., Karasev, M.M., Verkhusha, V.V., and Shcherbakova, D.M. (2022). Technologies for large-scale mapping of functional neural circuits active during a user-defined time window. Prog. Neurobiol. 216, 102290. 10.1016/j.pneurobio.2022.102290.

83. Pang, B., Wu, X., Chen, H., Yan, Y., Du, Z., Yu, Z., Yang, X., Wang, W., and Lu, K. (2023). Exploring the memory: existing activity-dependent tools to tag and manipulate engram cells. Front. Cell. Neurosci. 17, 1279032. 10.3389/fncel.2023.1279032.

84. Beaudoin, G.M.J., Lee, S.-H., Singh, D., Yuan, Y., Ng, Y.-G., Reichardt, L.F., and Arikkath, J. (2012). Culturing pyramidal neurons from the early postnatal mouse hippocampus and cortex. Nat. Protoc. 7, 1741–1754. 10.1038/nprot.2012.099.

85. Jiang, M., and Chen, G. (2006). High Ca2+-phosphate transfection efficiency in low-density neuronal cultures. Nat. Protoc. 1, 695–700. 10.1038/nprot.2006.86.

