## Supplementary Materials for "Fluorescent protein-based ticker tapes for multiplexed recordings of transcriptional histories in single cells in culture and in vivo"

**Contents**

Supplementary Movie 1

Supplementary Table 1

Supplementary Figures 1 – 39

**Supplementary Movie 1 Captions**

Live imaging of fiber growth commenced 6 hours post-transfection and continued for 48 hours, with images captured at 15-minute intervals under Sartorius Incucyte SX5 live cell analysis system. HEK293T cells were transfected with pCMV-iPAK4 (90%, w/w) and pCMV-mGreenLantern-iPAK4 (10%, w/w). Scale bar, 50 μm.

**Supplementary Table 1**. A comparison of FPTT with iPAK4-HaloTag system

| Characteristics | iPAK4-HaloTag system | FPTT |
|---|---|---|
| Temporal order of events | Yes | Yes |
| Spatial resolution | Single cell | Single cell |
| Absolute of time events | Yes | Yes |
| Multiplexed ability | Only 1 activity | **7 activities, 4 activities in multiplexed** |
| Synchronous, spontaneous, or asynchronous events | Synchronous events | **Synchronous, spontaneous, and asynchronous events** |
| Cell state disruption | Low, but fixation at the endpoint | Low, **do not need to fix cells** |
| Analysis time | Endpoint, need multiple steps to wash out dyes | Endpoint and real-time, **Unnecessary for medium manipulation** |
| Transient signal & long-term exposure | Hours | **Both transient (minutes to hours) and long-term (days)** |
| Temporal resolution | **2-hour** | 3-hour |
| Duration | 24 hours | **8 days** |
| *In vivo* application | Not demonstrated | **Yes (mouse liver and brain)** |
| Cell types | HEK293T, primary neurons | **HEK293T, HeLa, primary neurons, Jurat cells** |

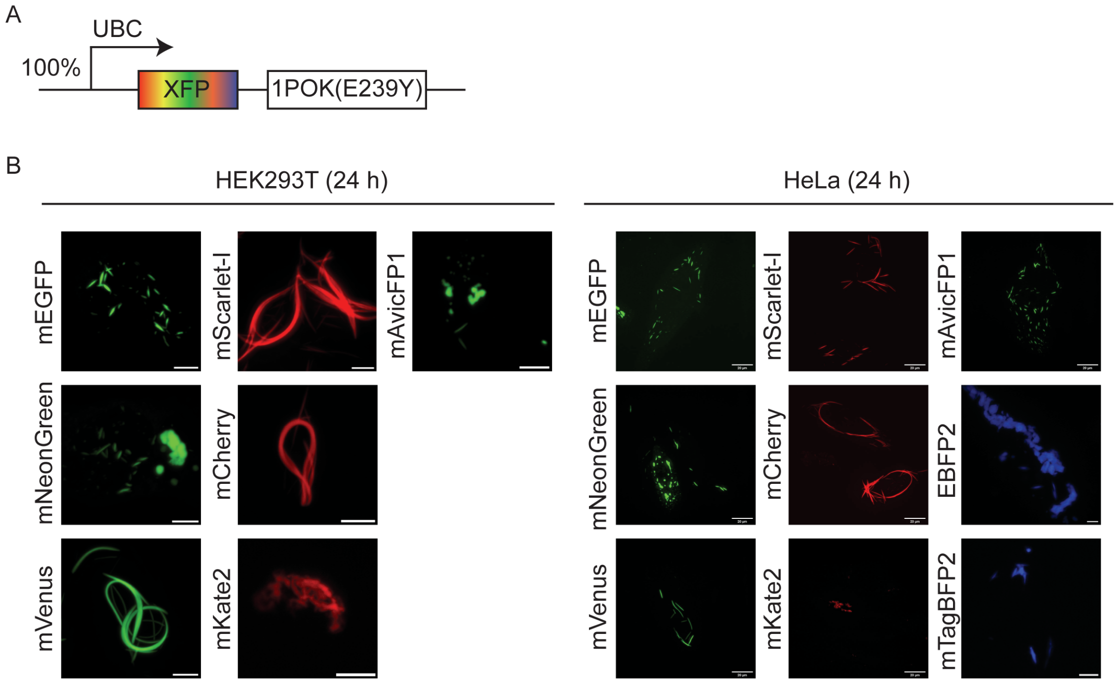


**Supplementary Figure 1.** Localization of the 1POK fusions with various fluorescent proteins in mammalian cell cultures.

**A**, Genetic constructs used for expression of fluorescent ticker tape in HEK293T and HeLa cells. The percentage indicates the ratio of the plasmids used for co-transfection (used here and throughout the rest of the figures). Here, XFP represents GFPs (mEGFP, mVenus, mNeonGreen, mAvicFP1) or RFPs (mScarlet-I, mCherry, mKate2) or BFPs (EBFP2, mTagBFP2), respectively. **B**, Left, Representative maximum intensity projection (MIP) confocal images of live HEK293T cells expressing the XFP-1POK fusions under the UBC promoter (n > 40 cells from 3 independent cultures each imaged 24 h post-transfection). The fusions with mEGFP, mScarlet-I, mCherry, and mVenus formed rod-shaped structures, and the fusions with mAvicFP1, mNeonGreen, and mKate2 formed irregular shape aggregates. Scale bars, 10 µm. Right, Representative confocal MIP images of live HeLa cells expressing the XFP-1POK fusions under the UBC promoter (n > 40 cells from 3 independent cultures each imaged 24 h post-transfection). The fusions with mEGFP, mScarlet-I, mAvicFP1, mCherry, and mVenus formed rod-shaped structures, and the fusions with mNeonGreen, mKate2, EBFP2, and mTagBFP2 formed irregular shape aggregates. Scale bars, 20 µm.

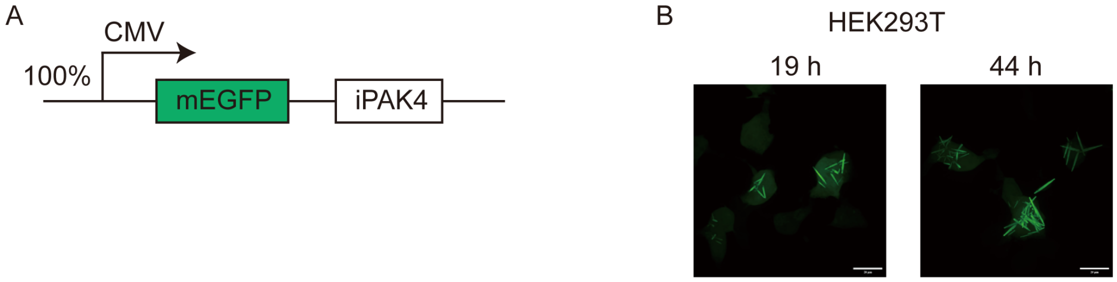


**Supplementary Figure 2.** Fiber structures by mEGFP-tagged iPAK4 in HEK293T cells.

**A**, Genetic constructs used for expression of fluorescent ticker tapes in HEK293T for experiments shown in panel b. **B**, Representative confocal MIP images of live HEK expressing only mEGFP-iPAK4 fusion in HEK293T cells (n > 20 cells from 3 independent cultures imaged 19 h and 44 h post-transfection). Scale bars, 20 μm.

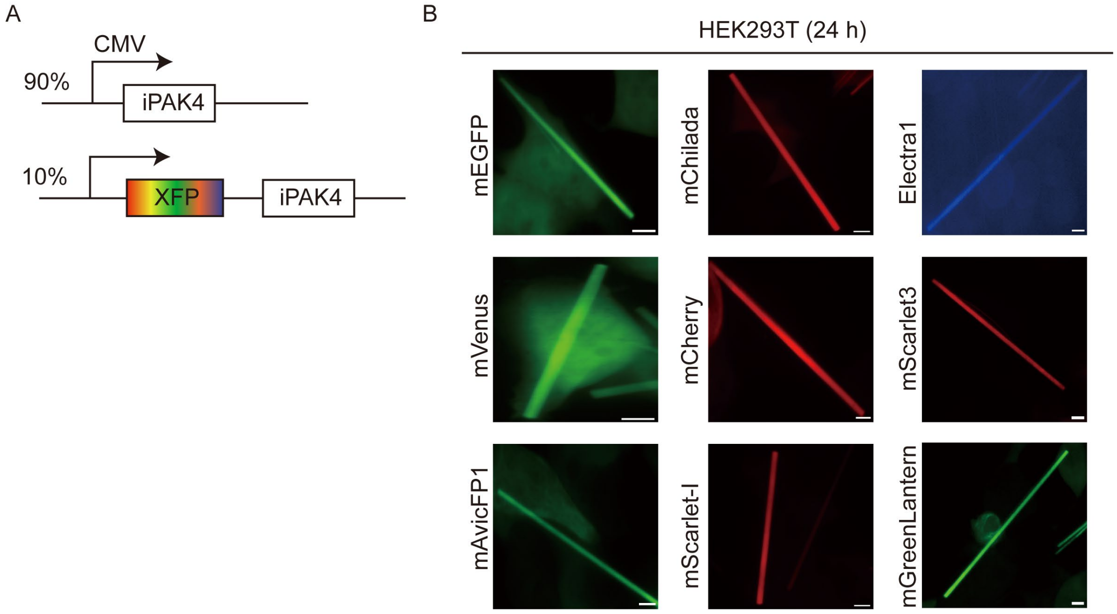


**Supplementary Figure 3.** Fiber structures by iPAK4 that contain 10% of monomeric FP-tags in HEK293T cells

**A**, Genetic constructs used for expression of fluorescent ticker tapes in HEK293T for experiments shown in panel b. Here, XFP represents GFPs (mEGFP, mVenus, mAvicFP1), RFPs (mScarlet-I, mCherry, mChilada), and BFPs (Electra1). **B**, Representative confocal MIP images of live HEK293T cells co-expressing iPAK4 with GFP-iPAK4, RFP-iPAK4, and BFP-iPAK4 ($n > 40$ cells from 3 independent cultures each imaged 24 h post-transfection). Scale bars, 5 µm

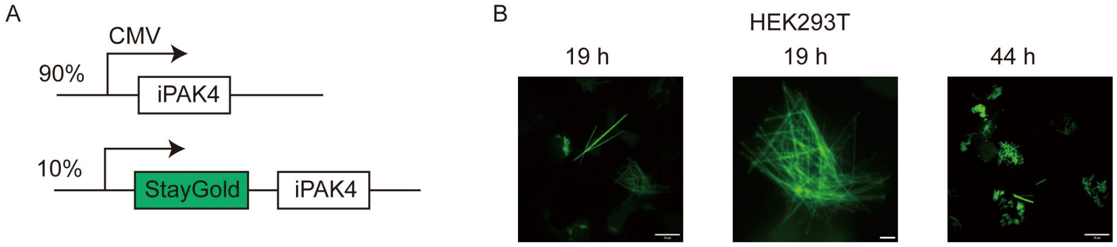


**Supplementary Figure 4.** Fiber structures by iPAK4 that contain 10% of dimeric FP-tags in HEK293T cells

**A**, Genetic constructs used for expression of fluorescent ticker tape in HEK293T for experiments shown in panel B. **B**, Representative confocal MIP images of live HEK cells co-expressing iPAK4 with dimeric StayGold-iPAK4 in HEK293T cells (n > 20 cells from 3 independent cultures imaged 19, 44 h post-transfection). Scale bars, 20 µm.

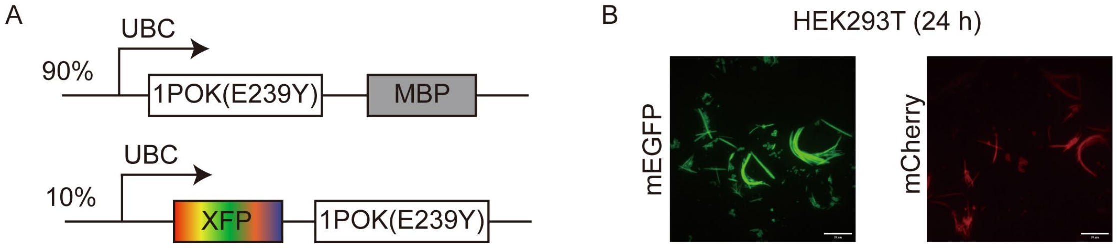


**Supplementary Figure 5.** Fiber structures by 1POK that contain 10% of monomeric FP-tags in HEK293T cells

**A**, Genetic constructs used for expression of fluorescent ticker tapes in HEK293T for experiments shown in panel B. Here, XFP represents mEGFP, and mCherry. MBP, maltose binding protein, was reasoned as a monomeric insulator for the linear growth of fibers. **B**, Representative maximum intensity projection confocal images of live HEK cells co-expressing 1POK-MBP with mEGFP-1POK (left panel) and mCherry-1POK (right panel) in HEK293T cells (n > 20 cells from 3 independent cultures each imaged 24 h post-transfection). Scale bars, 20 µm.

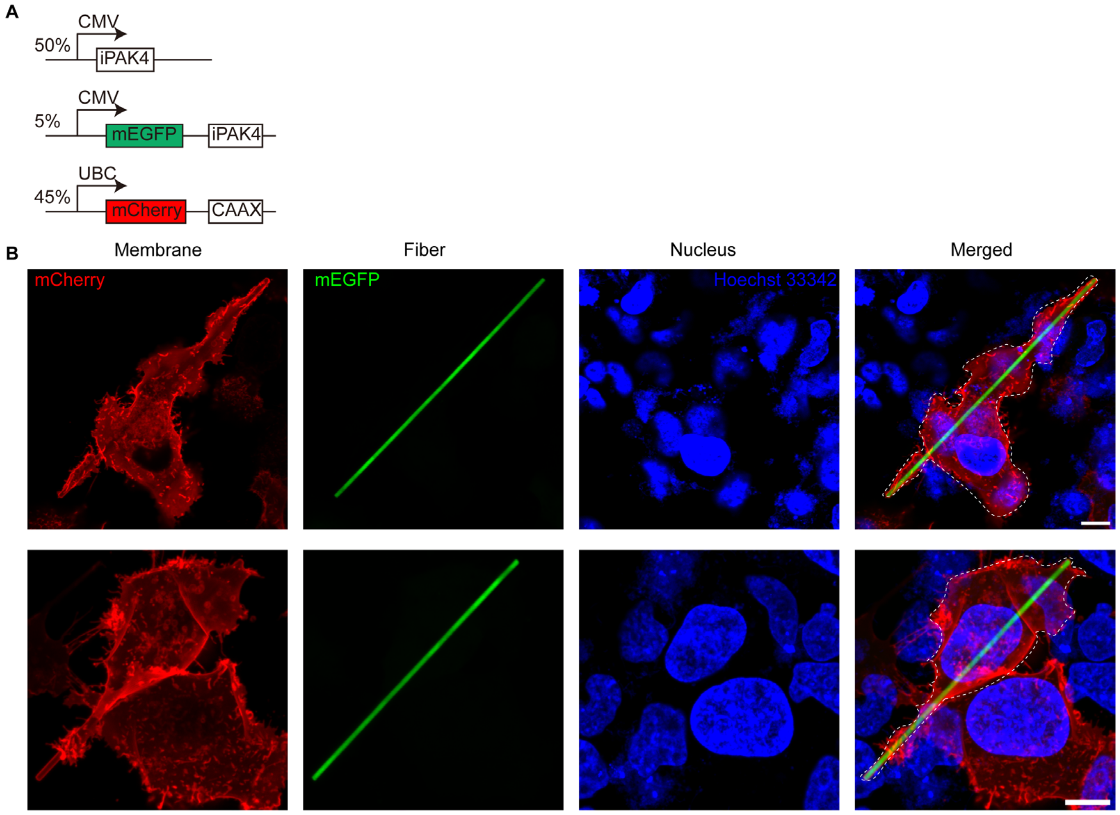


**Supplementary Figure 6**. Intracellular localization of iPAK4 fibers in HEK293T Cells

**A**, Schematic representation of the genetic cassette, which includes iPAK4, mEGFP-iPAK4, and a membrane-targeted mCherry-CAAX, all expressed in a defined ratio. **B**, Representative images showing the intracellular localization of fibers in HEK293T cells. Red indicates the mCherry-targeted membrane, green represents the mEGFP-fused iPAK4 fibers, and blue corresponds to the nucleus stained with Hoechst 33342. In cells where the fiber length exceeded the cell size, the membrane wrapped around the extended fiber. Scale bar, 10 μm.

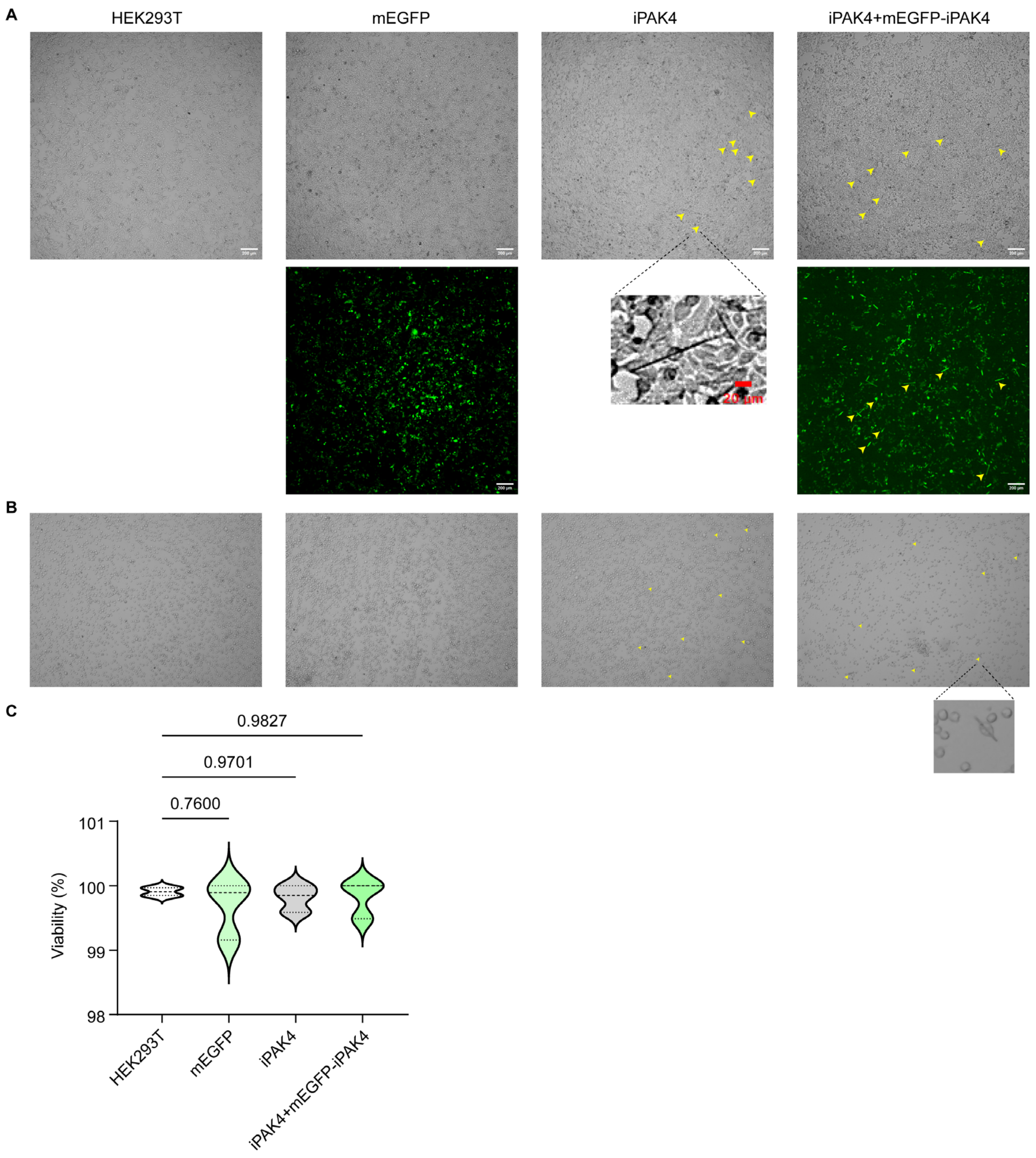


**Supplementary Figure 7**. Trypan Blue to assess effect of iPAK4 fibers on cell health.

**A**, HEK293T cells were transfected with the same amount of PEI and mEGFP as a control, along with iPAK4 and iPAK4: mEGFP-iPAK4 (9:1) for the experimental group. Images were captured 48 hours post-transfection, prior to trypsin digestion. Multiple fibers were observed in the experimental groups under bright field and GFP channel. **B**, Cell status following trypsin digestion showing that numerous fibers remained within the digested cells. **C**, Cell viability statistics presented as mean ± SD from three independent cultures. Statistical analysis was conducted using one-way ANOVA for *p* values.

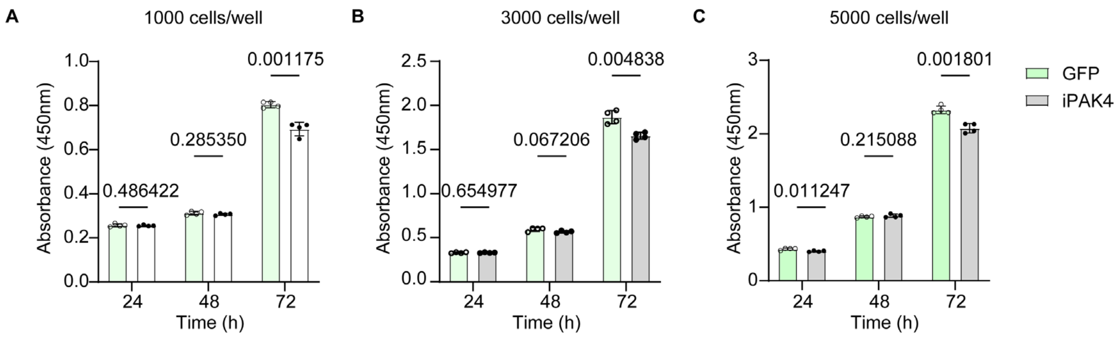


**Supplementary Figure 8**. Cell viability and proliferation assessment using CCK-8 assay.

**A**, HEK293T cells transfected with GFP or iPAK4, initially plated at 1000 cells, were assessed using the CCK-8 assay at 24, 48, and 72 hours following a 24-hour post-transfection period. **B**, HEK293T cells transfected with GFP or iPAK4 and initially plated at 3000 cells underwent CCK-8 assay analysis at 24, 48, and 72 hours after 24 hours post-transfection. **C**, HEK293T cells transfected with GFP or iPAK4, plated with a starting count of 5000 cells, were evaluated using the CCK-8 assay at 24, 48, and 72 hours following 24 hours post-transfection. Statistics are shown as mean ± SD from four independent cultures. Statistical significance was determined using multiple unpaired Student's *t*-tests for *p* values.

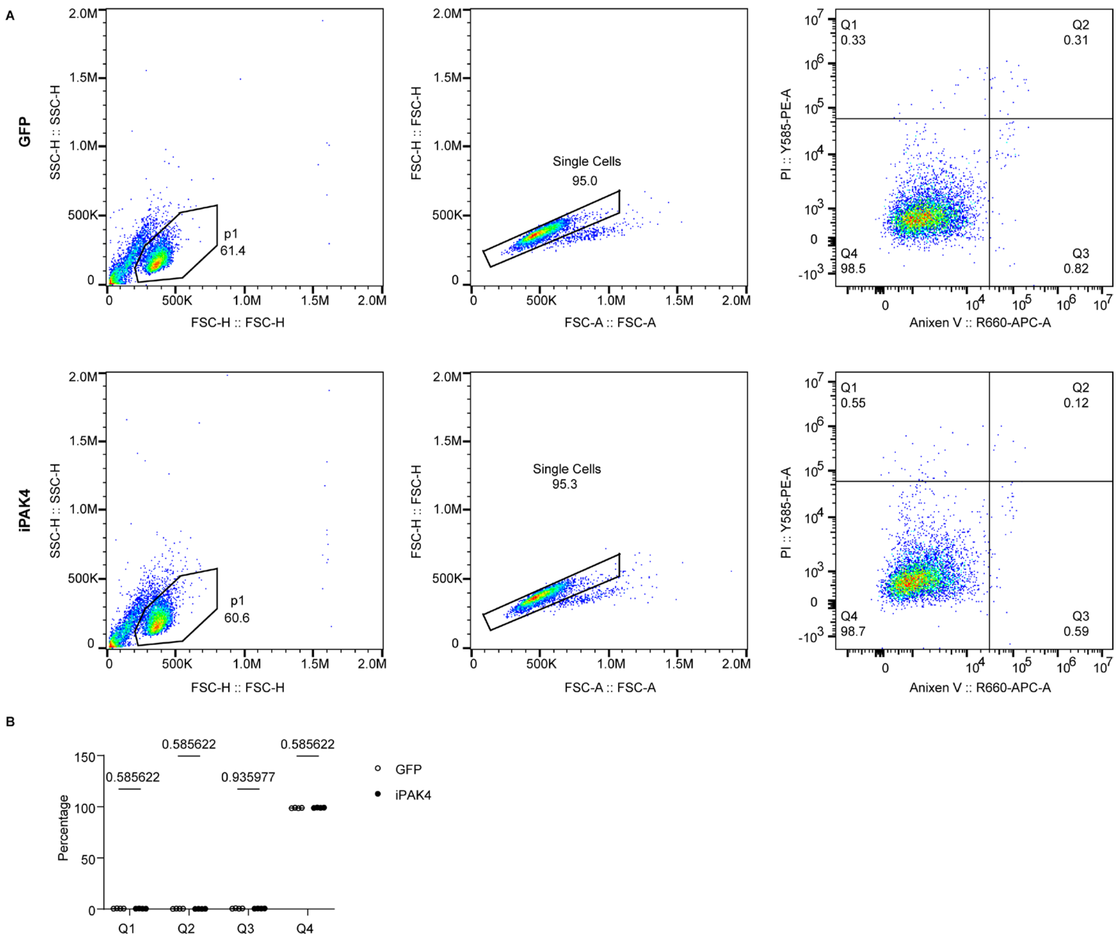


**Supplementary Figure 9**. Annexin V/PI Kit to assess effect of iPAK4 fibers on cell health

**A**, Gating strategy for analyzing the apoptosis cells with a live-dead kit using Annexin V and PI by flow cytometry. **B**, Percentage of cells with expressing iPAK4 fibers or GFP expression in each stage. Each black dot is from one independent transfection. Data are shown as the mean±SD of n = 4 independent experiments, $p$ values are calculated using two-way ANOVA. There is no statistically significance on the percentage of the apoptosis cells and live cells.

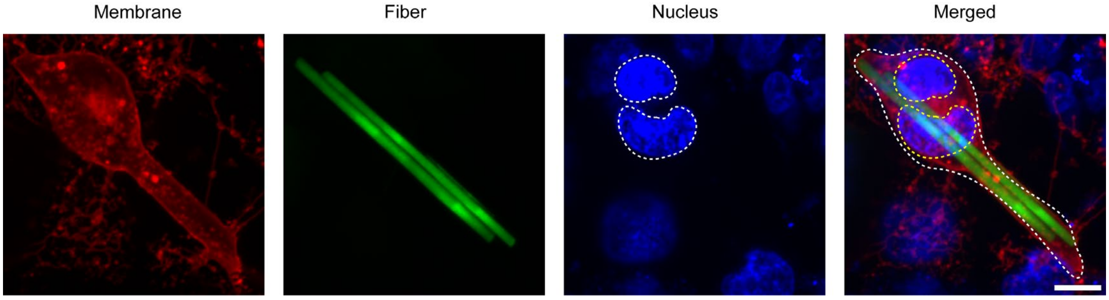


**Supplementary Figure 10**. Cell division in HEK293T cells containing iPAK4 fibers.

Representative snapshots showing mitosis and cell divisions of a HEK293T cell containing an iPAK4 fiber. Red indicates the mCherry-targeted membrane, green represents the mEGFP-fused iPAK4 fibers, and blue corresponds to the nucleus stained with Hoechst 33342. Scale bar, 10 µm.

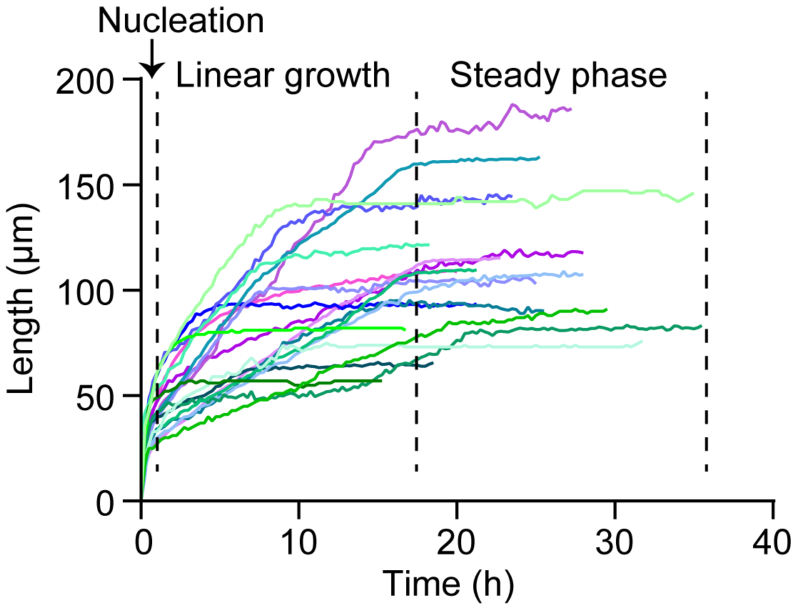


**Supplementary Figure 11**. Fiber growth profiles in HEK293T cells

Live imaging of fiber growth commenced 6 hours post-transfection and continued for 48 hours, with images captured at 15-minute intervals (see Supplementary Video 1). The starting point, denoted as time 0, is defined by the initial frame showing the onset of fiber growth. Typically, fibers experience a rapid nucleation phase within the first hour. This is followed by a nearly linear growth phase, with varying fiber lengths. Eventually, fibers progress into a steady growth phase, characterized by minimal growth.

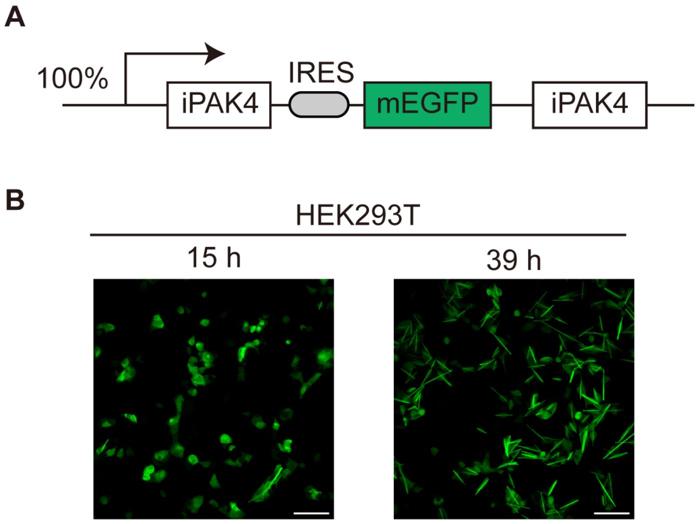


**Supplementary Figure 12**. Bicistronic Expression of FPTT with an IRES in HEK293T Cells

**A** Schematic illustration of the genetic cassette design, featuring iPAK4 and mEGFP-iPAK4 connected by an internal ribosome entry site (IRES). **B**, Representative wide-field images of HEK293T cells transfected with the FPTT construct described in panel A, captured at 15- and 39-hours post-transfection. Scale bar, 100 µm.

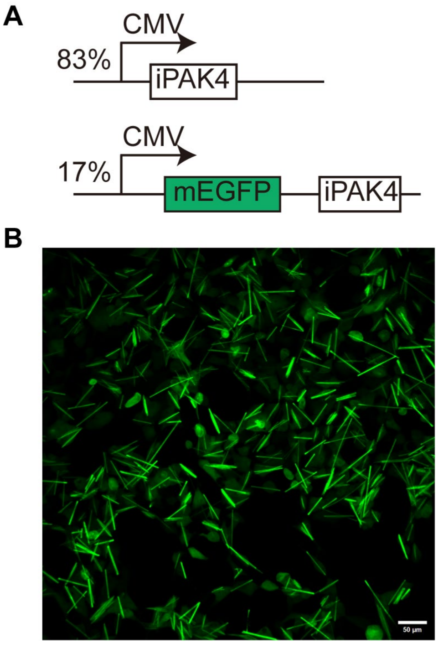


**Supplementary Figure 13**. Expression of FPTT at a 5:1 ratio in HEK293T cells

**A**, Schematic representation of the genetic cassette comprising iPAK4 and mEGFP-iPAK4 in an approximate 5:1 ratio. **B**, Representative images displaying a large view of HEK293T cells transfected with the designated FPTT ratio as described in **A** after 39 hours. Scale bar, 50 µm.

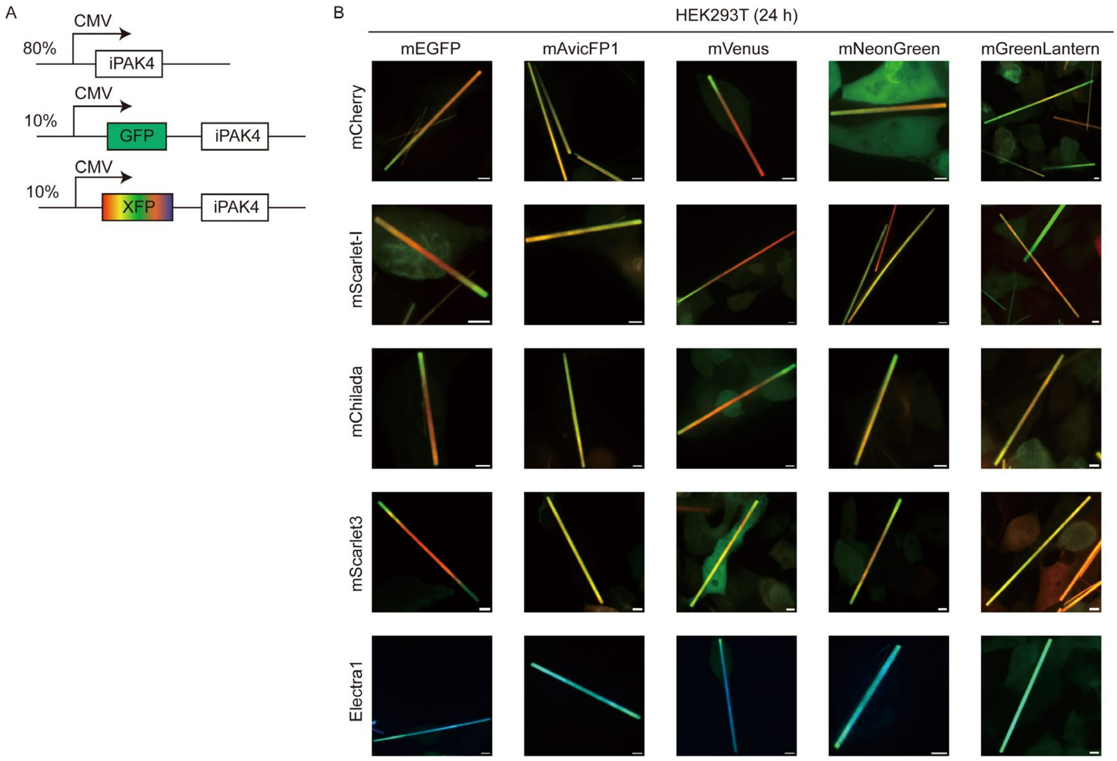


**Supplementary Figure 14.** Fiber structures by iPAK4 that contain multiple monomeric FP-tags in HEK293T cells

**A**, Genetic constructs used for expression of fluorescent ticker tapes in HEK293T for experiments shown in panel B. Here, GFP represents mEGFP, mAvicFP1, mVenus, mNeonGreen, mGreenLantern. XFP represents RFPs (mScarlet-I, mCherry, mChilada, mScarlet3), and BFP (Electra1). **B**, Representative confocal MIP images of live HEK293T cells co-expressing iPAK4 with GFP-iPAK4, RFP-iPAK4, and BFP-iPAK4 (n > 20 cells from 3 independent cultures each imaged 24 h post-transfection). Scale bars, 5 μm.

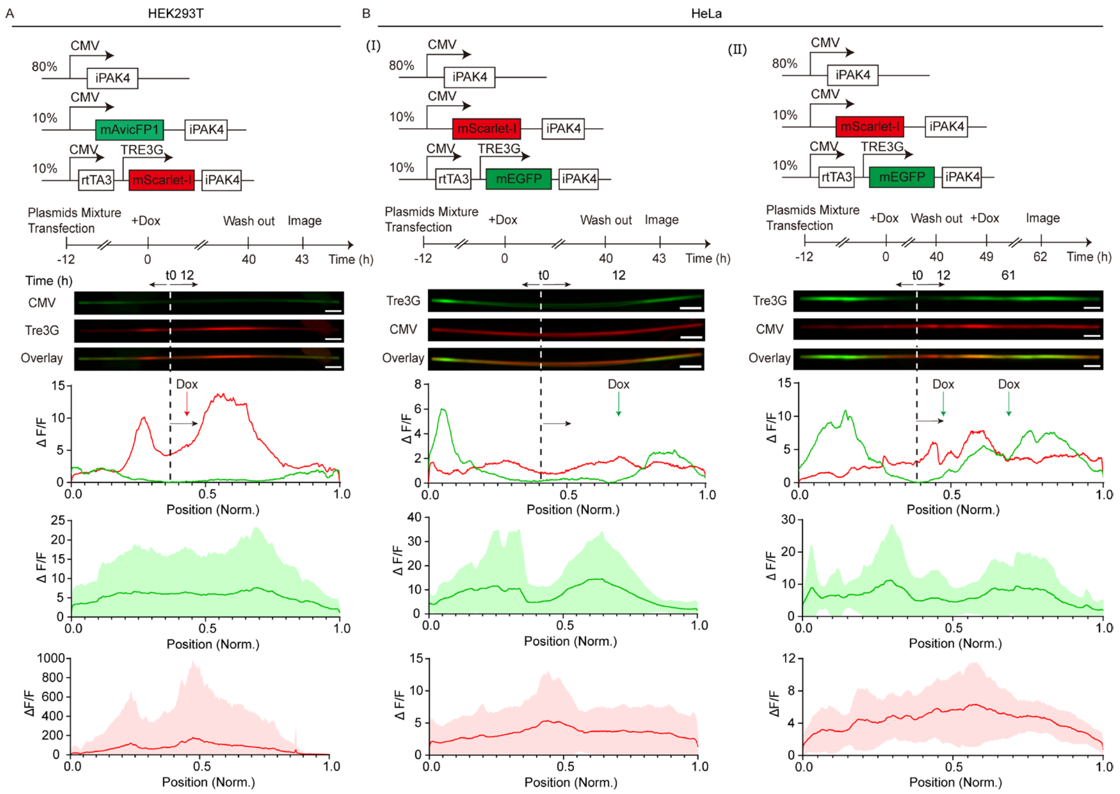


**Supplementary Figure 15.** Doxycycline-inducible Ticker tape-recording in HEK293T and HeLa cells

**A**, Genetic constructs used for expression of fluorescent ticker tapes. HEK-293T were co-transfected with a constitutive iPAK4 expression vector (80%, w/w), a constitutive mAvicFP1-iPAK4 expression vector producing green fluorescence (10%, w/w) and an rtTA-dependent expression vector for mScarlet-I-iPAK4 (10%, w/w) for doxycycline-inducible "inscription" of red fluorescence into mAvicFP1-containing iPAK4 fibers. Representative microscopic images were acquired at the experimental endpoint following one recurrent events of 10 ng/mL Doxycycline-stimulation. Averaged fluorescence intensity changes (ΔF/F) from n = 21 fibers (from 3 biological replicates) were monitored over the entire experimental timespan. t0 and dash lines, the start timepoints of fiber growth; black arrow, the direction of fiber growth; red arrow, the application of Dox. **B**, HeLa cells were co-transfected with a constitutive iPAK4 expression vector (80%, w/w), a constitutive mScarlet-I-iPAK4 expression vector (10%, w/w) and a rtTA-dependent mEGFP-iPAK4 expression vector (10%, w/w). Representative microscopic images were acquired at the experimental endpoint following (**I**) one or (**II**) two recurrent events of 10 ng/mL Doxycycline-stimulation. Averaged fluorescence intensity changes (ΔF/F) from n = 29 fibers in (I) and n = 11 fibers from 3 biological replicates in (II) were monitored over the entire experimental timespan. t0 and dash lines, the start timepoints of fiber growth; black arrow, the direction of fiber growth; green arrow, the application of Dox. Solid line, mean; shaded area, SD. Scale bars, 10 μm.

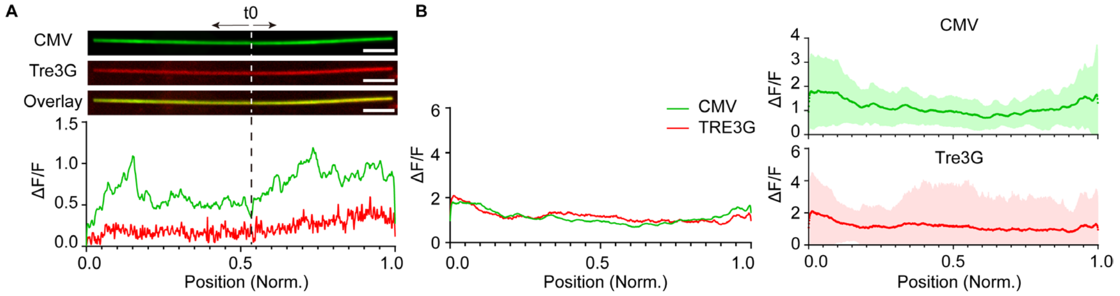


**Supplementary Figure 16**. Control experiment of Tet-ON FPTT without Dox stimulation. Related to Fig. 1B

**A**, Represented fiber and ΔF/F profile of same constructs in Fig. 1B without Dox treatments in same time window. t0 and dash lines, the start timepoints of fiber growth; black arrow, the direction of fiber growth. Scale bar, 10 µm. **B**, Averaged ΔF/F profile of same constructs in Fig. 1B without dox treatments in same time window. Red lines, mean of Tre3G fold changes; green line, mean of CMV fold changes; shadow area, SD. N = 23 fibers from two independent cultures.

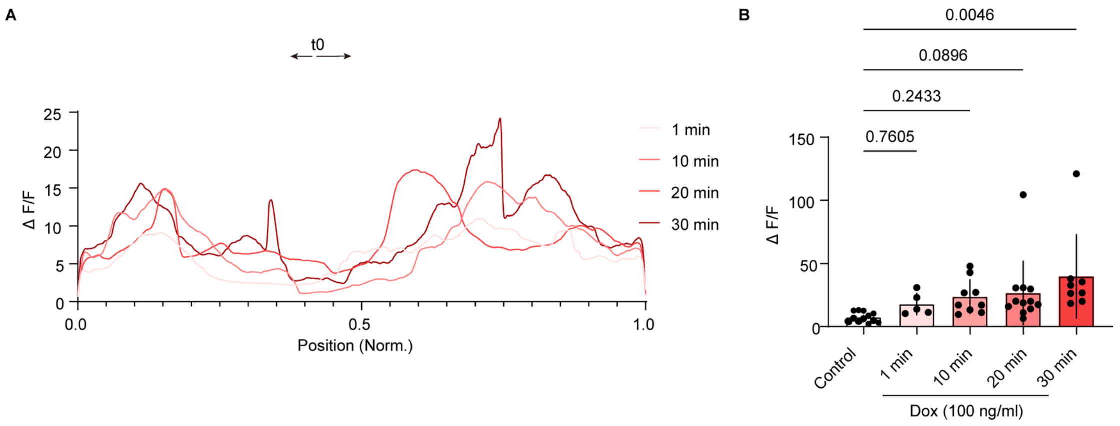


**Supplementary Figure 17**. Minutes scale of Dox-induced activity on FPTT. Related to Fig. 1F

**A**, Averaged profiles of FPTT treated with 100 ng/ml Dox with 1, 10, 20, and 30 minutes in Figure 1F. t0 and dash lines, the start timepoints of fiber growth; black arrow, the direction of fiber growth. **B**, The maximum fold changes (ΔF/F) of each fiber as treated without Dox or with 100 ng/ml Dox in corresponding times. Each black dot indicated one maximum ΔF/F of one fiber. Data are shown as the mean ± SD of n = 3 independent experiments, *p* values are calculated using one-way ANOVA.

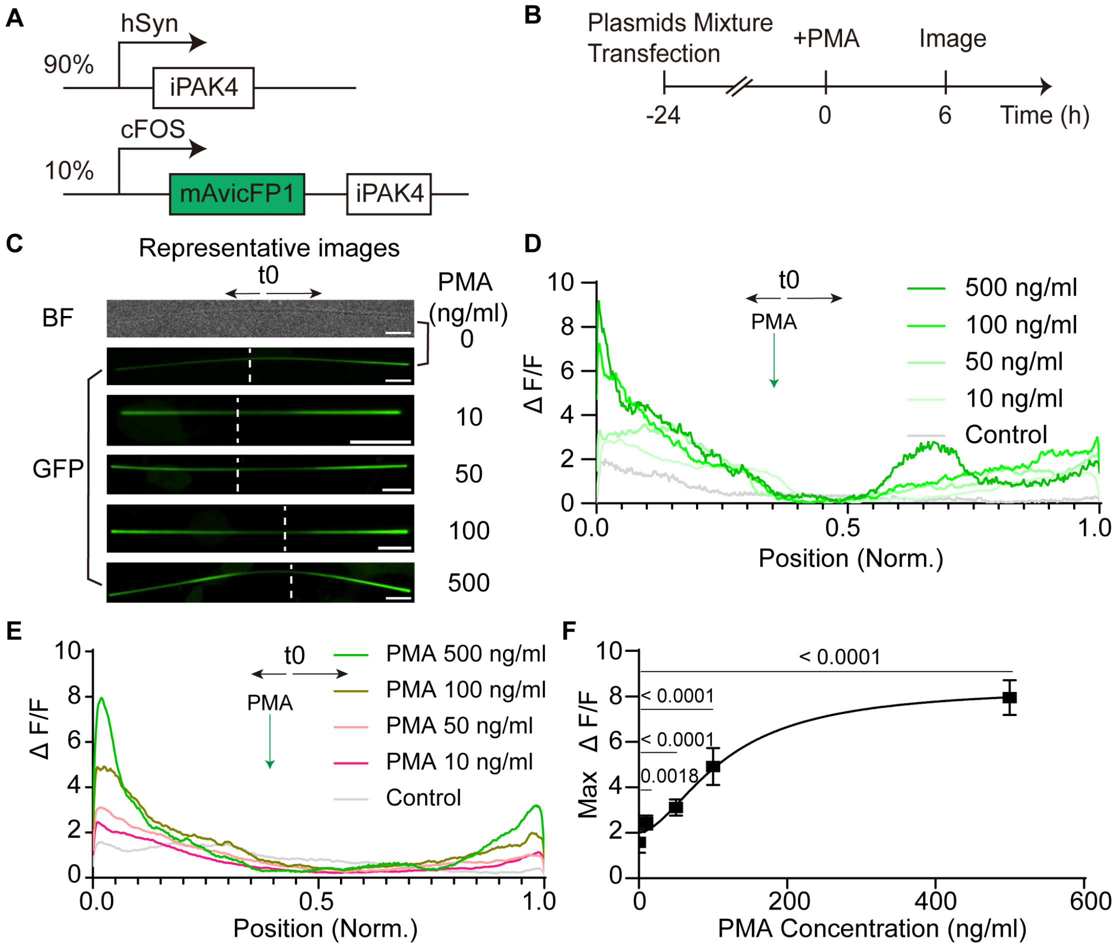


**Supplementary Figure 18.** Dose-dependent activation of endogenous cFos signaling in primary hippocampus neurons

**A**, The expression cassette used for PMA-induced cFos transcriptional activity recordings in mouse primary hippocampus neurons. **B**, Experimental protocol for recording PMA-induced cFos transcription activities. Transcription was activated via adding different concentrations of PMA at 24h post-transfection, and images were captured 6h later. **C**, Representative images of different concentrations of PMA treated as in B. Scale bars, 10 μm. **D**, Corresponding ΔF/F profiles of the fiber shown in C. **E**, Averaged ΔF/F profiles of the fiber as treated in B. Control without PMA, n = 8; 10 ng/ml PMA, n = 12; 50 ng/ml PMA, n = 13; 100 ng/ml PMA, n = 6; 500 ng/ml PMA, n = 6 fibers from 3 independent cultures. **F**, Averaged ΔF/F maxima for each treatment condition from the profiles of E. Data are shown as the mean ± SD of 3 independent experiments. $p = 0.0018$ (10 ng/ml PMA vs control), $p < 0.0001$ (50, 100, 500 ng/ml PMA vs control) using a one-way ANOVA. The curve is interpolated using nonlinear fit built in the GraphPad Prism (sigmoidal, 4PL, X is concentration). t0 and white dash lines, the start timepoints of fiber growth; black arrow, the direction of fiber growth; green arrow, the application of PMA.

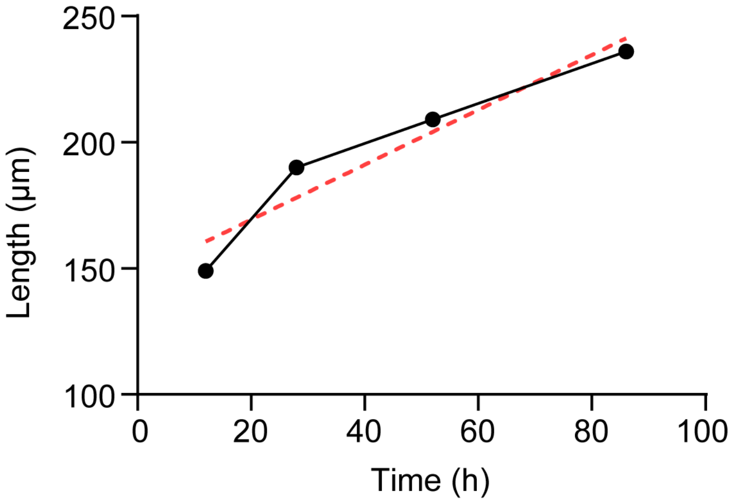


**Supplementary Figure 19**. Single fiber growth in primary neurons. Related to Fig. 2B

This plot displays the growth data for the same fiber featured in Fig. 2B. The black line depicts the actual fiber growth over time (in hours), while the red dashed line illustrates an interpolated linear model.

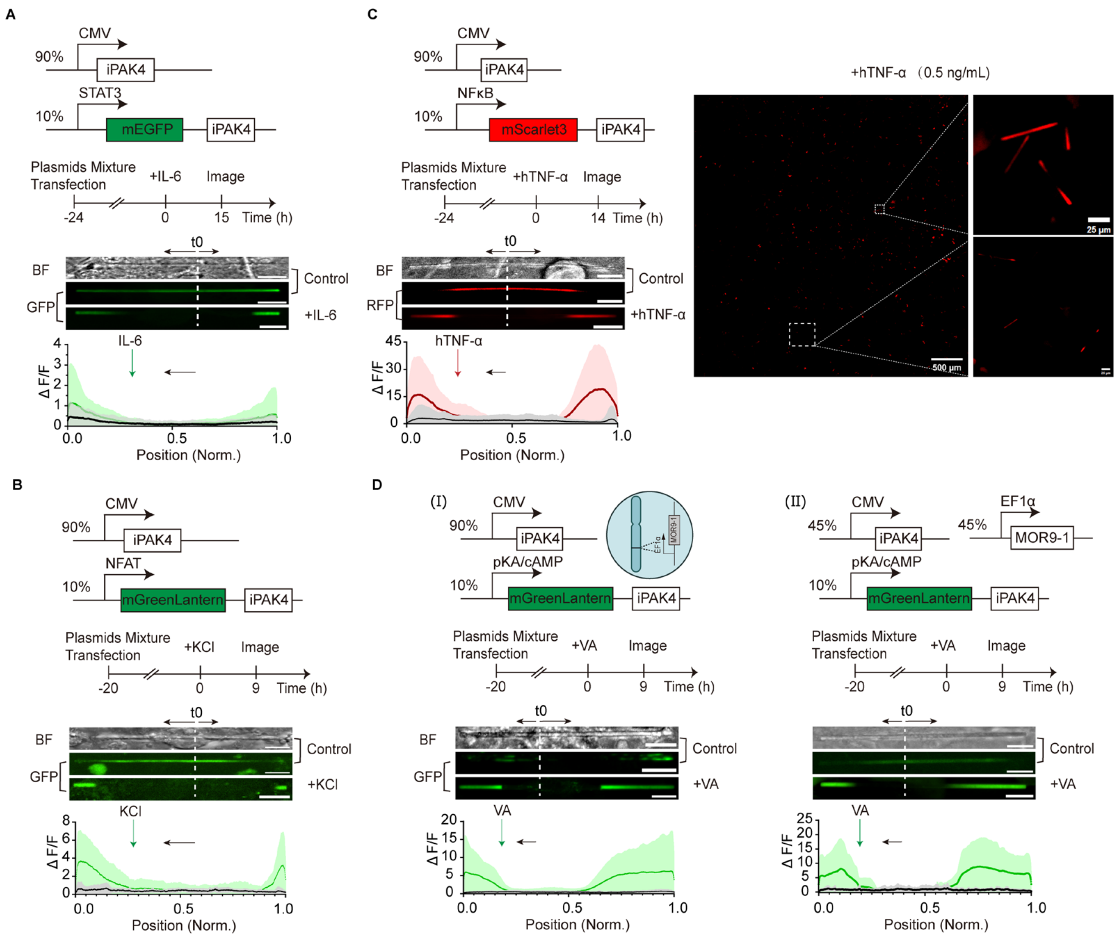


**Supplementary Figure 20.** Engineering of signaling-specific Ticker tapes in HEK293T

**A**, HEK-293T were transfected with a constitutive iPAK4 expression vector (90%, w/w) and 10% (w/w) of a STAT3-inducible mEGFP-tagged iPAK4 expression vector. Microscopic analysis with averaged fluorescence signals (ΔF/F) indicative for activation of STAT3 signaling pathway was performed at 15h after stimulation with 500 ng/ml IL-6. n = 20 in control and n = 32 fibers in IL-6 treated group were shown in ΔF/F plot. **B**, HEK-293T were transfected with a constitutive iPAK4 expression vector (90%, w/w) and 10% (w/w) of a NFAT-inducible mGreenLantern-tagged iPAK4 expression vector. Microscopic analysis with ΔF/F indicative for activation of NFAT signaling pathway was performed at 9h after stimulation with 45 mM KCl. n = 12 in control and n = 26 fibers in KCl treated group were shown in ΔF/F plot. **C**, HEK-293T were transfected with a constitutive iPAK4 expression vector (90%, w/w) and 10% (w/w) of a NFκB-inducible mScarlet3-tagged iPAK4 expression vector. Microscopic analysis with ΔF/F indicative for activation of NFκB signaling pathway was performed at 14h after stimulation with 0.5 ng/ml human derived TNF-α (hTNF-α). A broad view image with 0.5 ng/ml hTNF-α was shown on the right with represented fully activated fibers. n = 21 in control and n = 19 fibers in hTNF-α treated group were shown in ΔF/F plot. **D**, HEK-293T were transfected with a constitutive iPAK4 expression vector and a cAMP-inducible mGreenLantern-tagged iPAK4 expression vector. Microscopic analysis with ΔF/F indicative for activation of pKA/cAMP signaling pathway was performed at 9h after stimulation with 500 μM vanillic acid (VA). For cAMP-dependent systems, a vanillic acid-responsive GPCR coupling to the Gαs-PKA axis (MOR9-1) was stably integrated into HEK-293T (**I**) or co-transfected (**II**) with a designated ratio, as illustrated in the figure. (**I**), Control, n = 7 fibers, VA, n = 19 fibers; (**II**), Control, n = 22, VA, n = 54 fibers, from 3 biological replicates. The maxima ΔF/F in IL-6 treated: 1.14 vs control: 0.46, in KCl treated: 3.68 vs. control: 0.70, in TNF-α treated: 19.2 vs control: 3.07, in VA treated: 6.24 vs control: 0.68 (**I**), 9.02 vs control: 1.57 (**II**). Solid line, mean; shaded area, SD. t0 and white dash lines, the start timepoints of fiber growth; black arrow, the direction of fiber growth; red and green arrow, the application of respective agonist. Scale bars, 10 μm, unless indicated in figures.

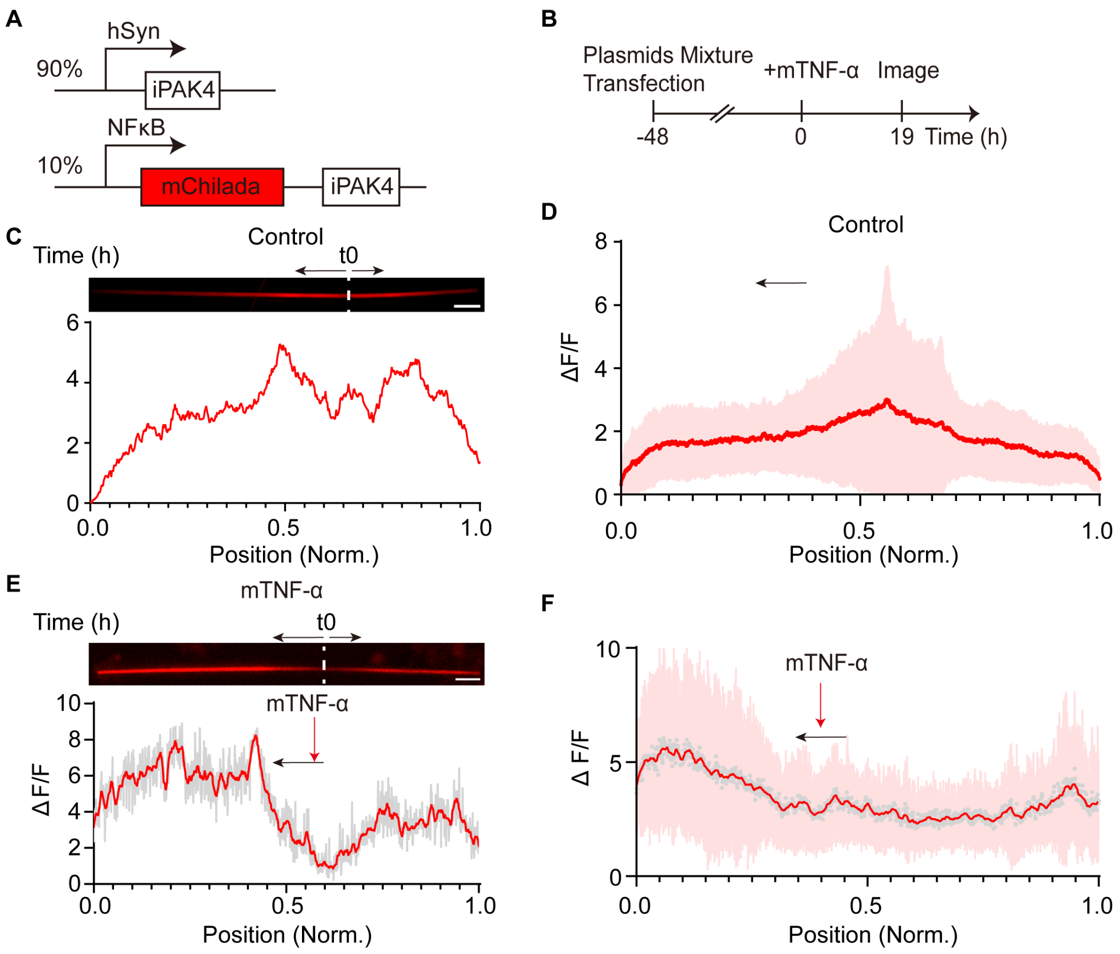


**Supplementary Figure 21**. Validation of NFκB - specific ticker tapes in primary neurons

**A**, Genetic constructs used for expression of fluorescent ticker tapes in cultured murine primary hippocampus neurons. **B**. Experimental protocol for validation of NFκB-specific Ticker tapes in primary neurons. 48 h post-transfection, neurons were stimulated with or without 200 ng/ml mTNF-α before microscopic images were acquired after 19 h. **C**, Representative image and corresponding ΔF/F profiles of the fiber without mTNF-α treatment. Red line, raw profiles. Scale bars, 10 µm. **D**, Averaged ΔF/F profiles of the fibers without mTNF-α treatment. Red line, mean; shaded area, SD. N = 20 fibers from 3 biological replicates. **E**, Representative image and corresponding ΔF/F profiles of the fiber with mTNF-α treatment. Gray area, raw data; Red line, smooth profiles by averaging 10 values on each side, and using a second order smoothing polynomial. Scale bars, 10 µm. **F**, Averaged ΔF/F profiles of the fibers with mTNF-α treatment. Gray dots, raw data of the mean; red line, smooth mean profiles by averaging 10 values on each side, and using a second order smoothing polynomial; shaded area, SD. N = 16 fibers from 3 biological replicates. t0 and white dash lines, the start timepoints of fiber growth; black arrow, the direction of fiber growth; red arrow, the application of mTNF-α.

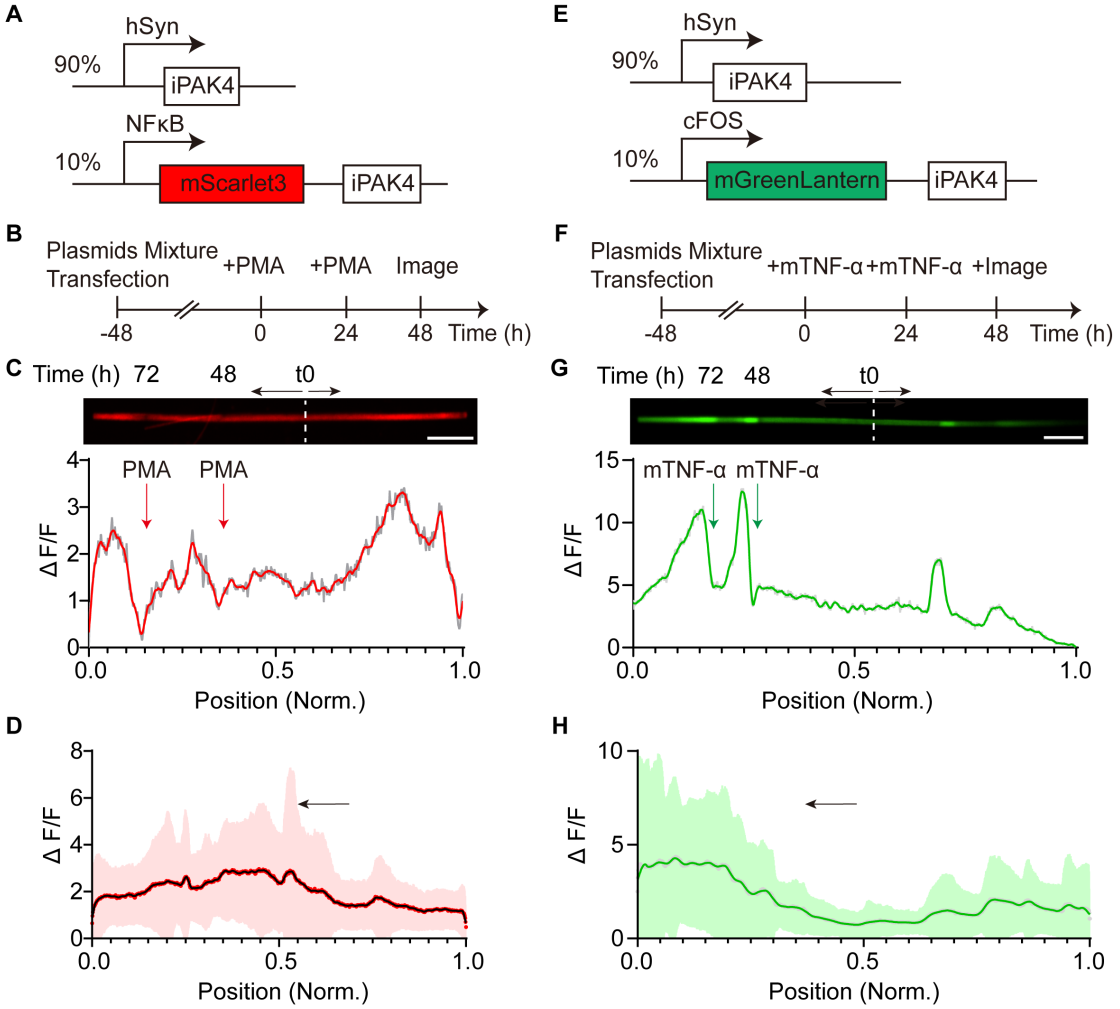


**Supplementary Figure 22.** Potential crosstalk between cFos and NFκB signaling in neurons

**A**, Genetic constructs used for recording NFκB signaling in cultured murine primary hippocampus neurons. **B**. Experimental protocol for recording PMA induced NFκB signaling in primary neurons. 48h post-transfection, neurons were stimulated with 500 ng/ml PMA on two consecutive days before microscopic images were acquired after 48h. **C**, Representative image and corresponding ΔF/F profiles of the fiber treated in B under A constructs. Scale bars, 10 µm. Gray area, raw values; Red line, smooth profiles by averaging 15 values on each side, and using a second order smoothing polynomial. **D**, Averaged ΔF/F profiles of the fibers treated in B. Red dots, raw values of the mean; black line, smooth mean profiles by averaging 10 values on each side, and using a second order smoothing polynomial; shaded area, SD. n = 31 fibers from 3 biological replicates. **E**, Genetic constructs used for recording cFos signaling in cultured murine primary hippocampus neurons. **F**. Experimental protocol for recording mTNF-α induced cFos signaling in primary neurons. 48h post-transfection, neurons were stimulated with 200 ng/ml mTNF-α on two consecutive days before microscopic images were acquired after 48h. **G**, Representative image and corresponding ΔF/F profiles of the fiber treated in F under E constructs. Scale bars, 10 µm. Gray area, raw values; green line, smooth profiles by averaging 10 values on each side using a second order smoothing polynomial. **H**, Averaged ΔF/F profiles of the fibers treated in B. Gray dots, raw values of the mean; green line, smooth mean profiles by averaging 20 values on each side using a second order smoothing polynomial; shaded area, SD. n = 38 fibers from 3 biological replicates. t0 and white dash lines, the start timepoints of fiber growth; black arrow, the direction of fiber growth; red arrow, the application of PMA; green arrow, the application of mTNF-α.

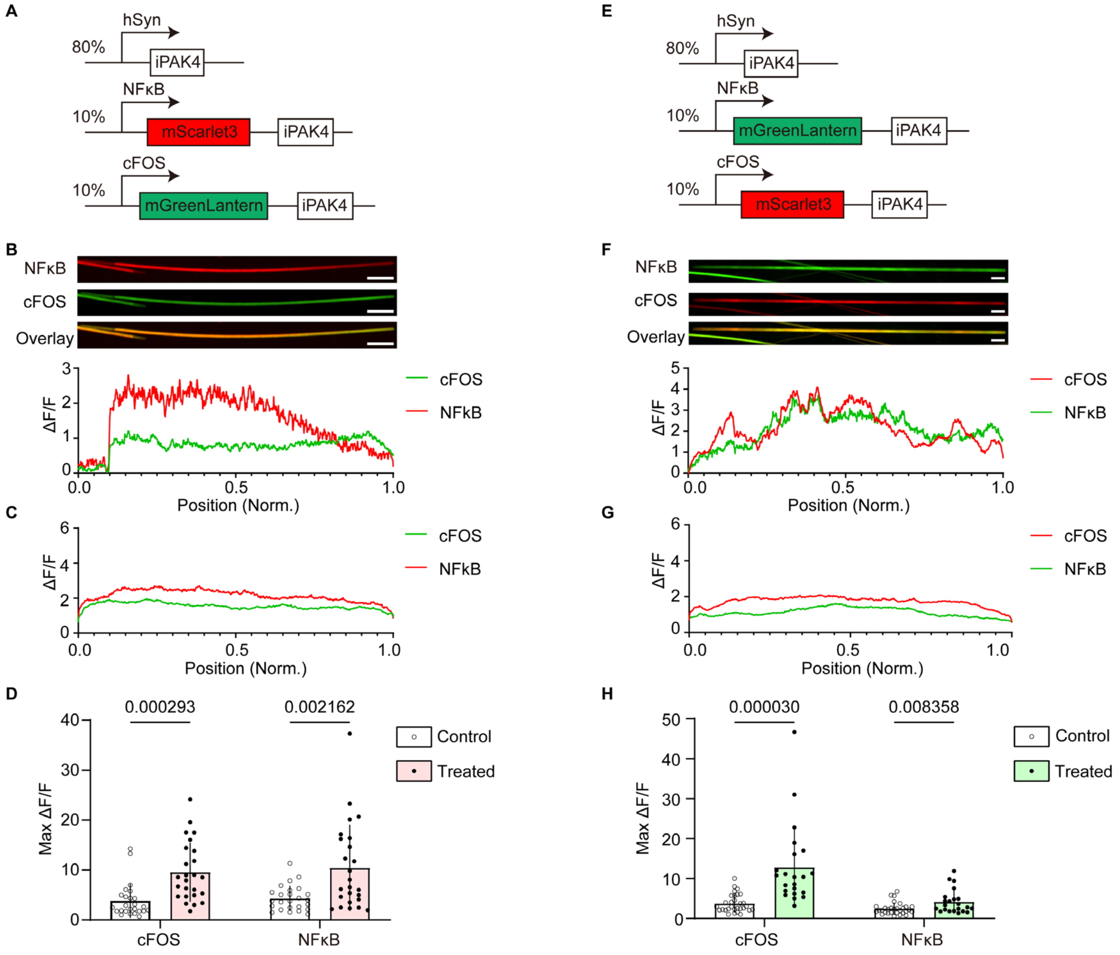


**Supplementary Figure 23**. Control condition in cFos and NFκB multiplexing FPTT. Related to Fig. 3

**A**, The expression cassette used for recording the crosstalk between cFOS and NFκB in mouse primary hippocampus neurons. **B**, Representative image and corresponding ΔF/F profiles of the fiber without PMA and mTNF-α treatment. Green line, cFOS activity; red line, NFκB activity. Scale bars, 10 µm. **C**, Averaged ΔF/F profiles of the fiber without PMA and mTNF-α treatment. Red solid line, mean of NFκB activity; green solid line, mean of cFOS activity. N = 25 fibers from 3 independent cultures. **D**, Statistical analysis of ΔF/F maxima comparing control conditions with PMA and mTNF-α treated conditions. Results are presented as mean±SD from 3 independent experiments. Individual data points are shown as white circles for control and black circles for treated conditions. Statistical significance (*p*-values) was determined via a multiple unpaired *t*-test. **E**, The expression cassette used for recording the crosstalk between cFOS and NFκB in mouse primary hippocampus neurons. **F**, Representative image and corresponding ΔF/F profiles of the fiber without PMA and mTNF-α treatment. Red solid line, cFOS activity; green solid line, NFκB activity. Scale bars, 10 µm. **G**, Averaged ΔF/F profiles of the fiber without PMA and mTNF-α treatment. Red solid line, mean of cFOS activity; green solid line, mean of NFκB activity. N = 30 fibers from 3 independent cultures. **H**, Statistical analysis of ΔF/F maxima comparing control conditions with PMA and mTNF-α treated conditions. Results are presented as mean±SD from 3 independent experiments. Individual data points are shown as white circles for control and black circles for treated conditions. Statistical significance (*p*-values) was determined via a multiple unpaired *t*-test.

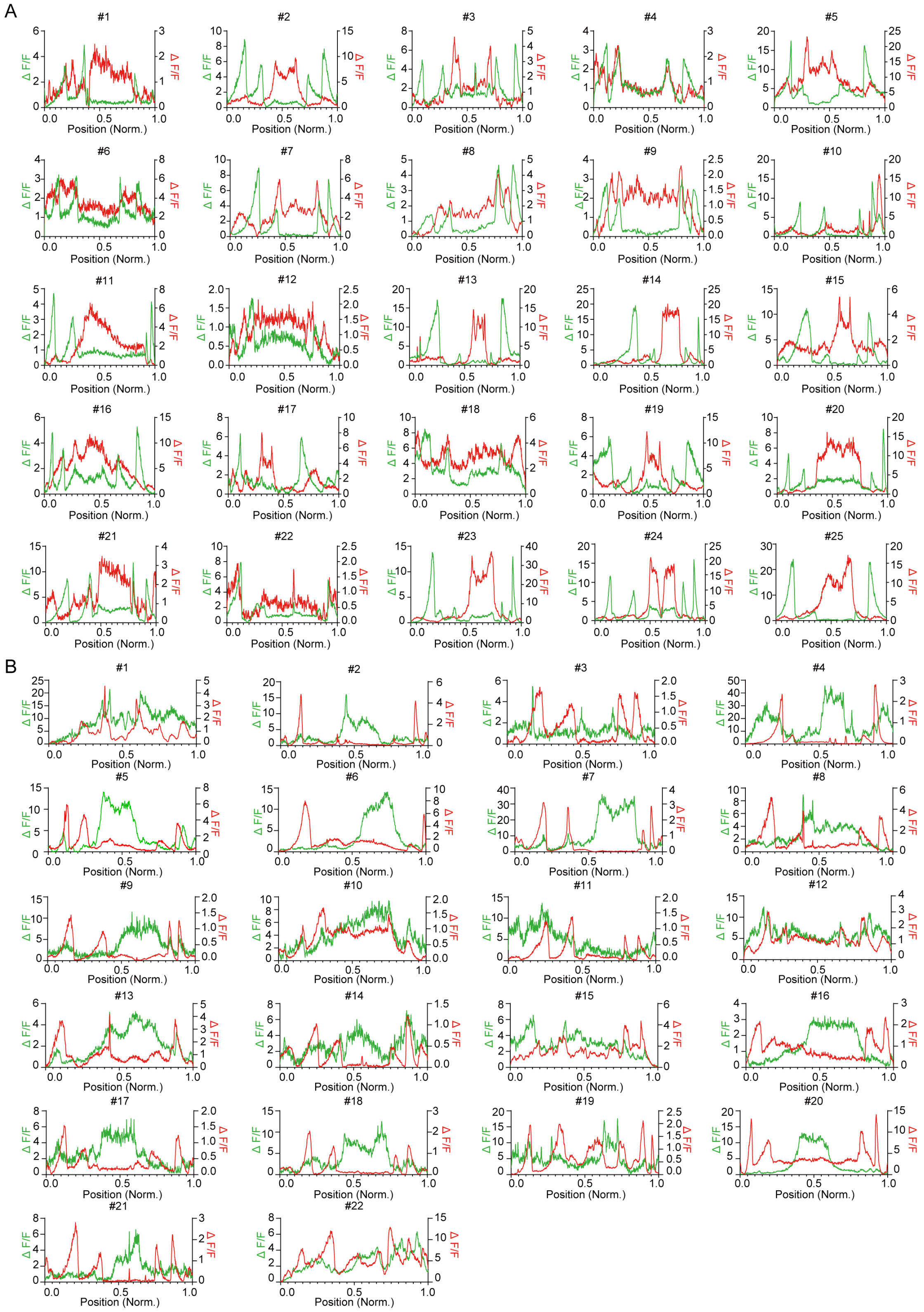


**Supplementary Figure 24.** Temporal recording of multiplexed cFos and NFκB activities in primary neurons

**A**, more representative ΔF/F profiles as shown in Fig. 5A-C; **B**, more representative ΔF/F profiles as shown in Fig. 5D-E.

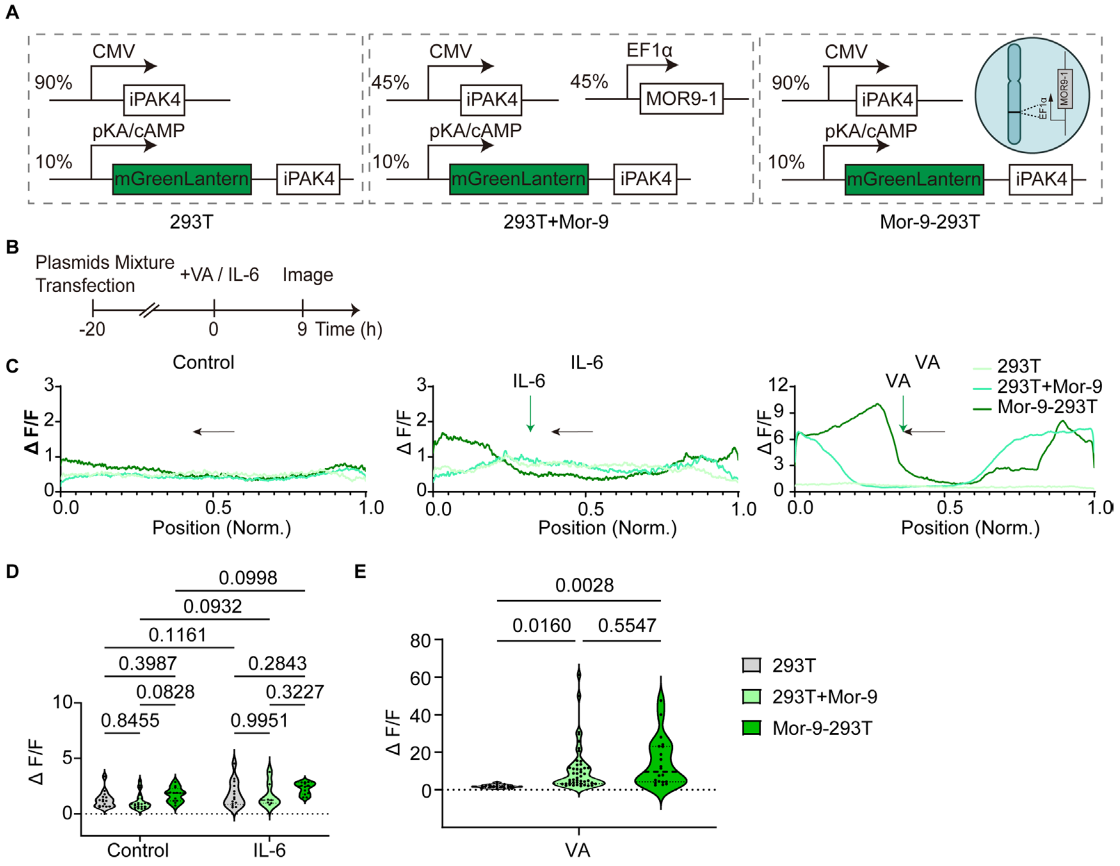


**Supplementary Figure 25**. vanillic acid activated cAMP signaling highly rely on its receptor Mor9-1

**A**, Genetic construct of ticker tape for reporting cAMP activation in HEK293T cells. While a co-transfection with Mor9-1 in 293T was named as 293T+Mor-9, stably integrated Mor9-1 in 293T cells was called Mor-9-293T, used here and throughout the rest of the figures. **B**, Experimental protocol for recording cAMP signaling in A construct. 20h post-transfection, cells were stimulated with either 500 µM vanillic acid (VA), or 500 ng/ml IL-6, or without any stimulation (Control) for 9 hours before microscopic images were acquired. **C**, Averaged ΔF/F profiles in each condition as shown in B. Control in 293T cells, n = 15 fibers, in 293T+Mor-9, n= 22 fibers, in Mor-9-293T, n = 12 fibers; IL-6 treated in 293T cells, n = 14 fibers, in 293T+Mor-9, n= 7 fibers, in Mor-9-293T, n = 12 fibers; VA treated in 293T cells, n = 17 fibers, in 293T+Mor-9, n= 46 fibers, in Mor-9-293T, n = 21 fibers. All fibers are from 3 biological replicates. Black arrow, the direction of fiber growth; green arrow, the application of IL-6 or VA. **D**, Averaged ΔF/F maxima of control and IL-6 treated conditions in 293T, 293T+Mor-9, and Mor-9-293T cells. Data are shown as the mean±SD of 3 independent experiments. *P* values were calculated using a Two-way ANOVA. **E**, Averaged ΔF/F maxima of VA treated conditions in 293T, 293T+Mor-9, and Mor-9-293T cells. Data are shown as the mean ± SD of 3 independent experiments, individual data points are indicated in black circles. *P* values were calculated using a One-way ANOVA.

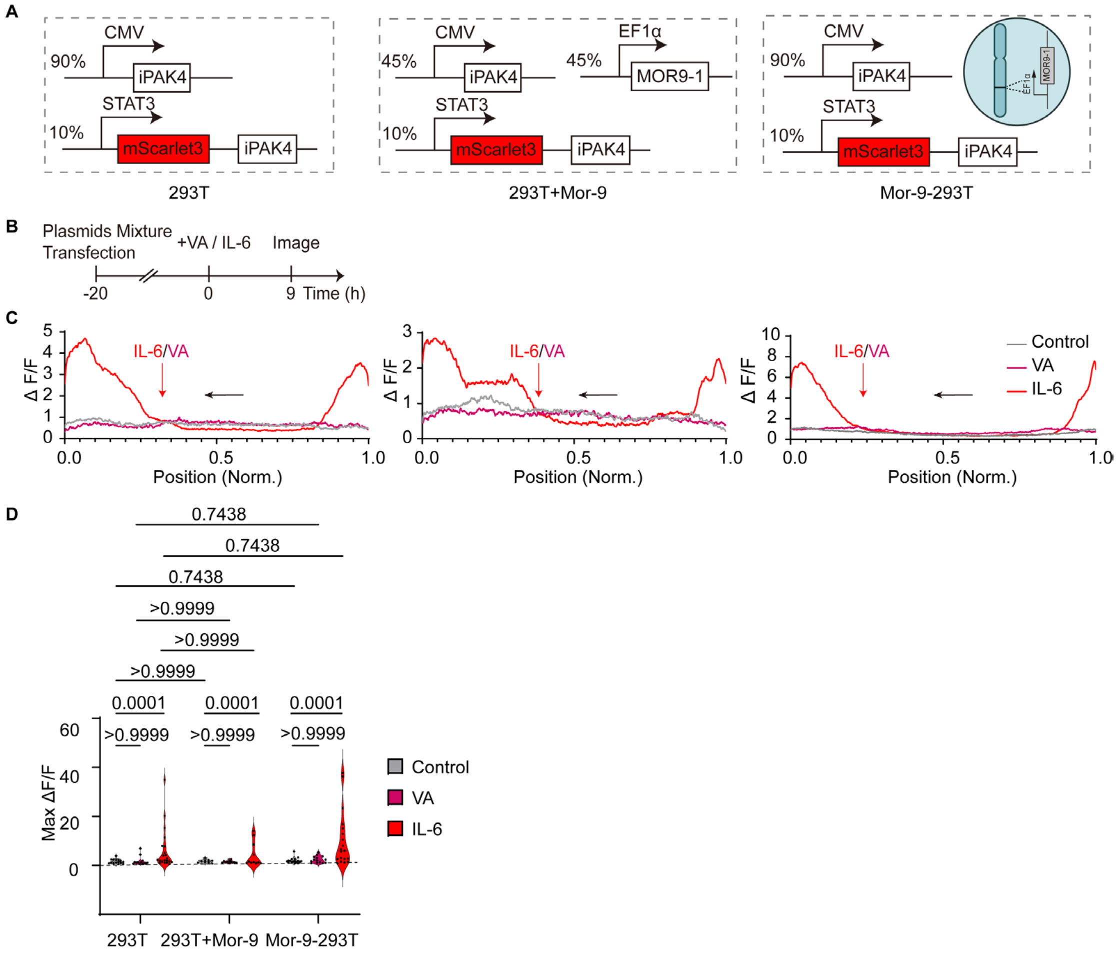


**Supplementary Figure 26**. Validation of vanillic acid effect on STAT3 by Ticker Tape

**A**, Genetic construct of ticker tape for reporting STAT3 activation in HEK293T cells. **B**, Experimental protocol for recording STAT3 signaling in A constructs. 20h post-transfection, cells were stimulated with either 500 µM vanillic acid (VA), or 500 ng/ml IL-6, or without any stimulation (Control) for 9 hours before microscopic images were acquired. **C**, Averaged ΔF/F profiles in each condition as shown in B. In 293T cells, control, n = 15 fibers, VA, n = 13 fibers, IL-6, n = 21 fibers; in 293T+Mor-9, control, n= 9 fibers, VA, n = 11 fibers, IL-6, n = 12 fibers; in Mor-9-293T, Control, n = 17 fibers, VA, n = 14 fibers, IL-6, n = 19 fibers from 3 biological replicates. Gray lines, mean ΔF/F of STAT3 expression fluorescence level in all fibers in control; magenta lines, mean ΔF/F of STAT3 expression fluorescence level in all fibers in VA treated group; red lines, mean ΔF/F of STAT3 expression fluorescence level in all fibers in IL-6 treated group. Black arrow, the direction of fiber growth; red arrow, the application of IL-6 or VA. **D**, Averaged ΔF/F maxima of control, VA and IL-6 treated conditions in 293T, 293T+Mor-9, and Mor-9-293T cells. Data are shown as the mean ± SD of 3 independent experiments, individual data points are indicated in black circles. A two-way ANOVA was performed to determine statistical significance (*p* values).

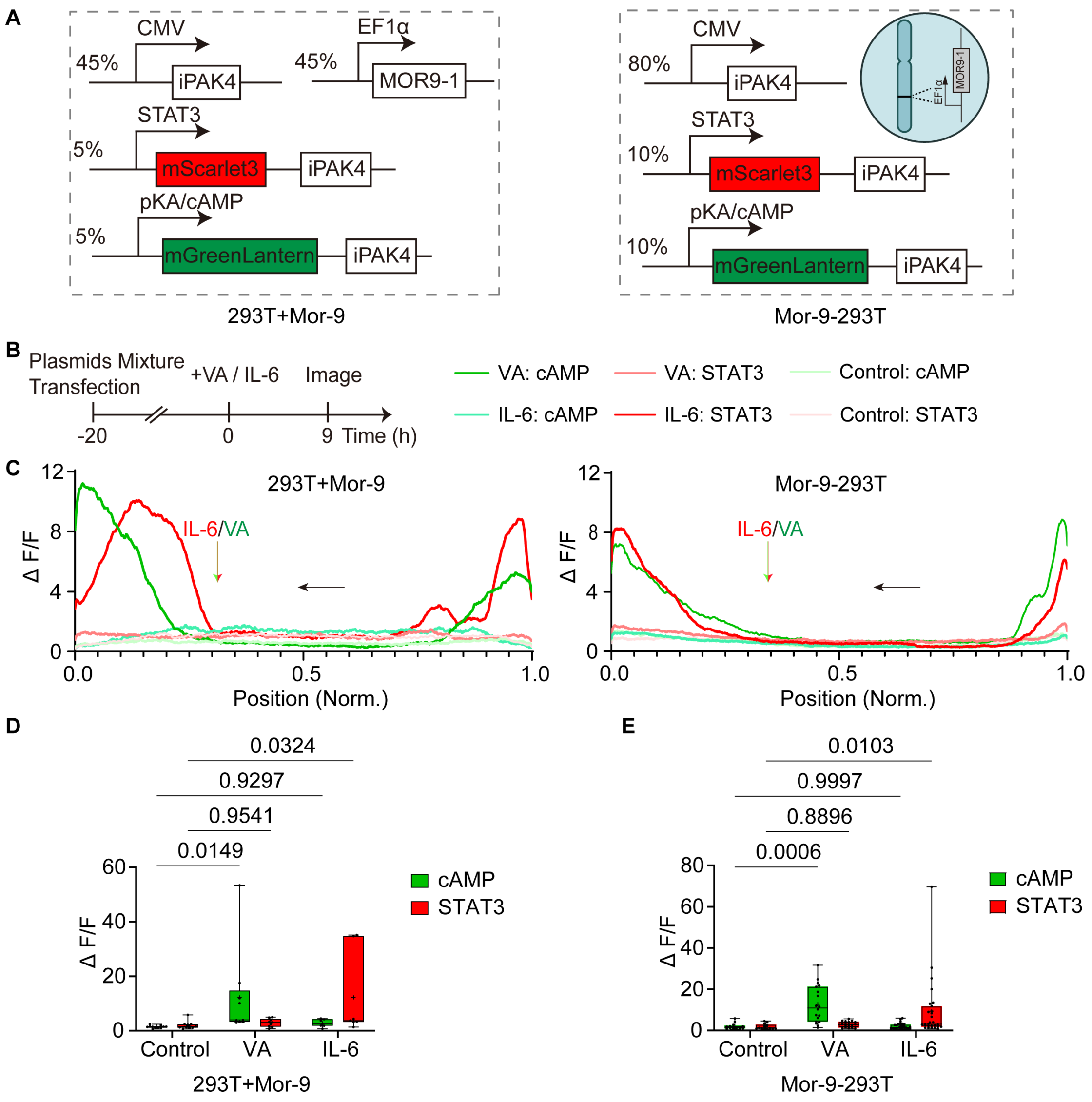


**Supplementary Figure 27**. cAMP and STAT3 are orthogonal in response to extracellular stimuli in HEK293T cells.

**A**, Genetic construct of ticker tape for reporting STAT3 and cAMP activation in HEK293T cells either with a Mor9-1 receptor co-transfection (293T+Mor-9) or in a Mor9-1 stably expressed cell lines (Mor-9-293T). **B**, Experimental protocol for recording STAT3 and cAMP signaling in A construct. 20h post-transfection, cells were stimulated with either 500 μM vanillic acid (VA), or 500 ng/ml IL-6, or without any stimulation (Control) for 9 hours before microscopic images were acquired. **C**, Averaged ΔF/F profiles in each condition of 293T+Mor-9 (Left) and Mor-9-293T cells (Right) were shown. In 293T+Mor-9 cells, control, n = 11 fibers, VA, n = 9 fibers, IL-6, n = 7 fibers; in Mor-9-293T, Control, n = 13 fibers, VA, n = 16 fibers, IL-6, n = 27 fibers from 3 biological replicates. Green lines, mean ΔF/F of cAMP induction level in all fibers at each condition; red lines, mean ΔF/F of STAT3 fluorescence level in all fibers at each condition. Black arrow, the direction of fiber growth; red arrow, the application of IL-6; green arrow, the application of VA. **D**, Averaged ΔF/F maxima of STAT3 and cAMP induction level in control, VA and IL-6 treated conditions in 293T+Mor-9 cells. **E**, Averaged ΔF/F maxima of STAT3 and cAMP induction level in control, VA and IL-6 treated conditions in Mor-9-293T cells. Data are shown as the mean±SD of 3 independent experiments: individual data points are indicated in black circles; error bars indicate the standard error. A Two-way ANOVA was performed to determine statistical significance (*p* values).

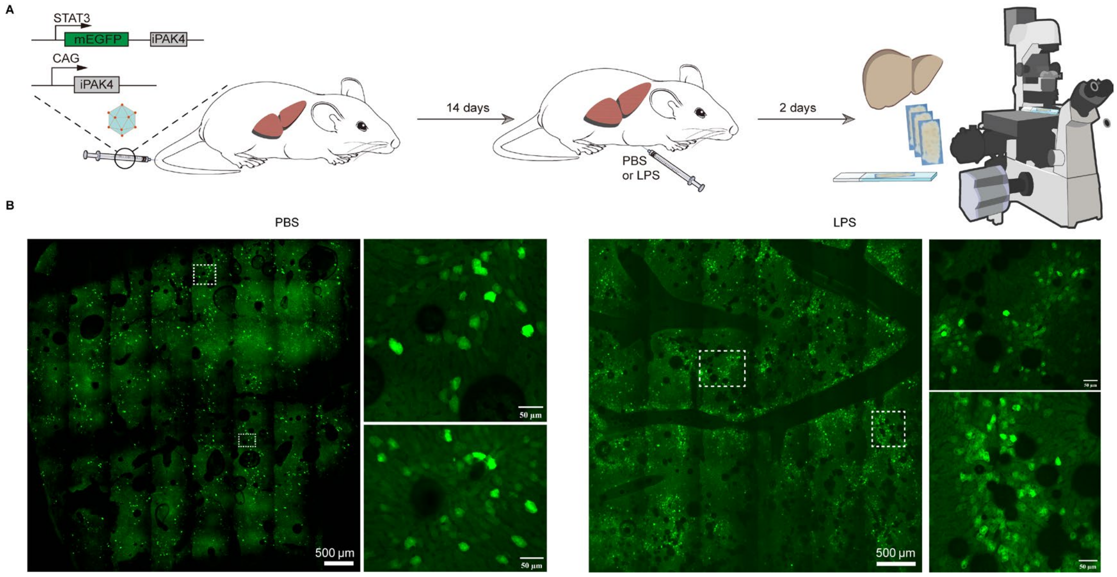


**Supplementary Figure 28**. AAV-mediated delivery of FPTT into mouse liver

**A**, Schematic illustration of two AAV types (CAG-iPAK4, STAT3-mEGFP-iPAK4) delivered to mouse liver via tail vein injection at 3×10^11 GC/mouse. 14 days later, mice received an intraperitoneal injection of PBS or 5 mg/kg LPS. Livers were isolated two days post-injection for microscopy to assess gene expression. **B**, Wide-field image of mouse liver under GFP channel showing green fluorescence only, with no fiber formation observed. Insets show the regions in the dashed boxes. Scale bar was indicated in the figures.

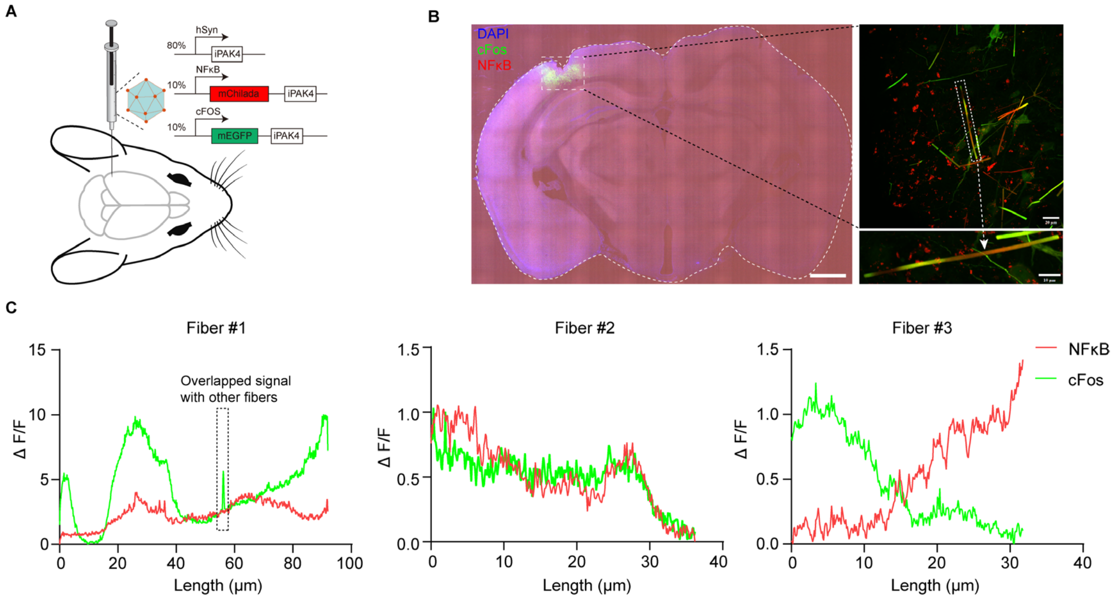


**Supplementary Figure 29.** Stereotaxic injections of FPTT AAVs in the cortex of mouse brain

**A**. Schematic representation of stereotaxic injections into the mouse brain cortex. The adeno-associated viruses (AAVs) used in this study include pSyn-iPAK4, NFκB-mChilada-iPAK4, and cFos-mEGFP-iPAK4, with a titer ratio of 8:1:1. **B**. Representative images displaying a broad view of the mouse brain post AAV injection, along with exemplary fiber formations. Each panel includes a scale bar for reference. **C**. Intensity fold change traces of three fibers indicating cFos and NFκB activity. Note that intersecting dense fibers may result in overlapping intensity profiles, which can complicate the quantification of average profiles.

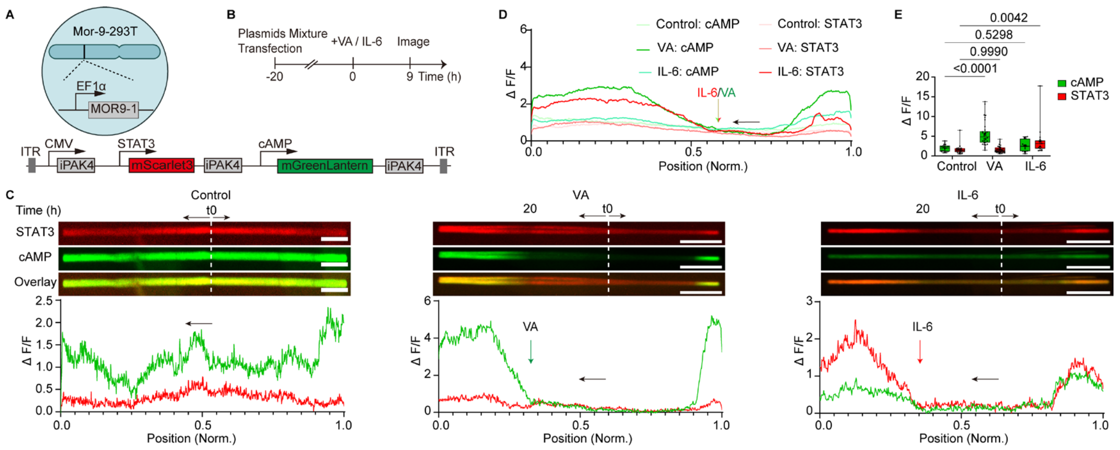


**Supplementary Figure 30.** Multiplexed recording of STAT3 and cAMP signaling by one single plasmid in HEK293T cells

**A**, The expression cassette used for recording multiplexed STAT3 and cAMP signaling in Mor9-1 stably integrated HEK293T cell line and in mouse liver. **B**, Experimental protocol for recording multiplexed STAT3 and cAMP signaling. Transfected Mor9-1 stable cell lines were exposed to stimulation with 500 ng/ml IL-6 or 500 μM vanillic acid (VA) for 9 hours before images were acquired. **C**, Representative image and corresponding ΔF/F profiles of the fiber treated without any stimulation (Control), VA, and IL-6 as indicated in B. Scale bars, 10 μm. t0 and white dash line, the start timepoints of fiber growth; black arrow, the direction of fiber growth; red arrow, the application of IL-6; green arrow, the application of VA. **D**, Averaged ΔF/F profiles of the fiber treated without any stimulation (Control), VA, and IL-6 as indicated in B. N = 22 fibers, 29 fibers, 19 fibers in control, VA, and IL-6 treatments from three biological replicates, respectively. **E**, Averaged ΔF/F maxima for each treatment condition from the profiles of D. Data are shown as the mean±SD of 3 independent experiments. *p* values were calculated using a Two-way ANOVA.

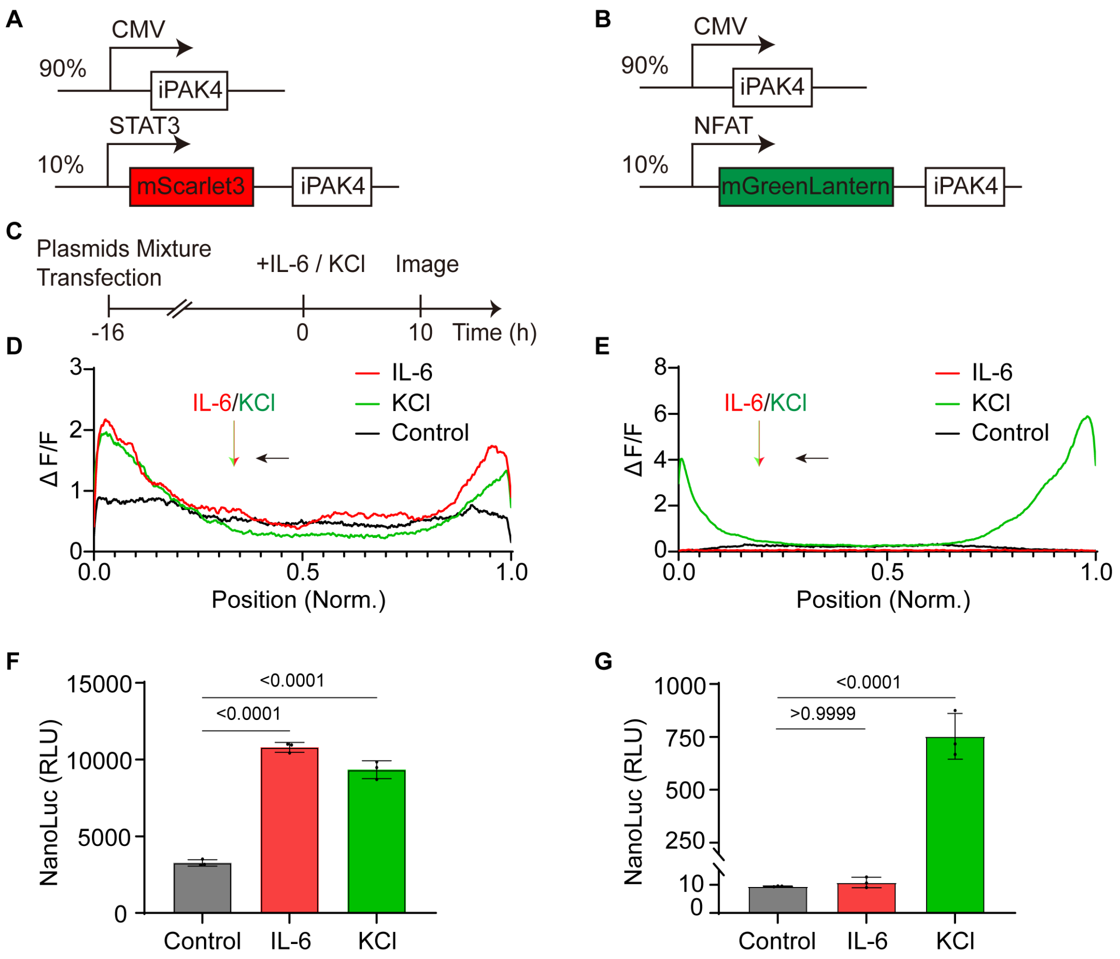


**Supplementary Figure 31**. Potential crosstalk between STAT3 and NFAT signaling validated by Ticker tape and NanoLuc assay in HEK293T cells

**A**, Genetic construct of ticker tape for reporting STAT3 activation in HEK293T cells. **B**, Genetic construct of ticker tape for reporting NFAT activation in HEK293T cells. **C**, Experimental protocol for recording STAT3 and NFAT signaling in both A and B constructs. 16h post-transfection, cells were stimulated with either 45 mM KCl, or 500 ng/ml IL-6, or without any stimulation (Control) for 10 hours before microscopic images were acquired. **D**, Averaged ΔF/F profiles in each condition of STAT3 expression level in A construct. Control, n = 11 fibers, KCl, n = 15 fibers, IL-6, n = 13 fibers from 3 biological replicates. Black lines, mean ΔF/F of STAT3 induction level in all fibers in control group; Green lines, mean ΔF/F of STAT3 induction level in all fibers in KCl treated group; red lines, mean ΔF/F of STAT3 induction level in all fibers in IL-6 treated group. **E**, Averaged ΔF/F profiles in each condition of STAT3 expression level in B construct. Control, n = 4 fibers, KCl, n = 20 fibers, IL-6, n = 3 fibers from 3 biological replicates. Black lines, mean ΔF/F of NFAT induction level in all fibers in control group; Green lines, mean ΔF/F of NFAT induction level in all fibers in KCl treated group; red lines, mean ΔF/F of NFAT induction level in all fibers in IL-6 treated group. Black arrow, the direction of fiber growth; red arrow, the application of IL-6; green arrow, the application of KCl. **F**, Cells were transfected with NanoLuc assays as the same design in A but replacing ticker tape by the NanoLuc luciferase reporter, and treated same protocol as in C. Averaged luminescence signal of STAT3 expression level in each condition. **G**, Cells were transfected with NanoLuc assays as the same design in B but replacing ticker tape by the NanoLuc luciferase reporter, and treated same protocol as in C. Averaged luminescence signal of NFAT expression level in each condition. Data are shown as the mean±SD of 3 independent experiments: individual data points are indicated in black circles; error bars indicate the standard error. A one-way ANOVA was performed to determine statistical significance (*p* values).

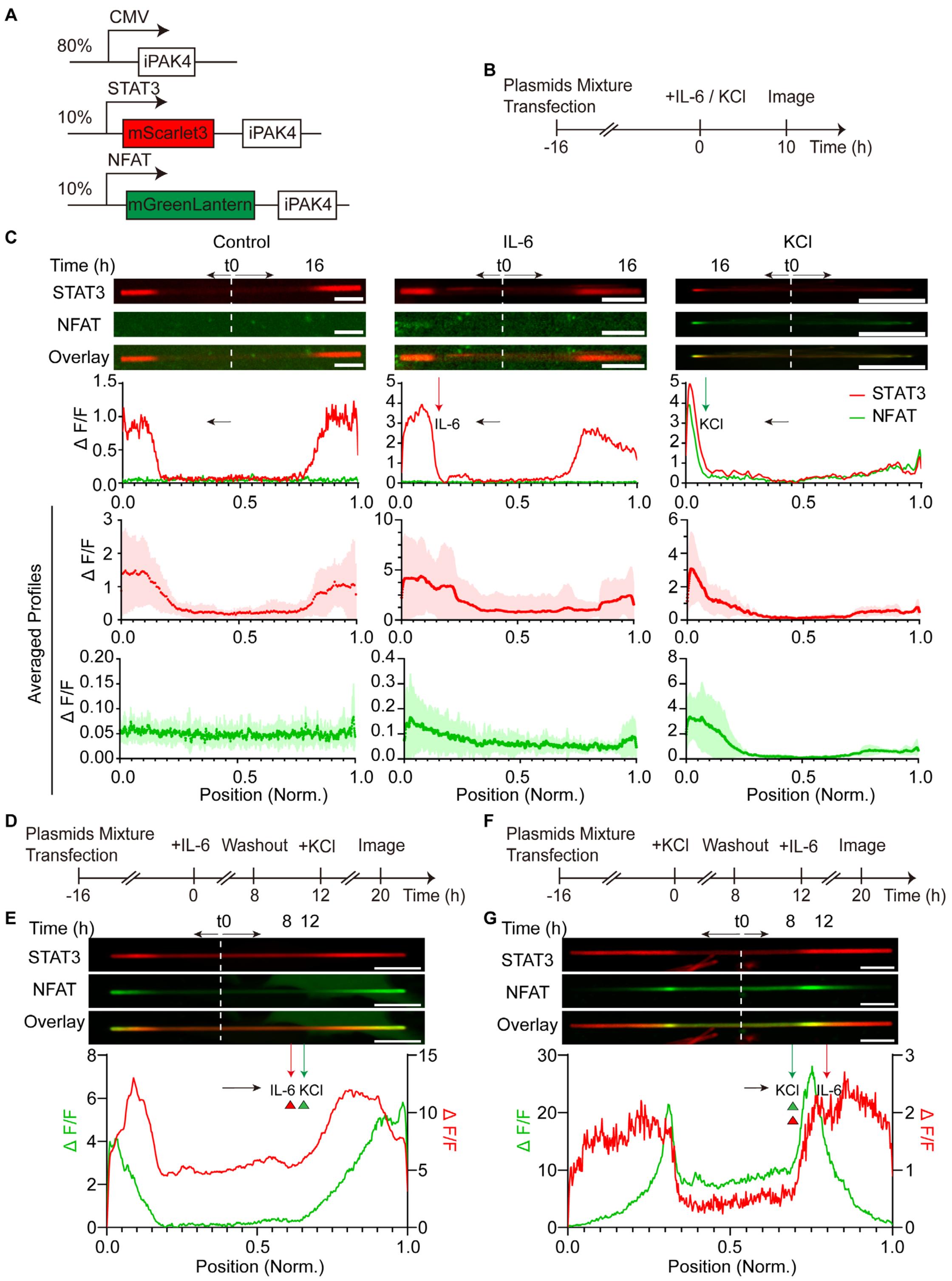


**Supplementary Figure 32.** Interdependent STAT3 and NFAT signaling by multiplexed Ticker tape recording

**A**, The expression cassette used for recording multiplexed STAT3 and NFAT signaling in HEK293T cell. **B**, Experimental protocol for recording multiplexed STAT3 and NFAT signaling. Transfected HEK293T were exposed to stimulation with 500 ng/ml IL-6 or 45 mM KCl for 10 hours before images were acquired. **C**, Representative image, corresponding and averaged ΔF/F profiles of the fiber treated without any stimulation (Control), IL-6, and KCl as indicated in B. N = 10 fibers, 9 fibers, 4 fibers in control, IL-6, and KCl treatments from two biological replicates, respectively. Scale bars, 10 μm. **D**, Experimental protocol for recording multiplexed STAT3 and NFAT signaling. Images were acquired after cells were first stimulated with IL-6 (500 ng/ml, 8h) followed by drug washout (4h) and switching to stimulation with KCl (45mM, 8h). **E**, Representative images and corresponding ΔF/F profiles of the fiber treated in D. t0 and white dash line, the start timepoints of fiber growth; black arrow, the direction of fiber growth; red arrow, the application of IL-6; green arrow, the application of KCl. Green triangle and red tangle indicate the activation of STAT3 and NFAT activity, respectively. Scale bars, 10 μm. **F**, Experimental protocol for recording multiplexed STAT3 and NFAT signaling. Images were acquired after cells were first stimulated with KCl (45mM, 8h) followed by drug washout (4h) and switching to stimulation with IL-6 (500 ng/ml, 8h). **G**, Representative images and corresponding ΔF/F profiles of the fiber treated in F. t0 and white dash line, the start timepoints of fiber growth; black arrow, the direction of fiber growth; red arrow, the application of IL-6; green arrow, the application of KCl. Green triangle and red tangle in same line indicate both STAT3 and NFAT activity were activated. Scale bars, 10 μm.

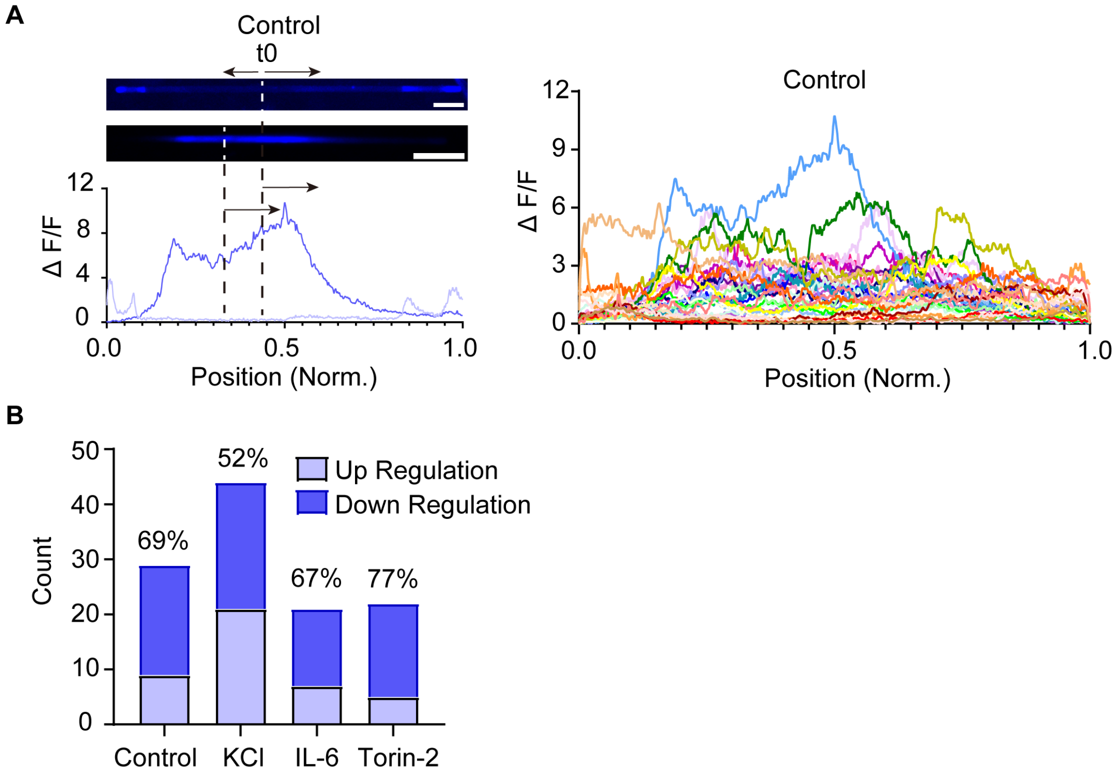


**Supplementary Figure 33.** mTOR driven FPTT in control condition. Related to Figure 5B.

**A**, Various ΔF/F profiles of the mTOR driven fiber in control condition without any stimulation. Representative image in "up regulation" and "down regulation" form and corresponding ΔF/F profiles were show, same as described in Figure 5B. t0 and dash line, the start timepoint of each fiber growth; black arrow, the direction of fiber growth. **B**, Percentage of fibers in "up regulation" and "down regulation" form in control, KCl, IL-6, and Torin-2 treated conditions.

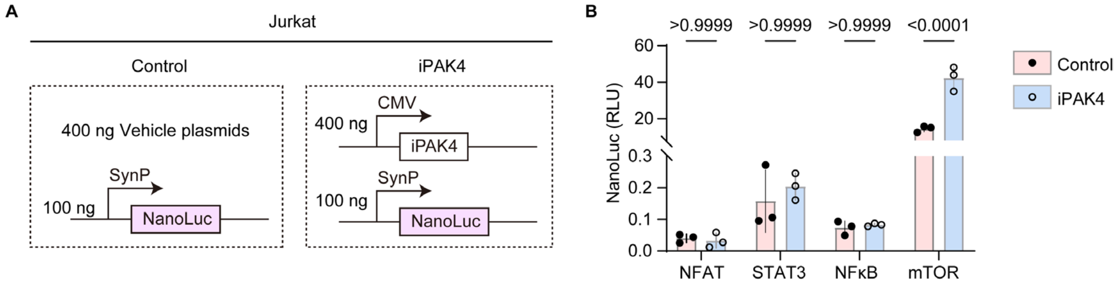


**Supplementary Figure 34.** Effect on specific signaling with iPAK4 expression in Jurkat cells

**A**, Schematic representation of the expression cassettes used to assess the effect of iPAK4 expression on the NFAT, STAT3, NFκB, and mTOR promoters. **B**, Activation of specific signaling pathways by their respective promoters, as measured using a NanoLuc assay. Data are presented as mean±SD from three independent biological replicates. Statistical significance (*p* values) was determined using two-way ANOVA.

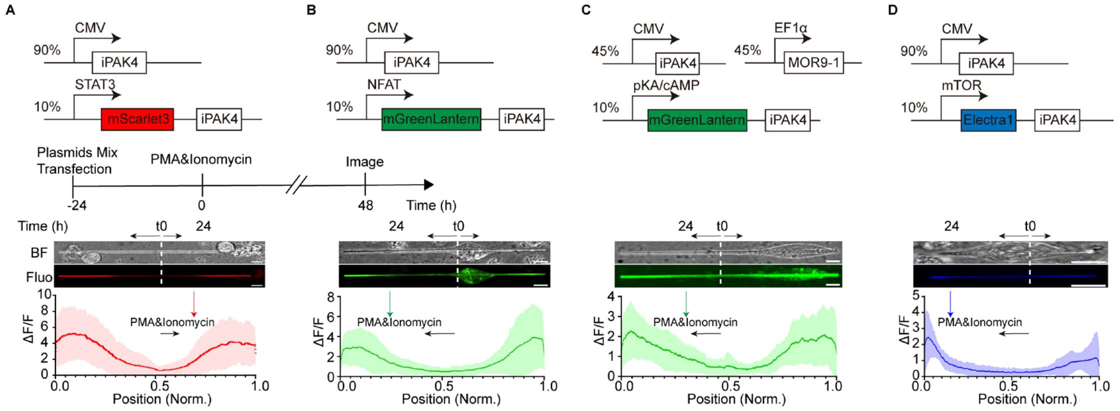


**Supplementary Figure 35.** Recording single physiological activity on ticker tape in Jurkat cells

**A**, Jurkat cells were transfected with a constitutive iPAK4 expression vector (90%, w/w) and 10% (w/w) of a STAT3-inducible expression vector for mScarlet3-tagged iPAK4. **B**, Jurkat cells were transfected with a constitutive iPAK4 expression vector (90%, w/w) and 10% (w/w) of a NFAT-inducible expression vector for mGreenLantern-tagged iPAK4. **C**, Jurkat cells were transfected with a constitutive iPAK4 expression vector (90%, w/w) and 10% (w/w) of a cAMP-inducible expression vector for mGreenLantern-tagged iPAK4. **D**, Jurkat cells were transfected with a constitutive iPAK4 expression vector (90%, w/w) and 10% (w/w) of a mTOR-inducible expression vector for Electra1-tagged iPAK4. For cAMP-dependent systems, a vanillic acid-responsive GPCR coupling to the Gαs-PKA axis (MOR9-1) was co-transfected. Microscopic analysis with averaged fluorescence signals (ΔF/F) indicative for activation of each signaling pathway was performed at 48h after stimulation with PMA (50 ng/mL) and ionomycin (1 µg/mL). Representative images and averaged ΔF/F plots for n > 10 fibers are shown (A, n = 21 fibers; B, n = 20 fibers; C, n = 14 fibers; D, n = 13 fibers, from 3 biological replicates). Solid line, mean; shaded area, SD. t0 and white dash line, the start timepoints of fiber growth; black arrow, the direction of fiber growth; red, green and blue arrow, the application of PMA and ionomycin. Scale bars, 10 µm.

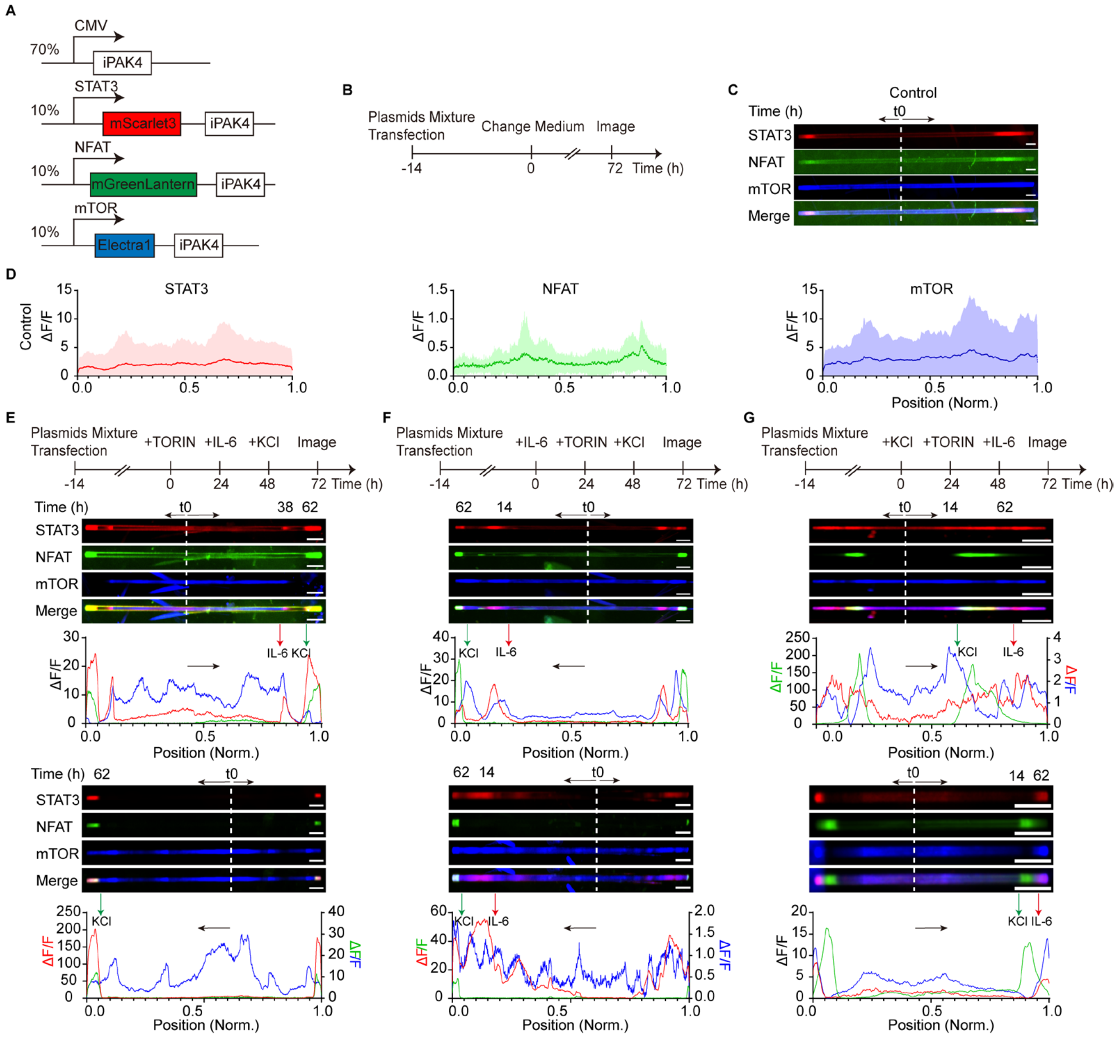


**Supplementary Figure 36.** Multiplexed recordings of STAT3, NFAT, mTOR activities in HEK293T cells

**A**, The expression cassettes used for STAT3, NFAT, and mTOR physiological activities recordings in HEK293T cells. **B**, Experimental protocol for recording physiological activities. For control in c, only medium was changed 14 hours after transfection. **C**, Representative images of control fibers without any treatment in each chanel. Scale bars, 10 µm. **D**, Averaged ΔF/F profiles of control fibers in each channel. Red, STAT3; Green, NFAT; Blue, mTOR (n = 12 fibers from three independent cultures). Solid line, mean; shaded area, SD. **E**, Experimental protocol for recording multiplexed STAT3, NFAT, and mTOR physiological activities. Torin-2 was added in the medium firstly to inhibited mTOR activity14 hours post-transfection, then IL-6 was added 24 hours later to induce STAT3 activity, and the last KCl was added to the medium for activating NFAT activity for another 24 hours. Two representative fibers and corresponding ΔF/F profiles in different cells for reporting STAT3, NFAT and mTOR activities was shown. Scale bars, 10 µm. **F**, Experimental protocol for recording multiplexed STAT3, NFAT, and mTOR physiological activities. IL-6 was firstly added 14 hours post-transfection for 24 hours to induce STAT3 activity, then Torin-2 was added in the medium to inhibited mTOR for another 24 hours and the last KCl was added for 24 hours to activate NFAT activity. Two representative fibers and corresponding ΔF/F profiles in different cells for reporting STAT3, NFAT and mTOR activities in this treatment order was shown. Scale bars, 10 µm. **G**, Experimental protocol for recording multiplexed

STAT3, NFAT, and mTOR physiological activities. KCl was firstly added 14 hours post-transfection for 24 hours to induce NFAT activity, then Torin-2 was added in the medium to inhibited mTOR for another 24 hours and the last IL-6 was added for 24 hours to activate STAT3 activity. Two representative fibers and corresponding ΔF/F profiles in different cells for reporting STAT3, NFAT and mTOR activities in this treatment order was shown. t0 and white dash line, the start timepoints of fiber growth; black arrow, the direction of fiber growth; red arrow, the application of IL-6; green arrow, the application of KCl. Scale bars, 10 μm.

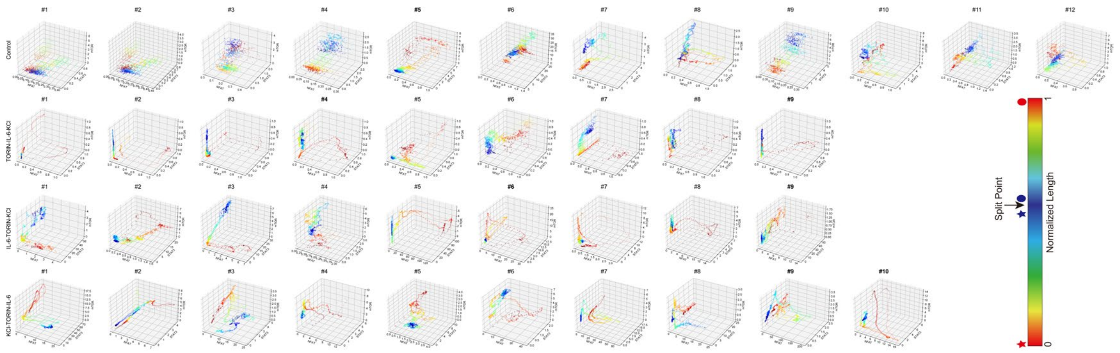

**Supplementary Figure 37.** 3D plots of multiplexed physiological activities in Fig. S20.

3D representation of multiplexed recording physiological activities in Control (Fig. S20 B-D), Torin2-IL-6-KCl (Fig. S20 E), IL-6-Torin2-KCl (Fig. S20 F), KCl-Torin2-IL-6 (Fig. S20 G) condition as illustrated in supplementary figure 20. X axis, NFAT; y axis, mTOR; z axis, STAT3. A split point, which has a highest Pearson correlation on both ends was described as the origin (0), the pentagram represents the traces on the fast-growing end and the circle represents the traces on the slow-growing end, with a dark blue color indicate closer to the split point and bright red color indicate closer the end.

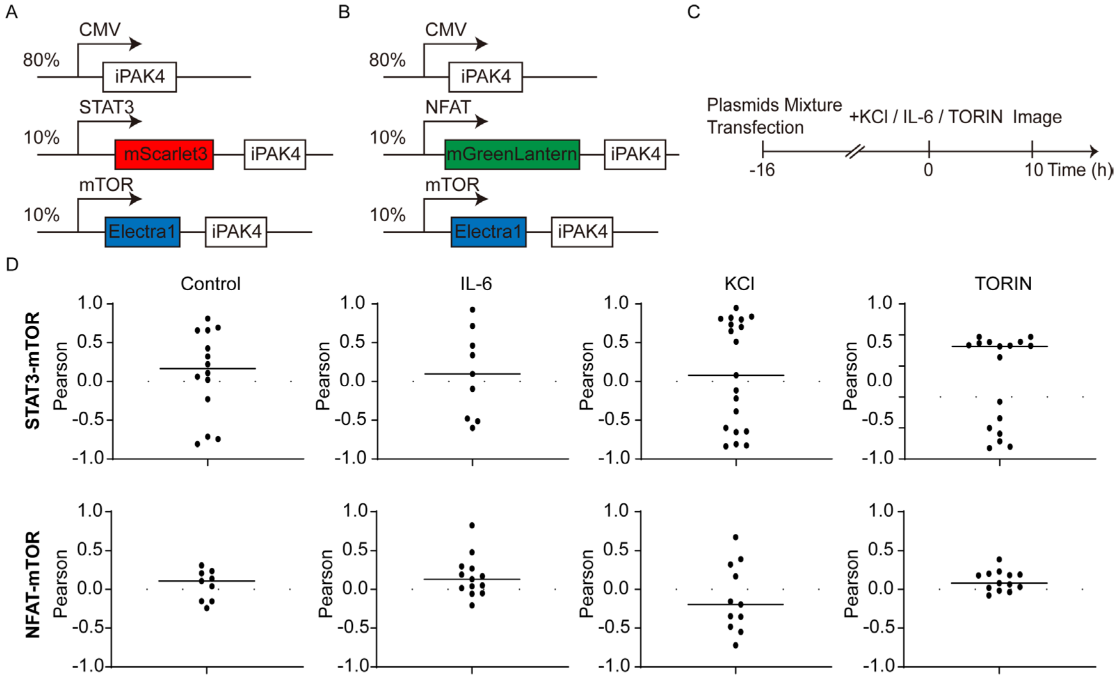


**Supplementary Figure 38**. Pearson correlations of STAT3 and NFAT with mTOR in HEK293T cells

**A-B**, The expression cassette used for recording multiplexed STAT3 (A) and NFAT (B) with mTOR signaling in HEK293T cell. **C**, Experimental protocol for recording multiplexed STAT3 (A) and NFAT (B) with mTOR signaling. Transfected HEK293T were exposed to stimulation with 500 ng/ml IL-6, or 45 mM KCl, or 250 nM Torin-2 for 10 hours before images were acquired. **D**, Pearson coefficient between STAT3 with mTOR (Top), and NFAT with mTOR (Bottom) in control, IL-6, KCl, and Torin-2 treated group. Each dot represents the number of Pearson coefficient in each fiber, solid line represent mean. For STAT3-mTOR correlations, n = 14 fibers in control, n = 9 fibers in IL-6 treated, n = 19 fibers in KCl treated, and n = 17 fibers in Torin-2 treated. For NFAT-mTOR correlations, n = 9 fibers in control, n = 13 fibers in IL-6 treated, n = 11 fibers in KCl treated, and n = 13 fibers in Torin-2 treated. Data was acquired from two biological replicates.

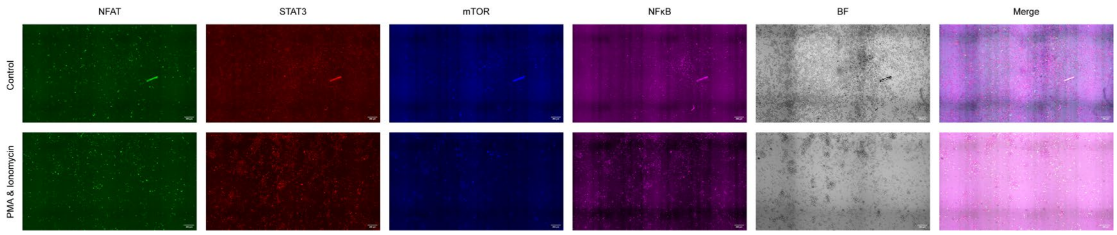


**Supplementary Figure 39**. Wide-field view of Jurkat$_{4s}$ in control and treated conditions. Related to Figure 6.

Jurkat$_{4s}$ in control (top) and treated with PMA & Ionomycin (bottom). Green, NFAT-driven mGreenlantern-iPAK4 expression; red, STAT3-driven mScarlet3-iPAK4 expression; blue, mTOR-driven Electra1-iPAK4 expression. Magenta, NFκB-driven emiRFP670-iPAK4 expression. BF, bright field. Scale bar, 200 µm. No fiber was found without PMA and ionomycin stimulation.
